# Bimodality of gene expression in cancer patient tumors as interpretable biomarkers for drug sensitivity

**DOI:** 10.1101/2020.09.08.288688

**Authors:** Wail Ba-Alawi, Sisira Kadambat Nair, Bo Li, Anthony Mammoliti, Petr Smirnov, Arvind Singh Mer, Linda Penn, Benjamin Haibe-Kains

## Abstract

Identifying biomarkers predictive of cancer cells’ response to drug treatment constitutes one of the main challenges in precision oncology. Recent large-scale cancer pharmacogenomic studies have boosted the research for finding predictive biomarkers by profiling thousands of human cancer cell lines at the molecular level and screening them with hundreds of approved drugs and experimental chemical compounds. Many studies have leveraged these data to build predictive models of response using various statistical and machine learning methods. However, a common challenge in these methods is the lack of interpretability as to how they make the predictions and which features were the most associated with response, hindering the clinical translation of these models. To alleviate this issue, we develop a new machine learning pipeline based on the recent LOBICO approach that explores the space of bimodally expressed genes in multiple large *in vitro* pharmacogenomic studies and builds multivariate, nonlinear, yet interpretable logic-based models predictive of drug response. Using our method, we used a compendium of three of the largest pharmacogenomic data sets to build robust and interpretable models for 101 drugs that span 17 drug classes with high validation rate in independent datasets.

## INTRODUCTION

Identifying reliable predictive biomarkers of drug response is a key step in the era of personalized medicine. Large scale cancer pharmacogenomic studies have boosted the research for finding predictive biomarkers by profiling thousands of human cancer cell lines at the molecular level and screening them with hundreds of drugs^1–5^. Genomic features have been so far regarded as the state-of-the-art method for predicting patients’ response to drugs in the clinic. However, it has been shown that most genomic biomarkers are found in small proportions of patients and within that subset, only a few have shown response to associated drugs^6^.

Several studies have investigated alternative sources for predictive biomarkers of drug sensitivity in cancer pharmacogenomics^7,8^. These studies have shown that gene expression outperforms other molecular features such as mutations and copy-number-variations (CNVs) in predicting drug response in human cancer cell lines^7,8^. Yet, a major criticism of gene expression as a source of predictive biomarkers is the lack of reproducibility due to dependency on profiling assays and batch effects. To overcome such limitations, several studies have focused their analyses on genes that have shown bimodal distribution of expression^9–11^. An advantage of a bimodal gene as a biomarker is that its modes can be used to robustly classify samples into two distinct expression states, allowing for easier interpretation and translation of the biomarker into the clinic. For example, estrogen receptor (ESR1) bimodal expression defines two biological states within breast cancer patients. These states have been used to stratify breast cancer patients into the clinically-relevant subtypes (ER +/-) and derive treatment decisions. Another example in cancer genomics is the use of 73 bimodal genes within ovarian cancer to define molecular subtypes with distinct survival rate^12^. We also have shown that epithelial-to-mesenchymal transition (EMT) related genes were found to be bimodal pan-cancer and predictive of response to statin class of drugs^13^.

Most pharmacogenomic studies that tackled the challenge of finding reliable predictive biomarkers for drug sensitivity employed univariate models for simplicity and interpretability^1–3,14,15^. However, such models do not account for dependencies between genes yielding suboptimal model predictions. Recent studies have applied more sophisticated machine learning techniques that capture dependencies between genes and produce more accurate biomarkers predictive of drug sensitivity^7,16–18^. However, it becomes hard to biologically interpret these predicted biomarkers due to the complexity of these models and how they define the dependencies between the genes. In this study, we developed a machine learning pipeline to explore the large space of bimodally expressed genes and build multivariate, nonlinear, yet interpretable logic-based models predictive of drug response in large *in vitro* pharmacogenomic studies (Fig 1A). Following our proposed approach, we developed robust and interpretable models predictive of drug sensitivity in a large set of more than 500 drugs that were validated and yielded high predictive rates (92% and 61% respectively) in two independent large test sets.

**Fig 1:**
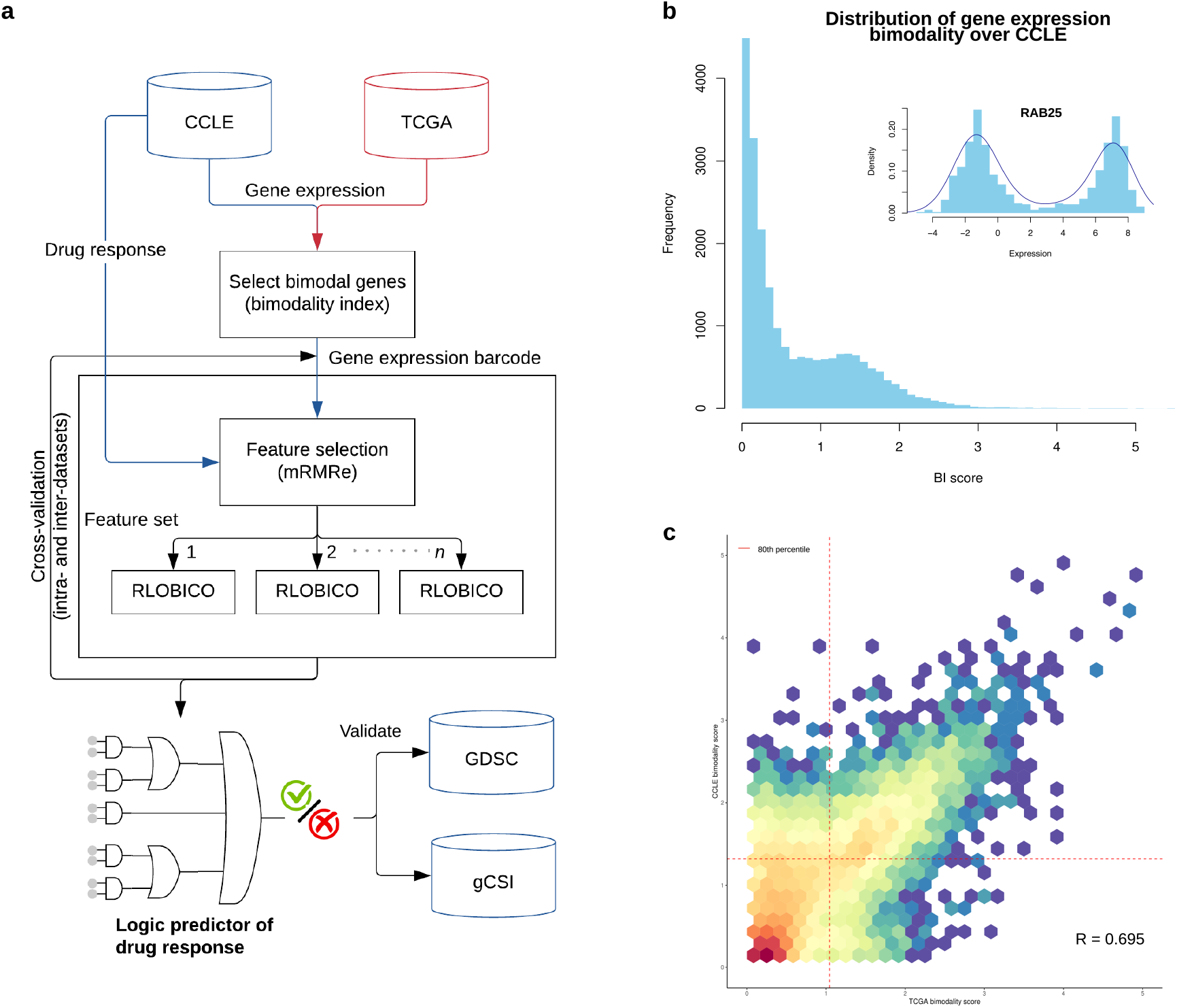
**(a)** An overview of the pipeline to create logic predictors of drug response. **(b)** Distribution of bimodality index scores (BIs) for all genes based on RNAseq gene expression profiles of cell lines in CCLE. **(c)** Distribution of BI scores across CCLE and TCGA. Genes showing high bimodality (>80thpercentile) in both data sets are chosen as global bimodal genes

## RESULTS

### Bimodality of gene expression

To comprehensively explore the space of bimodal gene expression, we performed a genome-wide characterization of gene expression distribution in large sets of patient tumors and immortalized cancer cell lines. Utilizing the gene expression data from the Cancer Cell Line Encyclopedia (CCLE; 945 cell lines from 23 tissue types)^15,19–21^, we determined the expression bimodality of a given gene by fitting a mixture of two Gaussian distributions across all samples and then calculating the bimodality index ^9^ (Fig 1B). We restricted this analysis to solid tumors as hematopoietic and lymphoid cell lines have distinctive molecular profiles and are generally more sensitive to chemical perturbations in comparison to solid tumors ^4,5,22^. Similarly, we computed the bimodality index for all genes using the gene expression of the solid tumors in The Cancer Genome Atlas (TCGA; 10534 tumors from 30 tissue types)^23^. We subsequently selected the protein-coding genes that showed high bimodality index (> 80th percentile) in both cancer cell lines and patient tumors (2816 out of 21903 genes; Fig 1C). Pathway enrichment analysis revealed a significant association of bimodal genes with G protein-coupled receptor signaling (GPCR) related pathways (Fig S1A), which are involved in the modulation of PI3K pathway, MAPK proteins, cAMP-dependent protein kinases, and cellular Ca2+^24,25^. Further characterization of these strongly bimodal genes revealed low redundancy (median: 0.03, IQR: 0.08) of their mRNA expression (Fig S1B).

### Development of interpretable models predictive of drug sensitivity

We implemented a machine learning approach based on logic-based models to identify reliable and interpretable biomarkers of sensitivity to different drugs. Logic-based models offer logic formulas using the ‘AND’, ‘OR’ and ‘NOT’ operators to build multivariate, nonlinear, yet interpretable predictive models. They overcome the limitations of univariate models that do not account for genes’ dependencies. To make such models broadly available, we developed RLOBICO, which is an R implementation of LOBICO method^26^, to find binary rules that predict sensitivity of samples to different drugs. To reduce the feature space and consequently the modeling computational cost, we used the ensemble minimum redundancy, maximum relevance (mRMRe) feature selection strategy (Fig 1A). The resulting models were represented as logic formulas including ≤ 10 genes to control the risk of overfitting and facilitate interpretation of the models. We assessed the predictive value of the logic models using the concordance index (CI; see Methods).

To fit the logic-based models, we used the pharmacogenomic data from the Cancer Therapeutics Response Portal (CTRP) by the Broad Institute, which represents the largest set of drug response data publicly available to date (version 2, including 544 drugs) ^5,22^ extracted from our PharmacoGx (version 1.14.0)^28^. We excluded drugs for which less than 10% of tested cancer cell lines are sensitive (area above the drug doseresponse curve [AAC] ≥ 0.2). Based on our approach, we were able to build models yielding a concordance index greater than 0.6 in a 5-fold cross-validation setting for 40% of the drugs in CTRPv2 (Fig 2A). The models cover a wide spectrum of drug classes such as EGFR signaling inhibitors and RTK signaling inhibitors (Fig 2B) supporting the generalizability of the predictive value of bimodal genes. The top-performing predictors include drugs targeting growth factor receptors such as EGFR, ERBB2 and VEGFR2. As mentioned earlier, the bimodal genes are enriched for several GPCR-related pathways. Transactivation of EGFR in cancer cell lines by GPCRs such as chemokine and angiotensin II receptors has been reported extensively^29–31^. Persistent transactivation of EGFR and ErbB2/HER2 by Protease-activated receptor-1 (PAR1), a GPCR activated by extracellular proteases, has been shown to promote breast carcinoma cell invasion ^32^. In addition, a strong complex formation between VEGFR2, another major growth factor, and the GPCR β2-Adrenoceptor has been reported resulting in VEGFR2 activation^33^. Among our top-performing models, we found that higher expression of fibroblast growth factor-binding protein 1 (FGFBP1) was correlated with increased sensitivity to Erlotinib (Fig 2C). FGFBP1 is a secreted chaperone that helps release fibroblast-binding factors (FGFs), stored in the extracellular matrix, and presents them to their cognate receptors, thereby enhancing FGF signaling. FGFBP1 mediated carcinogenesis has been implicated in many studies ^34^. According to Verbist et al. ^35^, FGFBP1 gene expression is downregulated by Erlotinib, resulting in decreased cell proliferation in cancer. These studies support our findings that high expression of FGFBP1 might be imparting sensitivity to Erlotinib via the inhibition of FGFBP1-FGF signaling axis. EGFR expression, a known biomarker for Erlotinib was excluded from our set of bimodal genes because its expression was not sufficiently bimodal in the TCGA cohort. Yet, we found a significant correlation between predictions based on rules that our method generated for Erlotinib and EGFR expression (PCC: 0.34, P-value: 8.03E-18) suggesting that our method was able to find a surrogate mimicking EGFR association with Erlotinib response. Moreover, predictions based on our method had better association with Erlotinib response (PCC: 0.36, P-value: 1.63E-20) than EGFR expression (PCC: 0.29, P-value: 2.17E-13). Another example of the top-performing models is that for Axitinib, (VEGFR inhibitor) in which low expression of G protein-coupled receptor, class C, group 5, member A (GPRC5A) was shown to be predictive of response (Fig 2C). GPRC5A, also known as Retinoic acid-induced gene 3 (RAI3) has been shown to elicit tissue-specific oncogenic and tumor-suppressive functions and is involved in the regulation of major cancer-related signaling pathways such as cAMP, NF-κB and STAT3^36–39^. Besides STAT3 and NF-κB signaling, GPRC5A is reported to impact cell cycle genes such as FEN1, MCM2, CCND1 and UBE2C in lung adenocarcinoma^40^. Knockout of GPRC5A has been reported to reduce proliferation and migration ability of PaCa cell lines and suppress the chemotherapy drug resistance of gemcitabine, oxaliplatin, and fluorouracil in PaCa cells^41^. Knockdown of GPRC5A has also been found to negatively impact FAK/Src activation, and RhoA GTPase activity, the key mediators of VEGF signaling in cancer cell lines^42–44^. These findings support a possible mechanism for Axitinib sensitivity imparted by low expression of GPRC5A, via VEGF-activated signaling intermediates. All trained models (CI > 0.6) from CTRPv2 and their associated predictive rules are shared in the supplementary data (Supplementary File 1).

**Fig 2:**
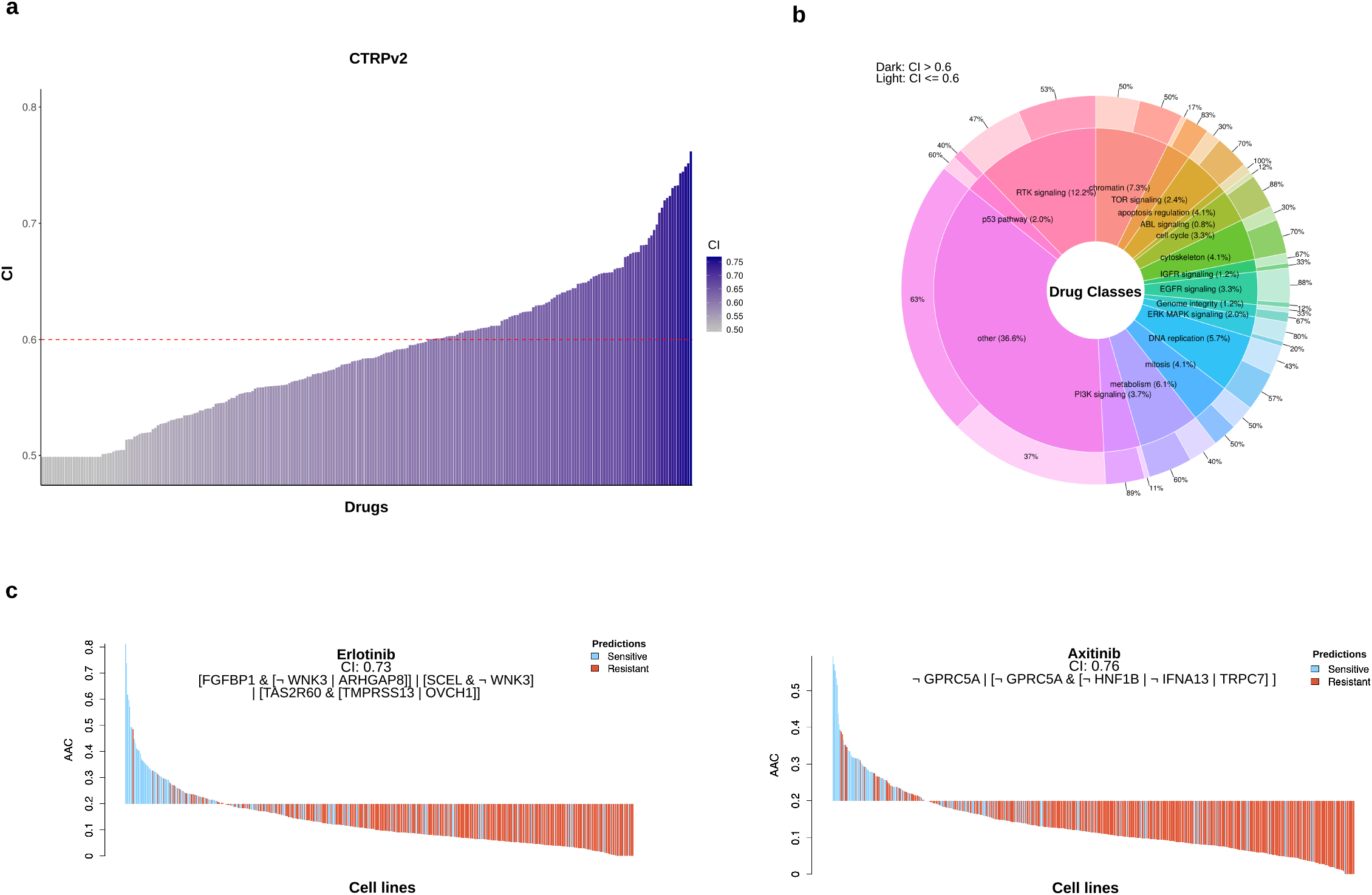
**(a)** Performance of developed logical models on the training data set for each drug in CTRPv2. Red-dashed line represent cutoff for good and bad models. **(b)** Distribution of good (CI>0.6; dark color) and bad (CI<=0.6; light color) models (outer ring) for each drug class in CTRPv2 and distribution of drug classes in CTRPv2 (inner ring). **(c)** Examples of top performing trained logical models along with the rules predicted to assess sensitivity to the respective drugs

### Validation of predictors

Recognizing that large-scale pharmacogenomic studies employ complex, potentially noisy experimental protocols ^19,45,46^, it is crucial to validate the performance of our new predictors in fully independent datasets (using both independent genomic and pharmacological profiles of cancer cell lines ^47^) to assess their generalizability. We, therefore, validated our models on two large pancancer pharmacogenomic datasets, namely the Genentech Cell Line Screening Initiative^14,46^ (gCSI, released in 2018) and the Genomics of Drug Sensitivity in Cancer^2,3^ (GDSC2, released in 2019), both included in our PharmacoGx package^28^. Among all the models in common with gCSI, our models achieved 92.3% validation rate (CI > 0.6 for 13 out of 27 drugs in common with CTRPv2; Fig 3A). On GDSC2, our models achieved a validation rate of 61% (CI > 0.6 for 16 out of 26 drugs in common with CTRPv2; Fig 3B).

**Fig 3:**
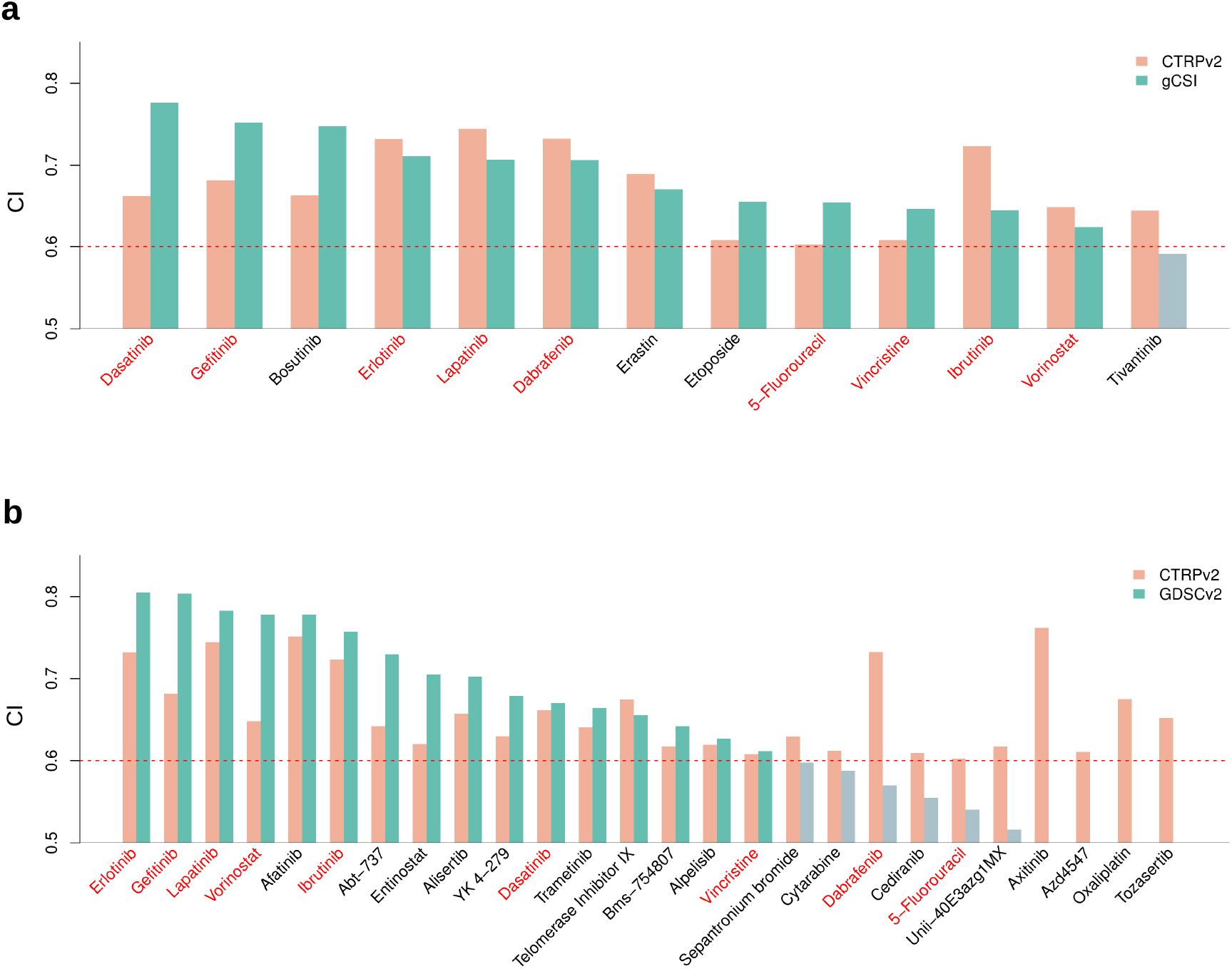
Validating developed logic models on external datasets; **(a)** gCSI, **(b)** GDSCv2. Red colored drugs are common between gCSI and GDSCv2.

There were 7 out of 9 (78%) predictive models that were validated on both external datasets (Fig 3), strongly supporting the generalizability of the logic rules predictive of drug response. The logic model predictive of Erlotinib response described previously (Fig 2C) yielded high predictive value in both independent datasets (CI of 0.79 and 0.73 in GDSC2 and gCSI, respectively). Dasatinib, whose predictive logic model was also validated in GDSC2 and gCSI, showed association to several genes including High Mobility Group AT-Hook 2 (HMGA2). HMGA2 is a member of the high motility group (HMG) protein family that binds to the DNA minor groove at sequences rich in A and T nucleotides, and acts as a transcriptional regulator. Apart from its role as a transcriptional co-regulator, HMGA2 has been found to induce epithelial-to-mesenchymal transition in lung cancer^48^. HMGA2 also functions as a positive regulator of cell proliferation and its expression is implicated as a prospective diagnostic biomarker in the assessment of endometrial serous cancer^49^. According to Turkson J et al.^50^, nuclear Src and p300 associate with HMGA2 promoter and regulate its gene expression in PDAC patient samples. Src inhibition by Dasatinib might negatively impact HMGA2 mediated cell oncogenesis, resulting in sensitivity in cancers with high HMGA2 expression as predicted in our study. Among the other top-performing drugs, the sensitivity of Gefitinib, an EGFR inhibitor has been attributed to the expression of ARHGAP8, a gene implicated in EGFR-mediated ERK1/2 phosphorylation and oncogenesis^51–53^. The expression of other bimodal genes associated with lapatinib sensitivity such as MARVELD3 and EPN3 has been reported to promote migration and invasion of cancer cells^54–56^.

### Bimodality of gene expression outperforms genomics as a source of predictive biomarkers

To test whether the gene expressions of the top bimodal genes compose a richer feature set for predicting drug response than other data types such as tissue of origin, mutation and copy-number-variation (CNV), we systematically analyzed all the data types by running them through the same computational pipeline used for bimodal genes. Our results indicate that the expression of bimodal genes significantly outperformed the other data types (mutations and CNVs) in 72% of the drugs (Fig 4 A and B). Tissue type of the sample was found to be the best model predicting sensitivity to 16% of the drugs suggesting a strong specificity of drug response^57^ (Fig 4D). Dabrafenib, for example, is an inhibitor of BRAF serine-threonine kinase that was predicted by our model to show a high association with skin cancer (Fig 4D). This drug is indeed approved by FDA as a single agent for the treatment of patients with unresectable or metastatic melanoma with BRAF V600E^58^. We also found that predictions based on bimodal genes were different than predictions based on tissues (Fig 4C). Mutation and CNV features were found to be the best in predicting sensitivity in 11.2% of the drugs (Fig 4 E and F). An example of these drugs is Nutlin-3A, an MDM2 inhibitor that activates wild-type p53 mutation^59,60^. TP53 wild-type mutation was predicted by our approach to indicate sensitivity to Nutlin-3A (Fig 4E). This outperformance of expression data in comparison to other data types conforms with previous studies and community efforts that investigated the relevance of different data types to predict drug sensitivity and showed that gene expression has more rich information and predictive power than other data types^7^. These results also suggest that combining these different data types in a multi-omics model could improve the resultant predictors given the heterogeneity of the chosen feature sets we observed for different drugs (Fig 4A).

**Fig 4:**
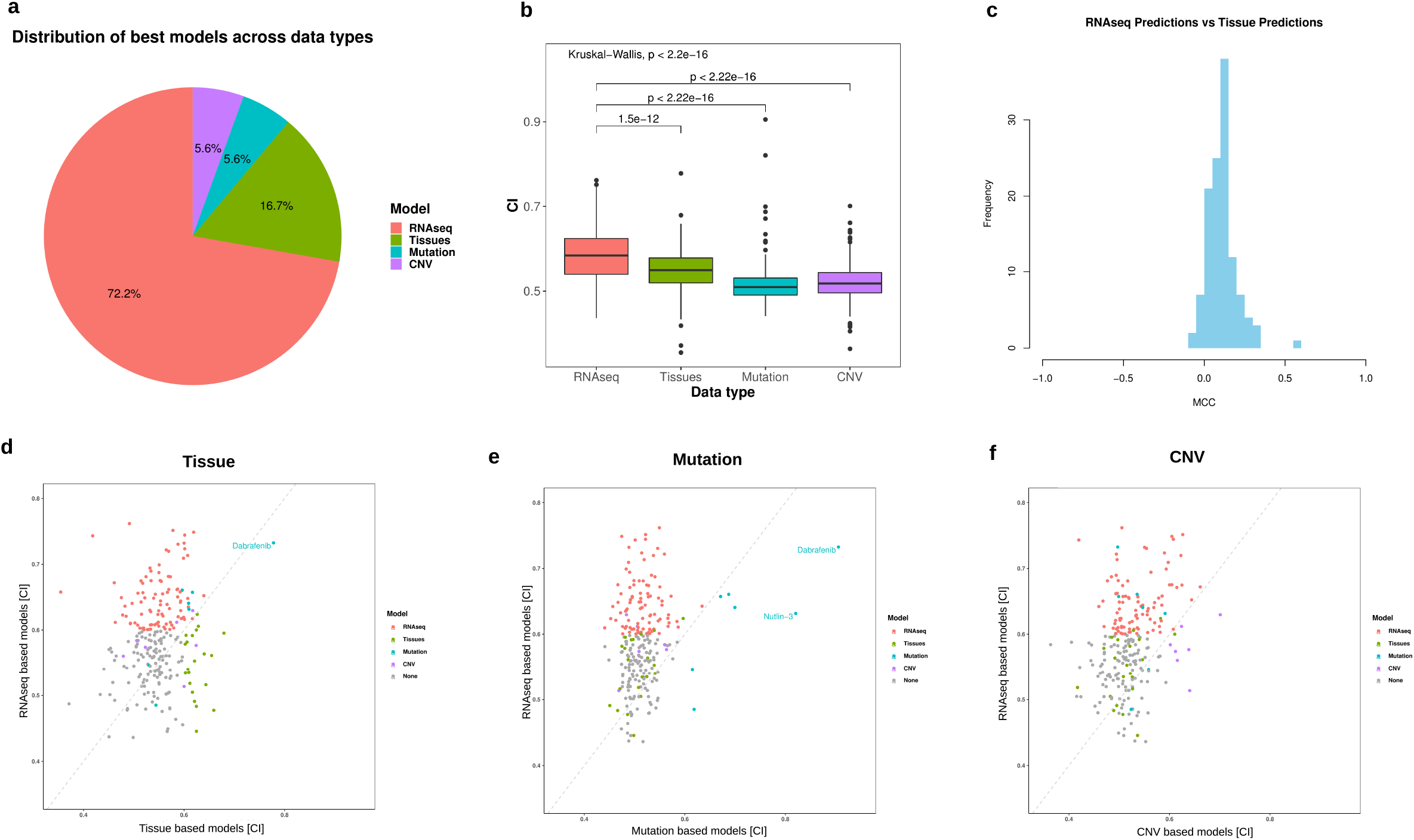
**(a)** Distribution of best models across data types. **(b)** Statistical comparison between models across data types. p-values are based on Wilcoxon signed-rank test. **(c)** Comparison between RNAseq-based predictions and Tissue-based predictions (median: 0.11, IQR: 0,09). **(d,e,f)** Comparing RNAseq based models with: **(d)** Tissues, **(e)** Mutation, **(f)** CNV. color indicates best models across all data types.

### Tissue-specific models

Heterogeneity within cancer tissues constitutes another layer of complexity. We investigated whether bimodal genes within a specific tissue could generate a more accurate predictor of sensitivity for samples of that tissue type. Lung cancer was chosen as a case study, given the number of samples available in both CCLE and TCGA to extract reliable bimodal genes. We applied our pipeline to these samples and developed logic-based models with minimum predictive value (CI > 0.6) for about 30% of drugs in CTRPv2. ABT-737, a selective inhibitor of BCL-2 that showed a therapeutic effect in lung cancer, was among the best performing models we found (CI = 0.78). We validated our predicted rules on an external dataset of lung cancer samples in GDSCv2 screened with ABT-737 (CI = 0.73). We then compared the lung-specific rules based on lung-specific bimodality with pan-cancer rules in predicting drug response in lung samples. We compared the pancancer and tissue-specific models on lung samples in gCSI and GDSC2 and found that both features sets yielded similar associations with response (Fig 5). These results suggest that both sets of rules can be predictive of drug response and provide different levels of biomarker granularity.

**Fig 5:**
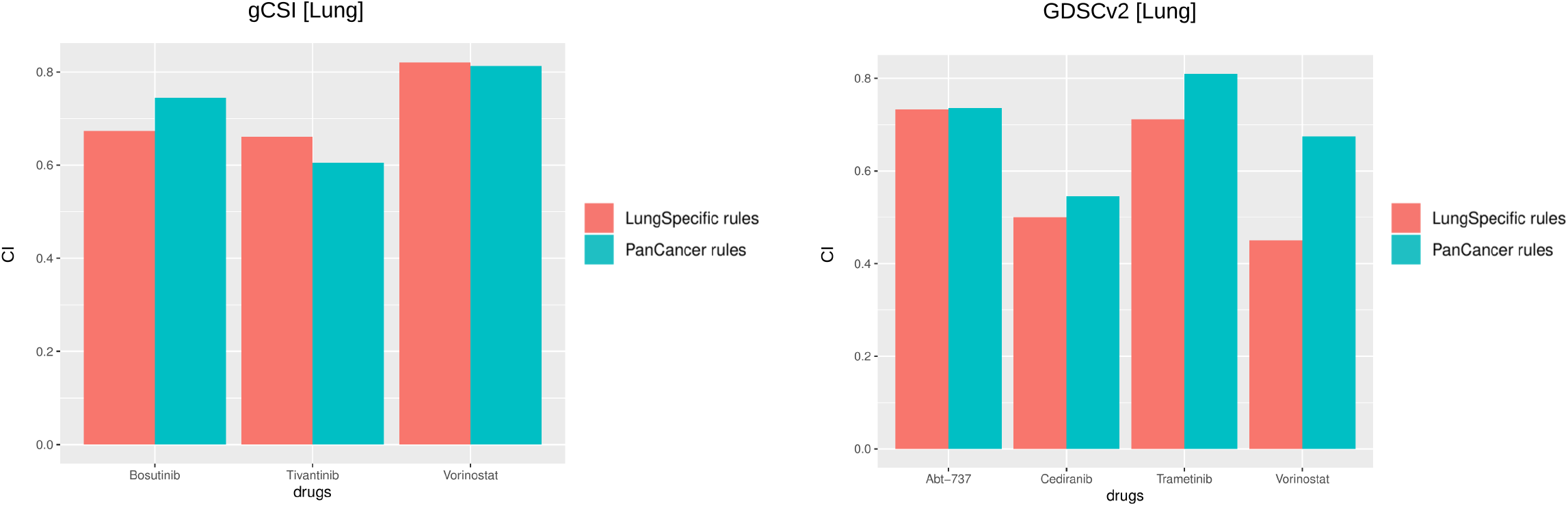
Comparing lung-specific rules vs pan-cancer rules in predicting drug response within lung samples in: **(a)** gCSI, and **(b)** GDSCv2.

## DISCUSSION

Bimodality of gene expression represents an interesting phenomenon associated with several biological processes. One of the advantages of bimodal genes as a biomarker is that it can be used to robustly classify samples into two distinct expression states based on its modes, allowing for easier interpretation and translation of the biomarker into the clinic. In this study, we showed that top bimodal genes are mostly associated with extracellular membrane pathways which have a downstream effect on important cancer-related processes such as MEK and PI3K signaling. We introduced the largest comprehensive set of bimodal genes derived from a large panel of cancer cell lines tested against hundreds of drugs and patient data from TCGA. We found a high correlation between the bimodality scores of the corresponding genes within the cell line and patient datasets (PCC: 0.695, p < 2.2e-16), which showcases the reliability of the chosen genes to be globally bimodal within cancer. We found a subset of genes that exhibited a bimodal distribution in one dataset but not the other probably due to differences in tissue distribution of samples, or to intrinsic transcriptional differences between the in vitro models and the patient tumors.

Although the bimodality of expression provides multiple advantages in biomarker discovery, restricting the modeling to only bimodal genes filters out many known drug biomarkers because their expressions do not follow a bimodal distribution. EGFR expression, for example, is a known biomarker for Erlotinib. However, it is not bimodal in TCGA which excluded it from our set of bimodal genes that we used for training the models. Yet, we found a high concordance between predictions based on rules our method generated for Erlotinib and EGFR expression (PCC: 0.34, P-value: 8.03E-18) suggesting that our method was able to find a surrogate mimicking EGFR association with Erlotinib response. Moreover, predictions based on our method had better association with Erlotinib response (PCC: 0.36, P-value: 1.63E-20) than EGFR (PCC: 0.29, P-value: 2.17E-13). Despite the constraint on the number of bimodal genes we use, we have shown that this set of features along with our novel method of applying logic-based models were able to predict sensitivity to 101 drugs from 17 different drug classes suggesting global utility of these features (Fig 2B).

We also showed that bimodal genes outperformed other data types, mutations and copy number variations, in predicting sensitivity to different drugs. An interesting follow-up to this analysis would be to investigate the complementary effect of merging these data types in building more accurate models. Challenges that we anticipate are the availability of data types across datasets, data normalization and computational complexity to query the larger search space for candidate rules.

Finally, investigating the bimodality as a source for biomarkers within tissues showed promising results that suggest a more in-depth association within tissue-specific cancer subtypes that would not be captured in pan-cancer studies. This variation in defining bimodal genes is mostly due to the difference in the distributions of genes within tissues and across different cancer types. A challenge that we anticipate is the lack of sufficient samples within different tissue types to generate a reliable and robust set of bimodal genes within each tissue type.

## CONCLUSION

Finding reliable and interpretable biomarkers that can predict patients’ response to drugs remains a formidable challenge. We showed that bimodally expressed genes represent an interesting subset of features for biomarker discovery and that they cover important cancer-associated pathways. Our results, utilizing logic-based models to generate rules that predict sensitivity to drugs, show that we can predict biomarkers based on bimodal genes with high accuracy and validation rate across datasets. These bimodal predictive biomarkers have a high potential of clinical translatability given the clear separation they provide between patient cohorts who would and would not benefit from different drugs, and the practicality of measuring few genes for treatment planning using various low-throughput assays instead of whole-genome sequencing.

## METHODS

### Datasets

CCLE, CTRPv2, gCSI, GDSC and TCGA were all processed using the same pipeline utilizing the PharmacoGx R package pipeline. Gene expression profiles were generated using Kallisto pipeline^61^ with GRCh38 as human reference.

### Bimodality of gene expression profiles

Gene expression profiles, obtained from CCLE dataset, were used to characterize the bimodality feature of each gene in the set by fitting its distribution into a mixture of two Gaussian distributions. For those genes with a good fit, a bimodality score was calculated using the following formula:

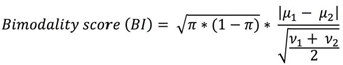

where is the proportion of samples in one group, μ_1_ and μ_2_ are the means of the expression level of the two modes; and _1_ and _2_ are the variances. Similar characterization was done using TCGA dataset. Genes, then, were ranked according to their bimodality scores and the common protein-coding genes in the top 80th percentile of bimodality scores distribution in both CCLE and TCGA were chosen as top bimodal genes feature set. A binarization cutoff for each gene distinguishing relatively low vs high expression was calculated by taking the average point between the modes of the two fitted gaussian distributions.

### Logic-based models

Logic-based models are machine learning models aiming at constructing boolean logic functions that model the relationship between a binary set of features and a class label. Interpretability of the modeled associations is a key advantage of these types of models in comparison to other traditional machine learning models, which is an important feature for clinical translation of biomarkers. We developed RLOBICO, which is an R implementation of LOBICO method ^26^, to find binary rules that predict sensitivity of samples to different drugs. Our proposed pipeline starts with a binarized expression matrix followed by a feature selection method (mRMRe) to choose highly relevant and complementary features that are then fed into RLOBICO to search the space of possible rules and associate these rules with a drug effect.

For each drug, we create a binarized expression matrix based on top bimodal genes features’ set and represent the effect of the drug on samples using the area above the dose-response curve (AAC) metric. LOBICO requires binarizing the effect of the drug and so we chose AAC of 0.2^62^ as a threshold classifying samples to be either resistant (AAC < 0.2) or sensitive (AAC > 0.2) to each drug. However, the continuous values of AACs are still used as weights to optimize the modeling step such that a higher penalty would be incurred if a highly sensitive sample was misclassified as resistant. Generated rules by LOBICO are described using the disjunctive normal form, which is a standard notation to express logic functions. The disjunctive normal form is parameterized by two parameters: K, the number of disjuncts, and M, the number of terms per disjunct. We varied K and M to represent models of different complexities, i.e. from single predictors (K=1,M=1) to more complex models [(K, M): (1,2), (1,3), (1,4), (2,2), (2,1), (3,1), (4,1)]. We use mRMRe to limit the search space of all possible logical combinations of features to the top ten highly relevant and complementary features to control the risk of overfitting and facilitate interpretation of the model. We then apply RLOBICO to find the best rule predicting sensitivity of samples to drugs. Finally, to achieve more robust results, we create an ensemble rule based on a majority vote from rules generated by three different mRMRe features sets followed by RLOBICO. For evaluation of models, we use a modified version of the concordance index (CI) [https://github.com/bhklab/wCI]. This modification accounts for noise in the drug screening assays as we found that repeating the same drug-cell line experiment in CTRPv2 resulted in inconsistencies in terms of measured drug response (AAC). We further investigated this observation and found that 95% of the replicates of the same drug and cell line experiments showed differences (Δ AAC = | AAC_replicate1_ - AAC_replicate2_ |) within 0.2 range (Fig S2). Hence, we remove the pairs of AACs that have Δ AAC < 0.2 from the calculation of the regular CI as they can flip directions within that range randomly.

### Research reproducibility

CCLE, CTRPv2, gCSI and GDSC2 can be downloaded using PharmacoGx R package^28^. Code to reproduce the results and figures is available at https://github.com/bhklab/Gene_Expression_Bimodality. RLOBICO R package was used to generate the logic-based models (https://github.com/bhklab/RLOBICO).

## ACKNOWLEDGEMENTS

The authors would like to thank the investigators of the Genomics of Drug Sensitivity in Cancer (GDSC), the Cancer Cell Line Encyclopedia (CCLE), Genentech (gCSI), and the Cancer Therapeutics Response Portal (CTRP) who have made their valuable pharmacogenomic data available to the scientific community.

---

**Fig S1:**
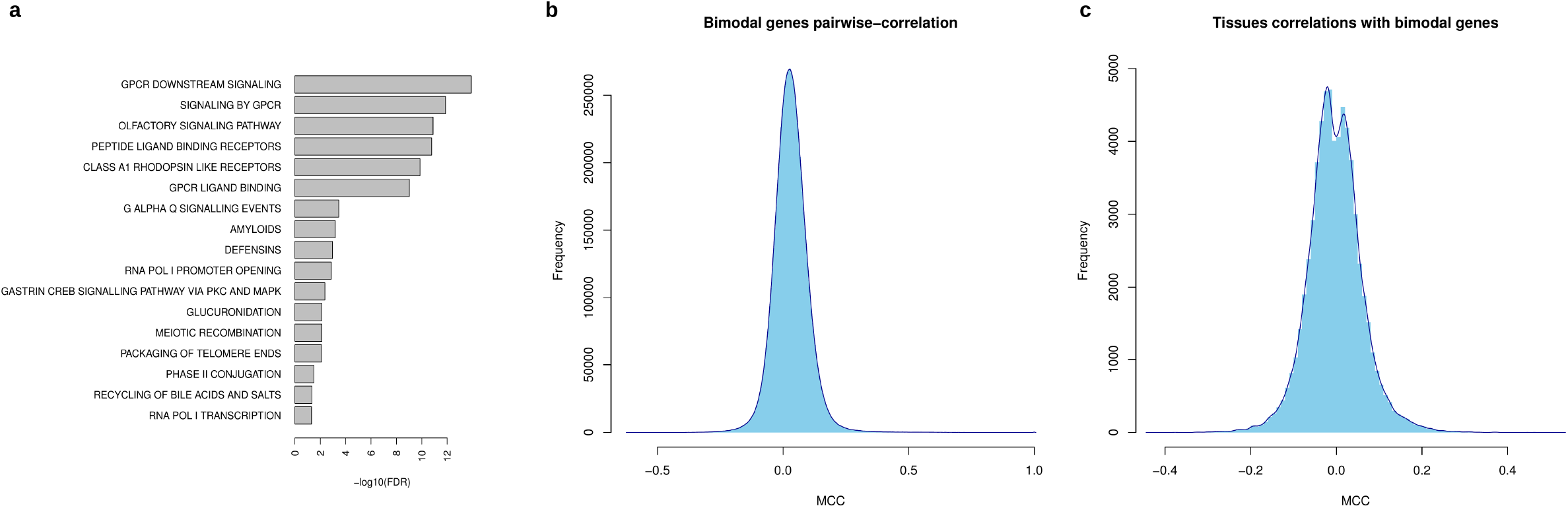
**(a)** Pathways enriched with the set of bimodal genes. **(b)** Pairwise correlation between all global bimodal genes using Matthews correlation coefficients. **(c)** correlation between all global bimodal genes and tissues.

**Fig S2:**
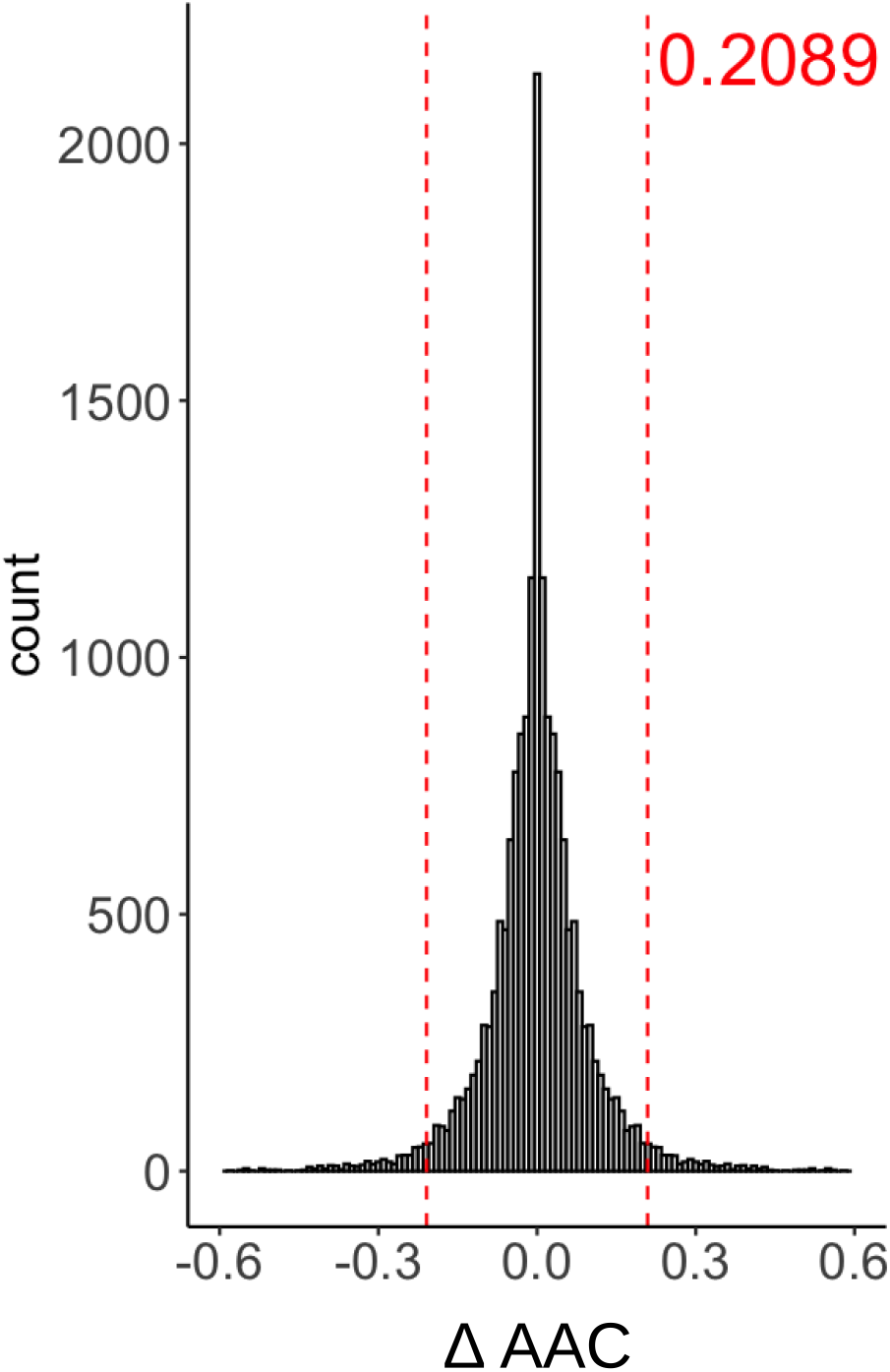
Difference in drug response (AAC) for the same drug-cell line experiments in CTRPv2

**Supplementary File 1: All models that yielded CI > 0.6 on CTRPv2 data**

