## Supplementary File 1 for "Bimodality of gene expression in cancer patient tumors as interpretable biomarkers for drug sensitivity"

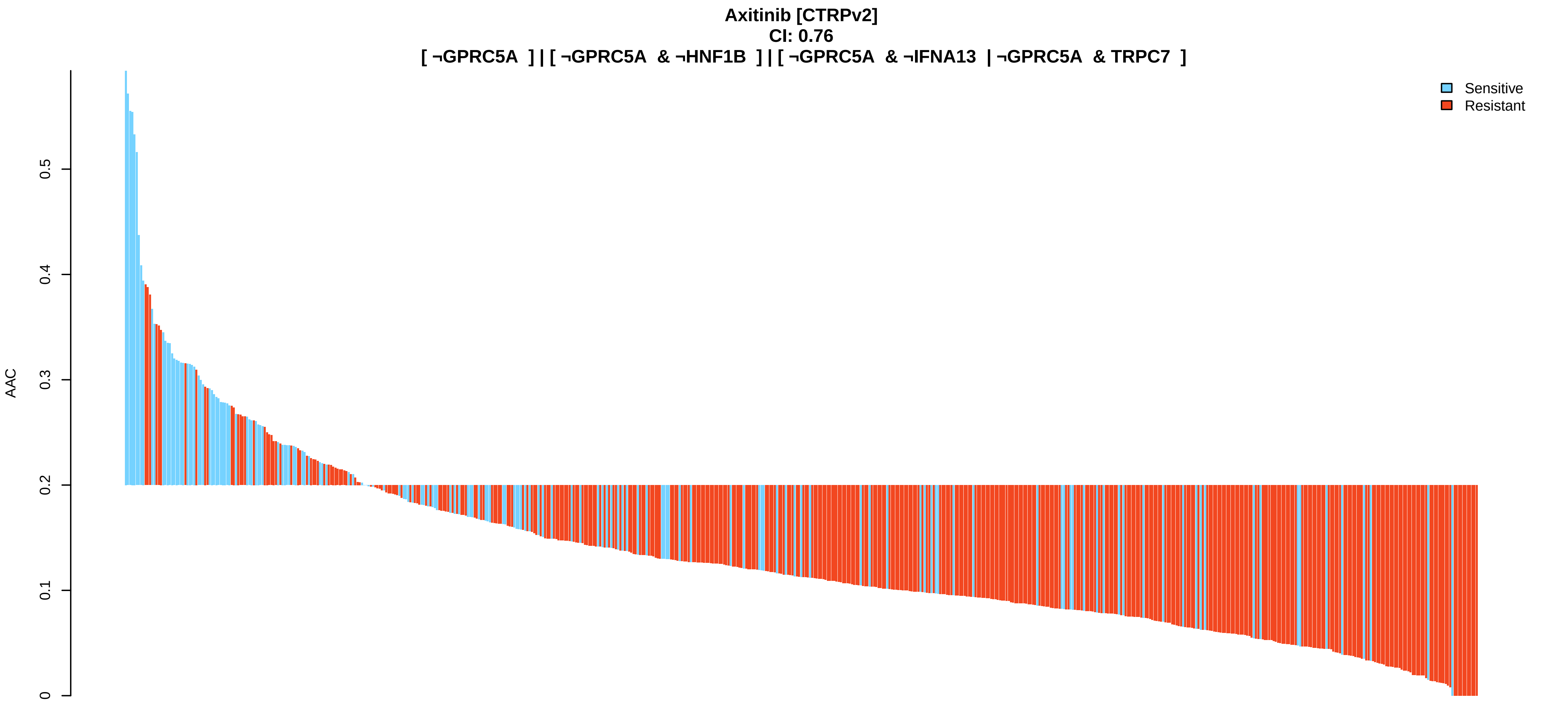
Axitinib [CTRPv2]
CI: 0.76
[ ¬GPRC5A ] | [ ¬GPRC5A & ¬HNF1B ] | [ ¬GPRC5A & ¬IFNA13 | ¬GPRC5A & TRPC7 ]
Sensitive
Resistant
AAC
0
0.1
0.2
0.3
0.4
0.5

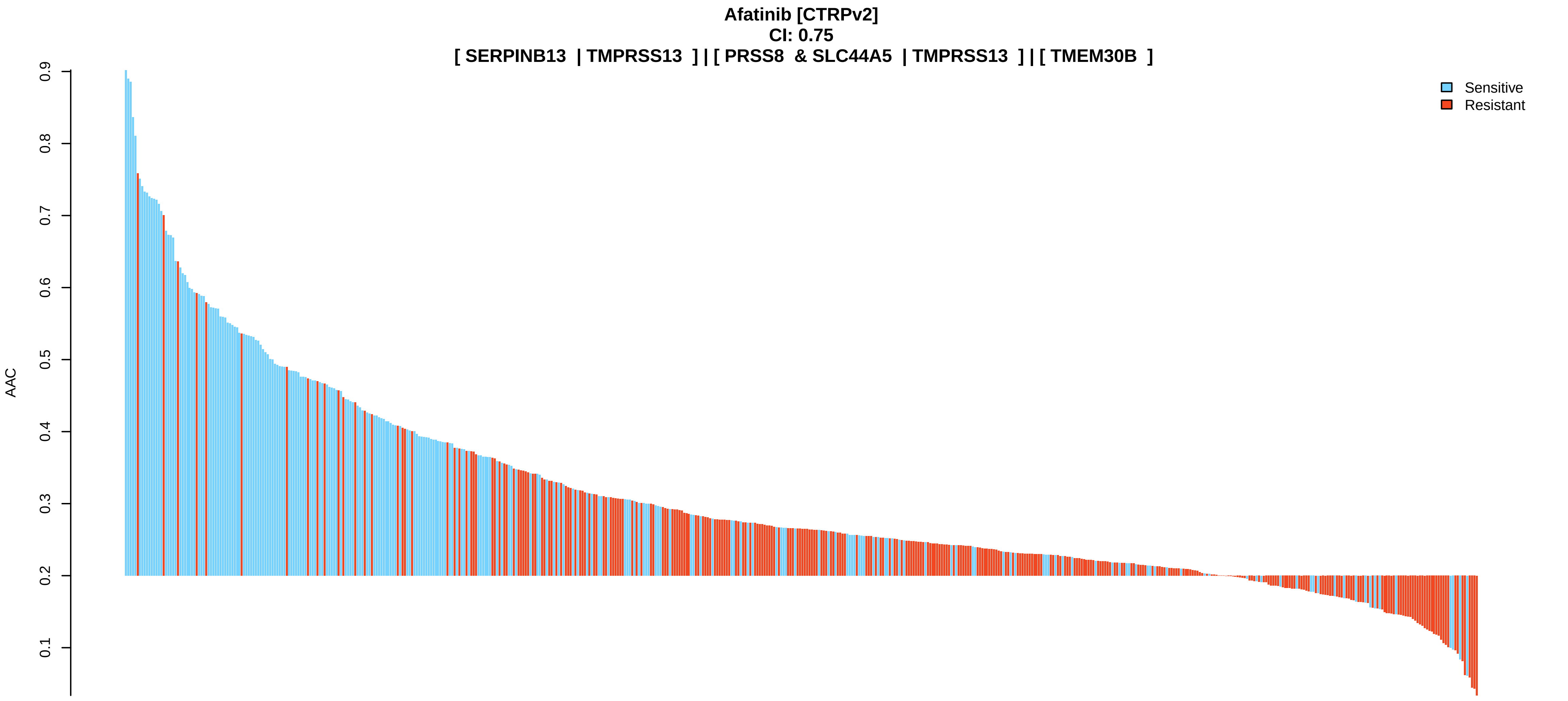
Afatinib [CTRPv2]
CI: 0.75
[ SERPINB13 | TMPRSS13 ] | [ PRSS8 & SLC44A5 | TMPRSS13 ] | [ TMEM30B ]
Sensitive
Resistant
AAC
0.1
0.2
0.3
0.4
0.5
0.6
0.7
0.8
0.9

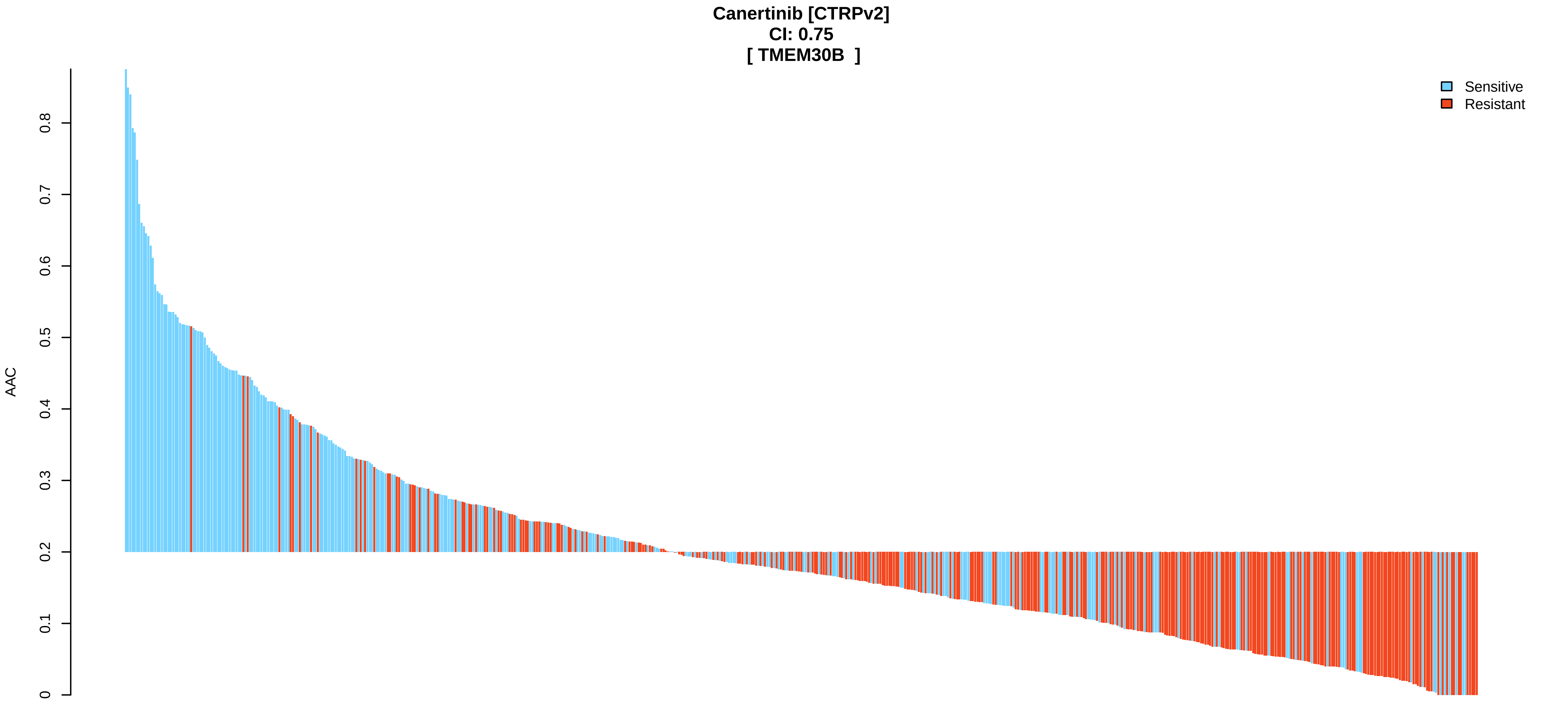
Canertinib [CTRPv2]
CI: 0.75
[ TMEM30B ]
Sensitive
Resistant
AAC
0
0.1
0.2
0.3
0.4
0.5
0.6
0.7
0.8

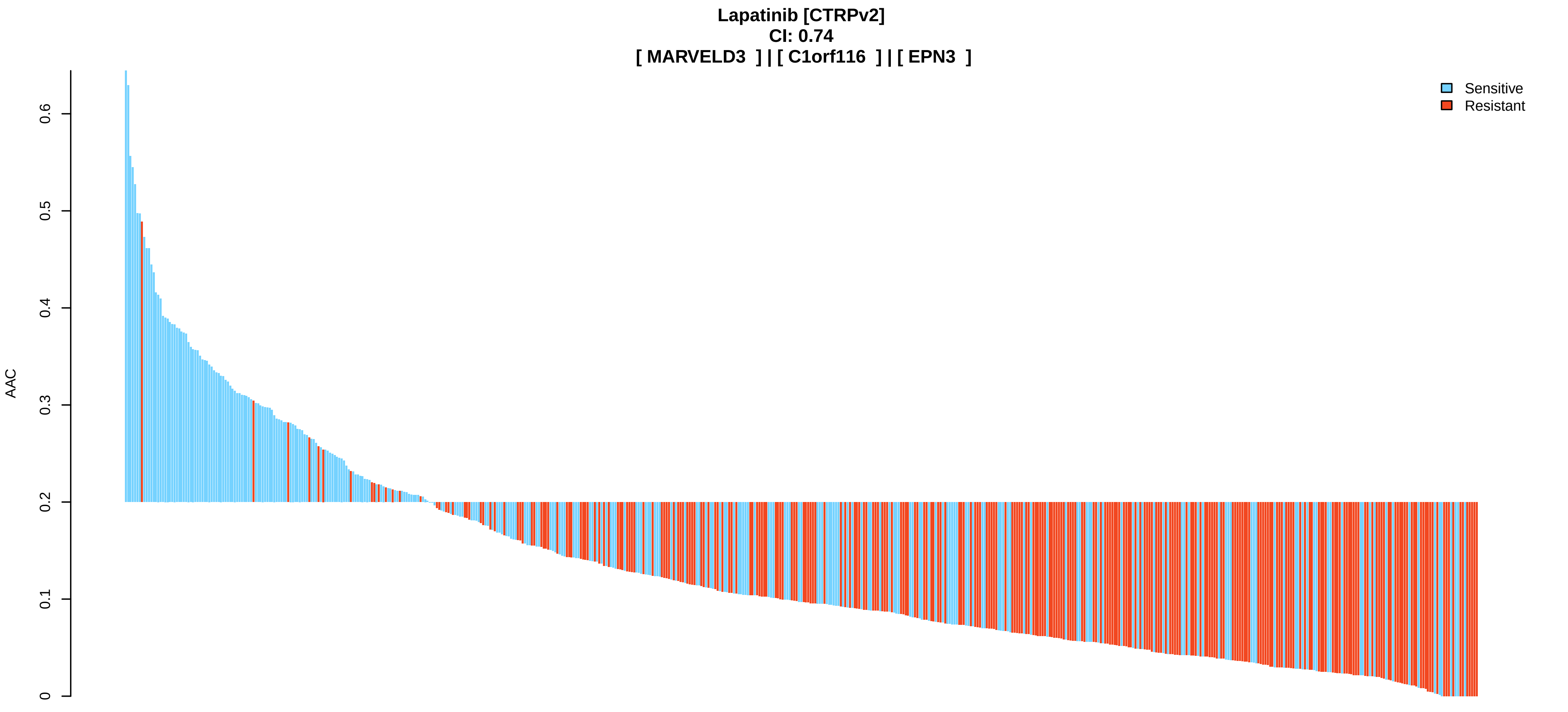
Lapatinib [CTRPv2]
CI: 0.74
[ MARVELD3 ] | [ C1orf116 ] | [ EPN3 ]
AAC
0
0.1
0.2
0.3
0.4
0.5
0.6
Sensitive
Resistant

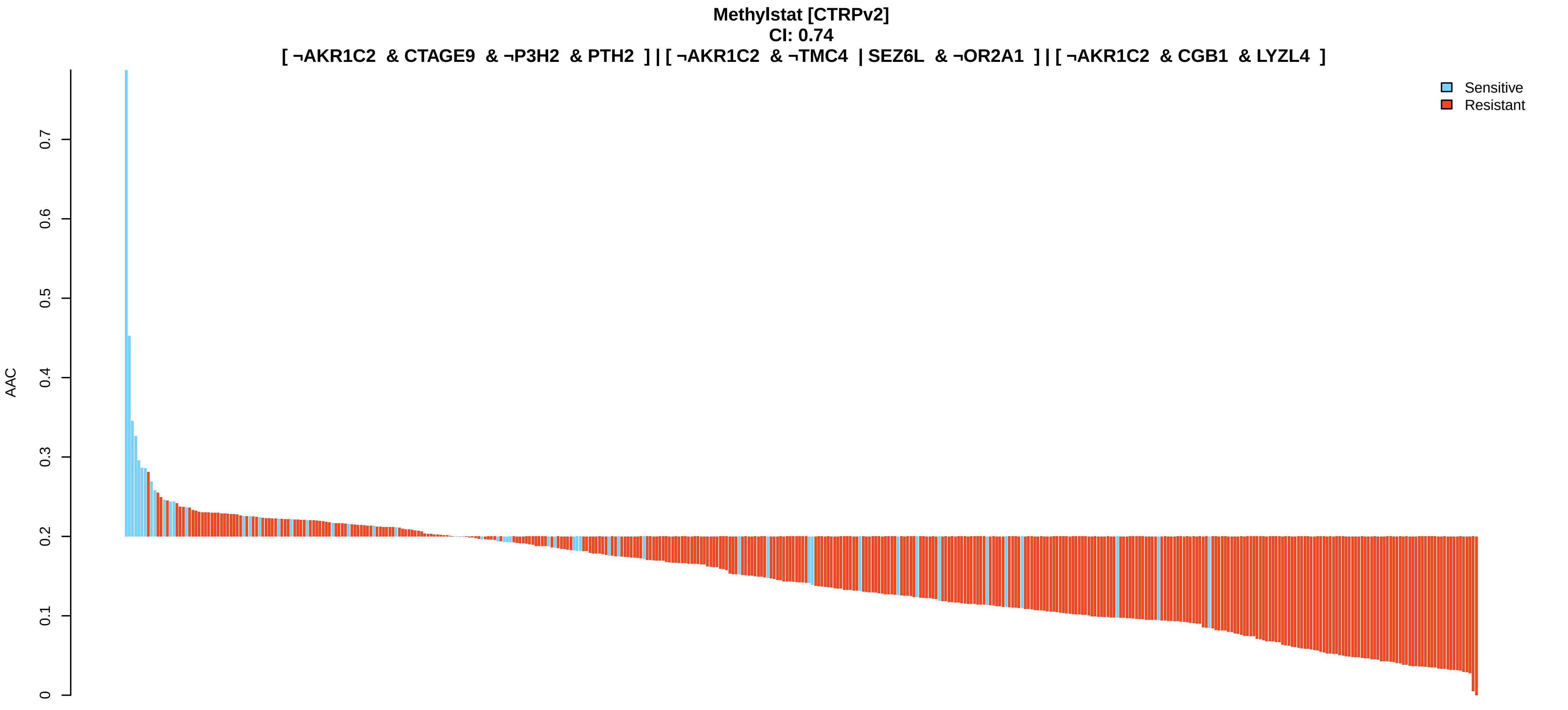
Methylstat [CTRPv2]
CI: 0.74
[ ¬AKR1C2 & CTAGE9 & ¬P3H2 & PTH2 ] | [ ¬AKR1C2 & ¬TMC4 | SEZ6L & ¬OR2A1 ] | [ ¬AKR1C2 & CGB1 & LYZL4 ]
AAC
0
0.1
0.2
0.3
0.4
0.5
0.6
0.7
Sensitive
Resistant

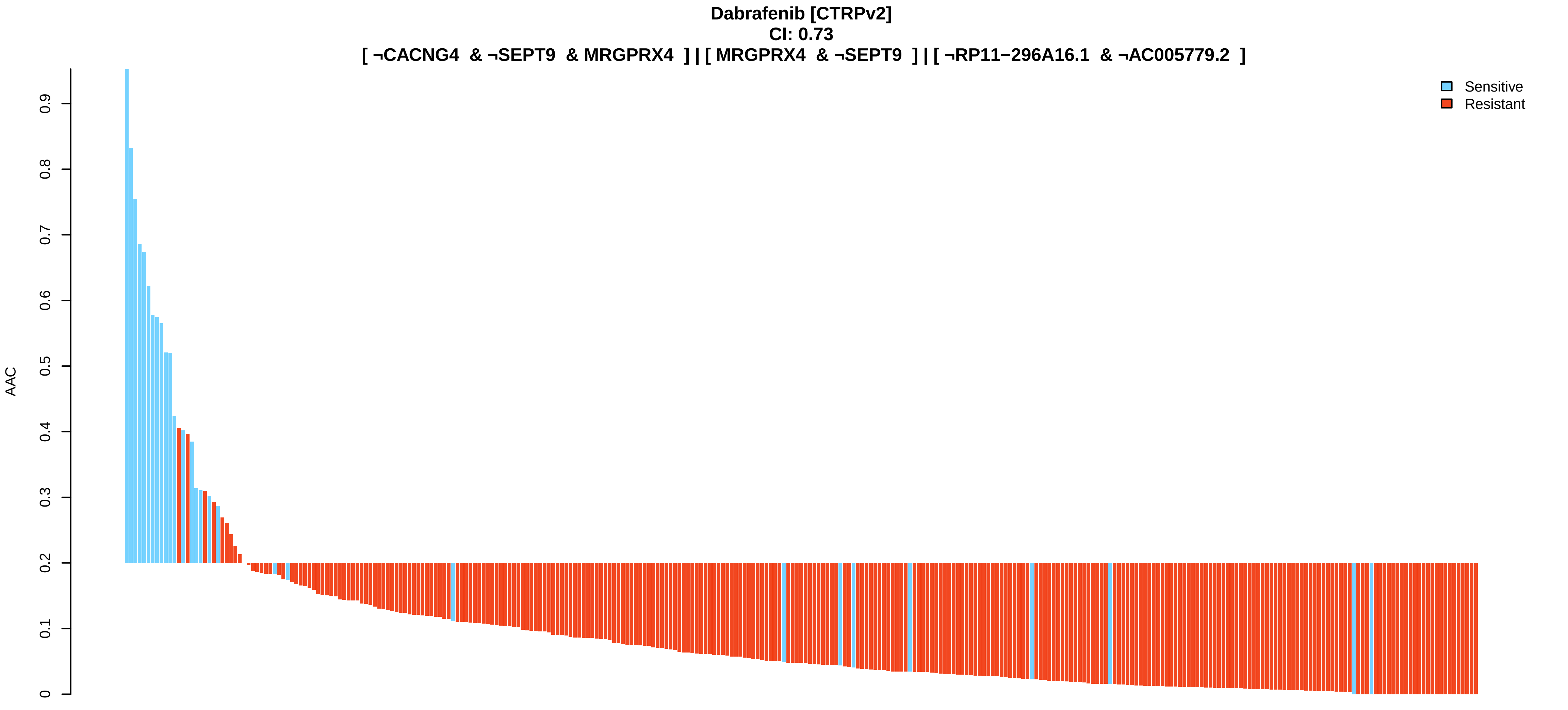
Dabrafenib [CTRPv2]
CI: 0.73
[ ¬CACNG4 & ¬SEPT9 & MRGPRX4 ] | [ MRGPRX4 & ¬SEPT9 ] | [ ¬RP11−296A16.1 & ¬AC005779.2 ]
AAC
0
0.1
0.2
0.3
0.4
0.5
0.6
0.7
0.8
0.9
Sensitive
Resistant

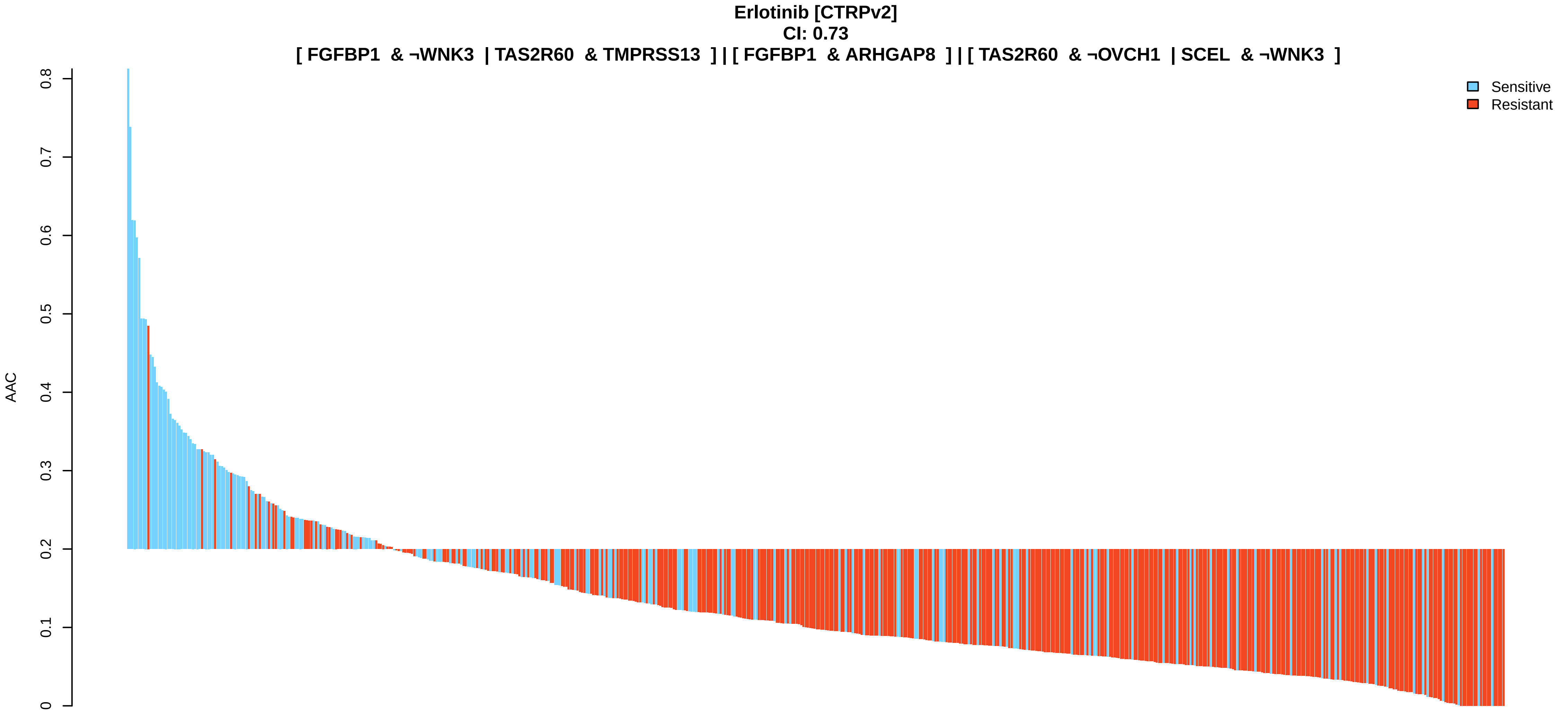
Erlotinib [CTRPv2]
CI: 0.73
[ FGFBP1 & ¬WNK3 | TAS2R60 & TMPRSS13 ] | [ FGFBP1 & ARHGAP8 ] | [ TAS2R60 & ¬OVCH1 | SCEL & ¬WNK3 ]
AAC
0
0.1
0.2
0.3
0.4
0.5
0.6
0.7
0.8
Sensitive
Resistant

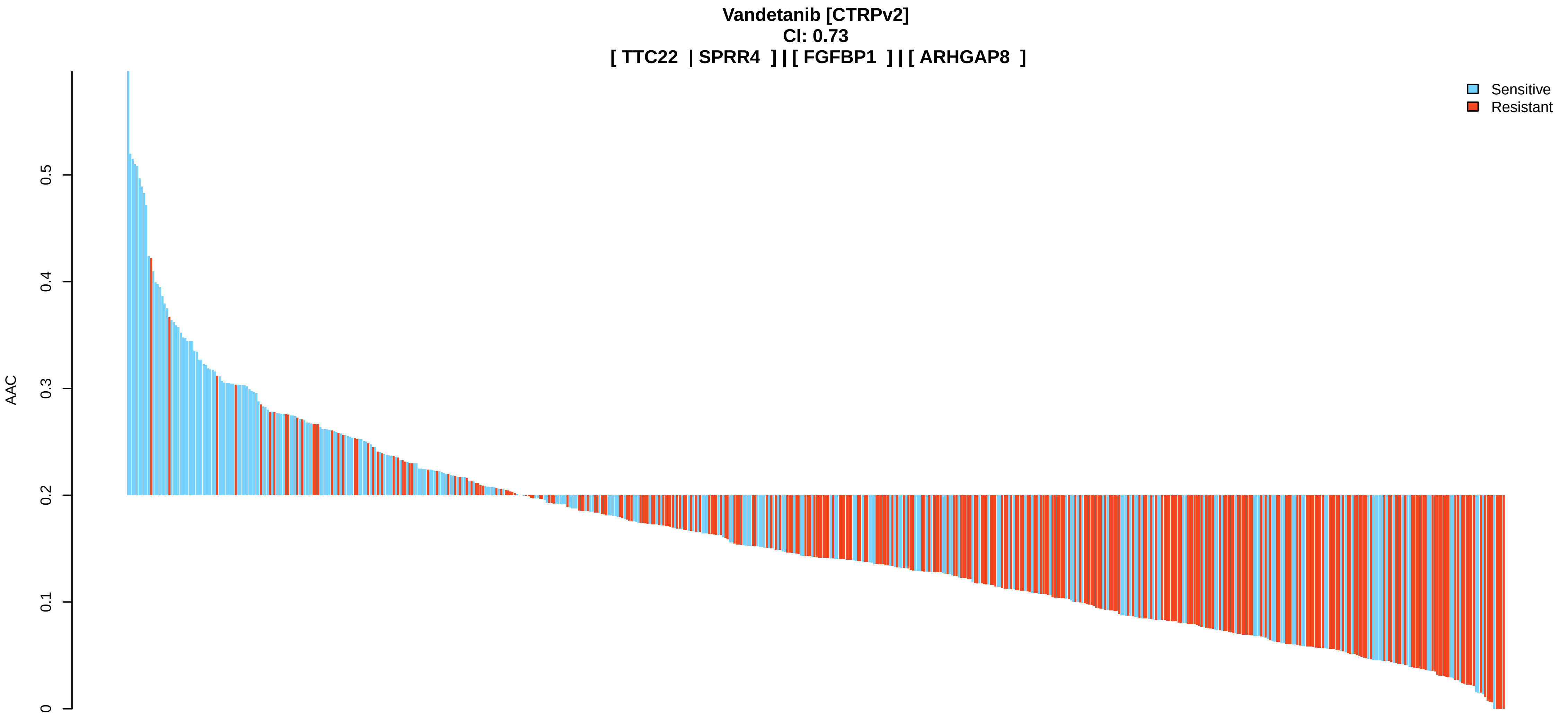
Vandetanib [CTRPv2]
CI: 0.73
[ TTC22 | SPRR4 ] | [ FGFBP1 ] | [ ARHGAP8 ]
AAC
0
0.1
0.2
0.3
0.4
0.5
Sensitive
Resistant

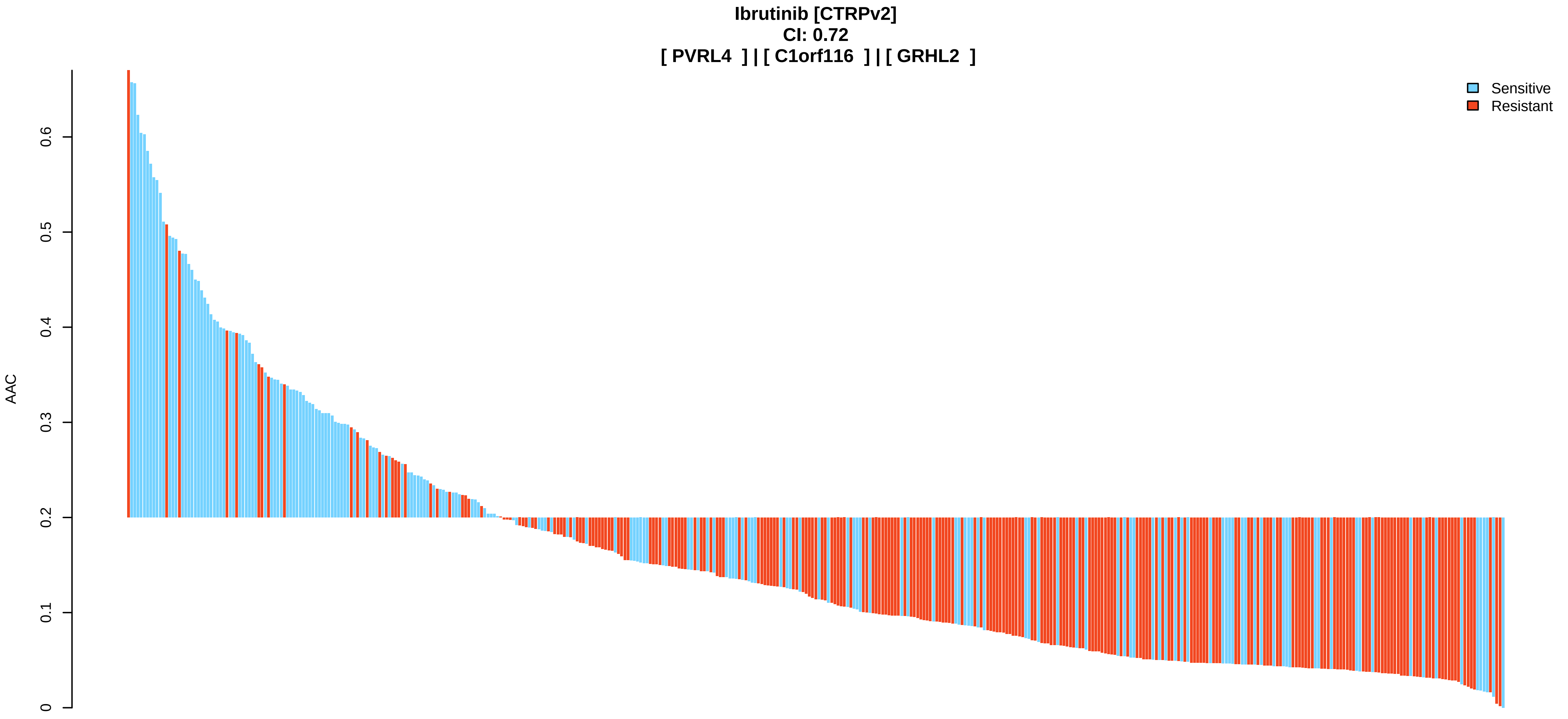
Ibrutinib [CTRPv2]
CI: 0.72
[ PVRL4 ] | [ C1orf116 ] | [ GRHL2 ]
AAC
0
0.1
0.2
0.3
0.4
0.5
0.6
Sensitive
Resistant

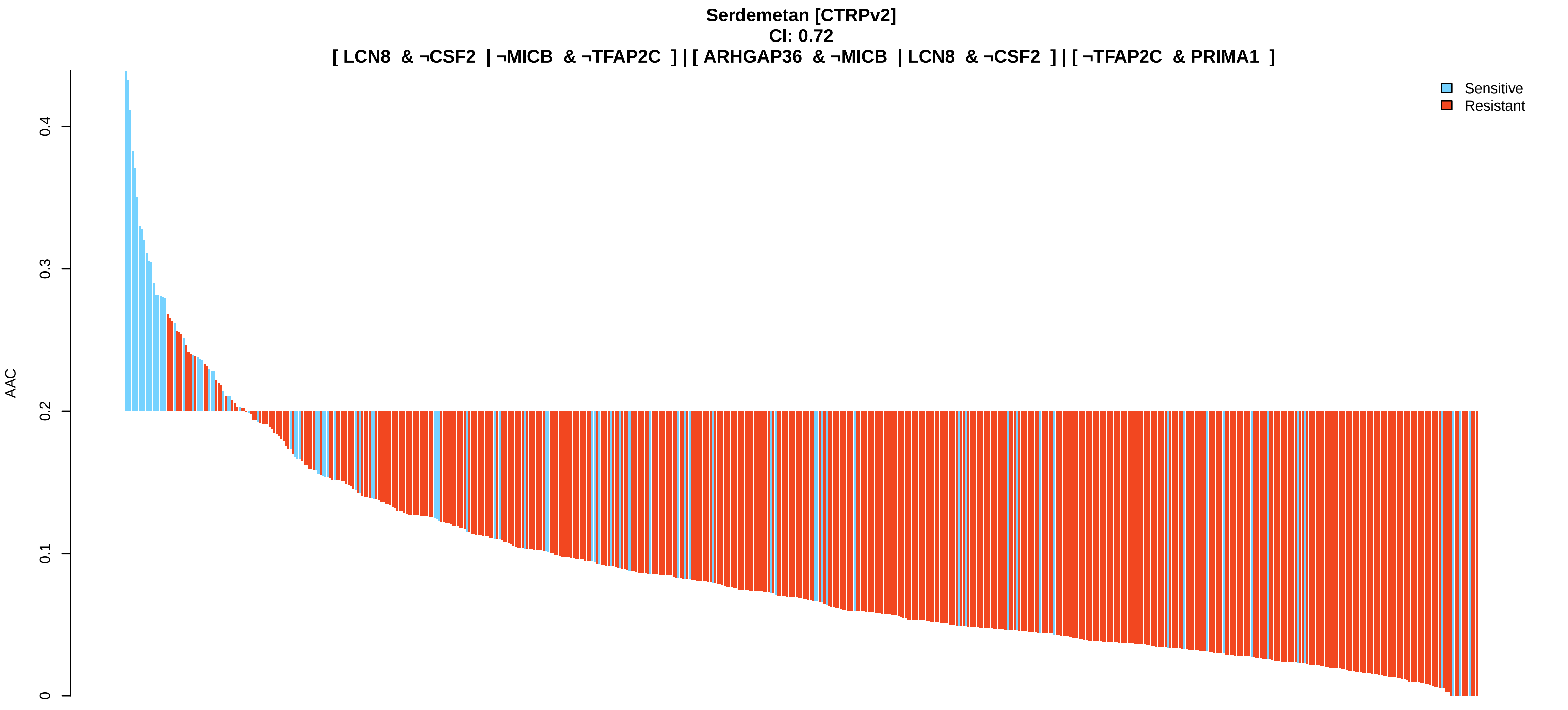
Serdemetan [CTRPv2]
CI: 0.72
[ LCN8 & ¬CSF2 | ¬MICB & ¬TFAP2C ] | [ ARHGAP36 & ¬MICB | LCN8 & ¬CSF2 ] | [ ¬TFAP2C & PRIMA1 ]
Sensitive
Resistant
AAC
0
0.1
0.2
0.3
0.4

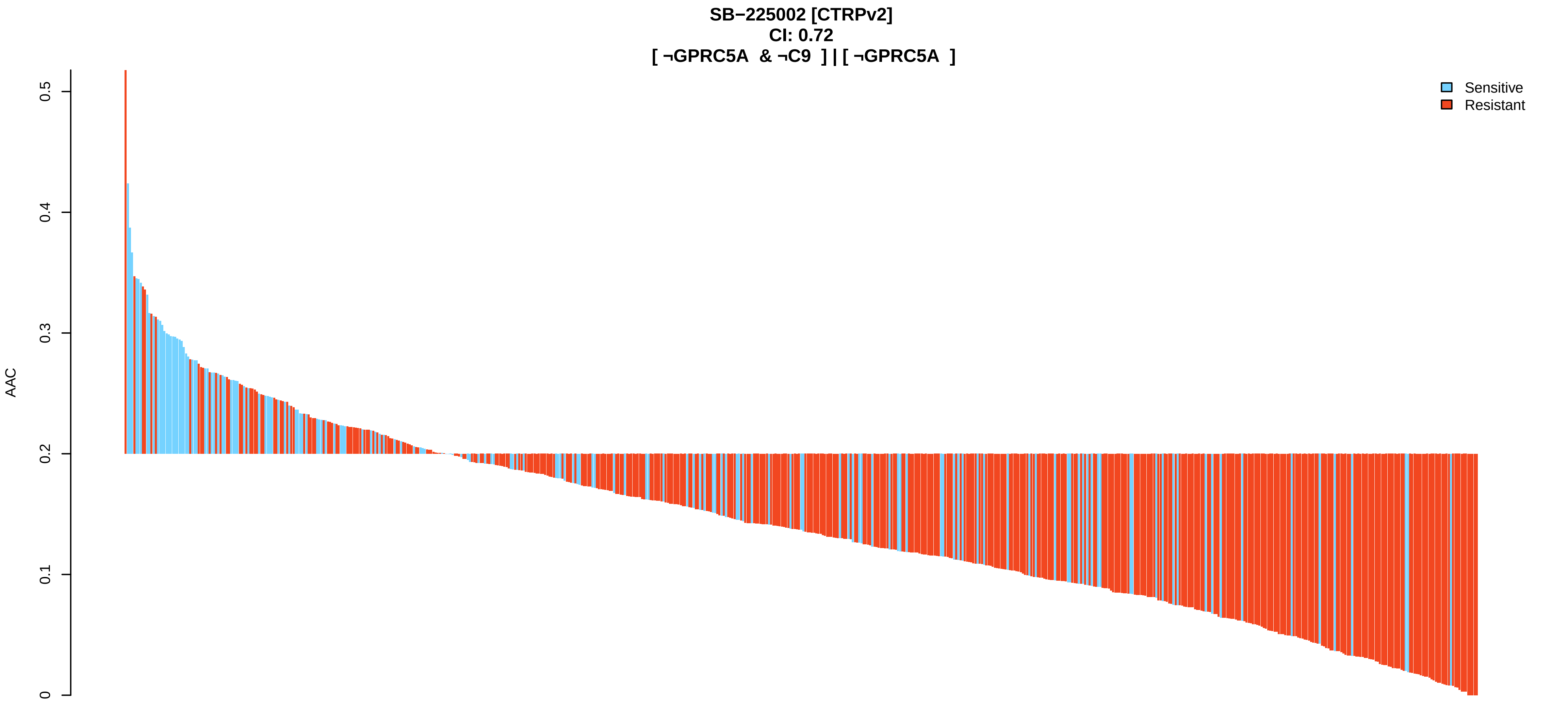
SB−225002 [CTRPv2]
CI: 0.72
[ ¬GPRC5A & ¬C9 ] | [ ¬GPRC5A ]
Sensitive
Resistant
AAC
0
0.1
0.2
0.3
0.4
0.5

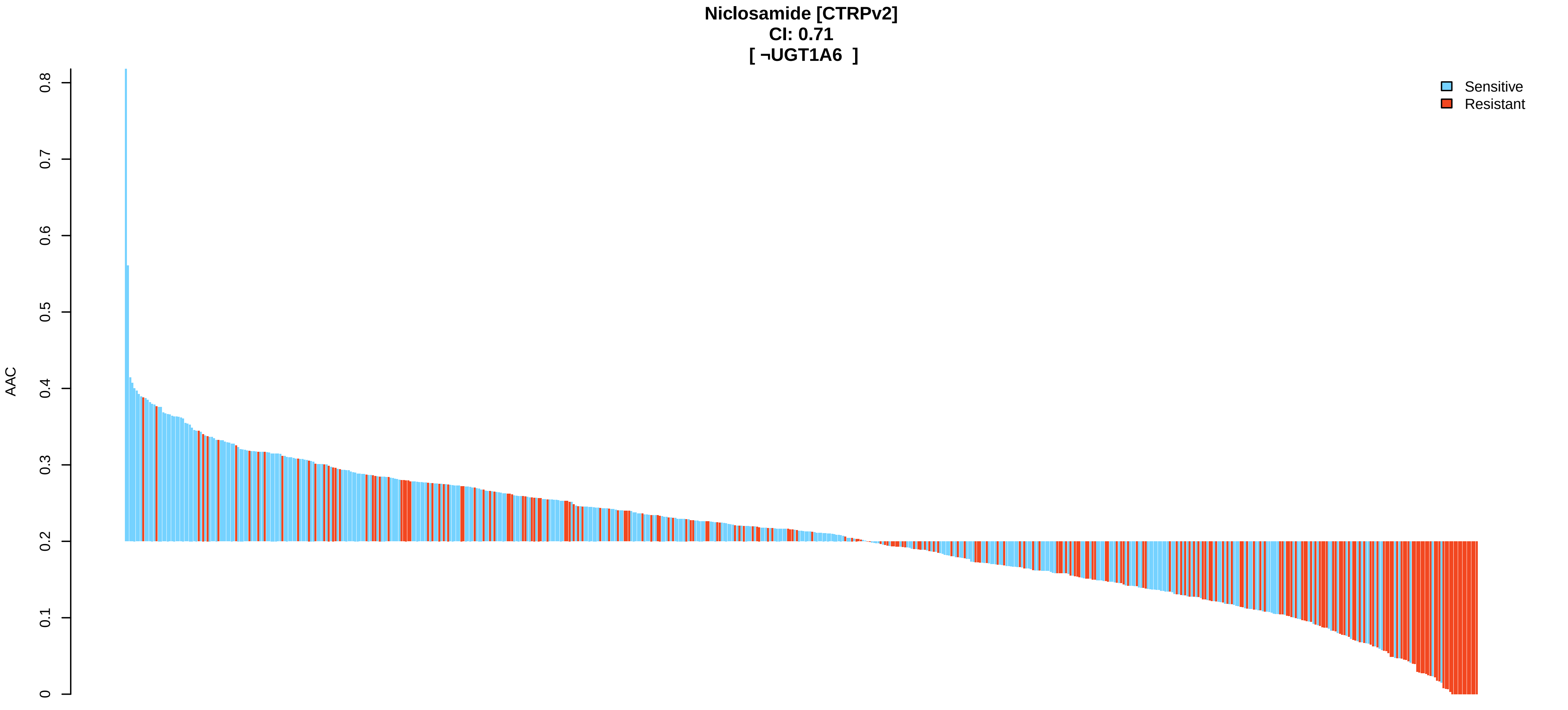
Niclosamide [CTRPv2]
CI: 0.71
[ ¬UGT1A6 ]
Sensitive
Resistant
AAC
0
0.1
0.2
0.3
0.4
0.5
0.6
0.7
0.8

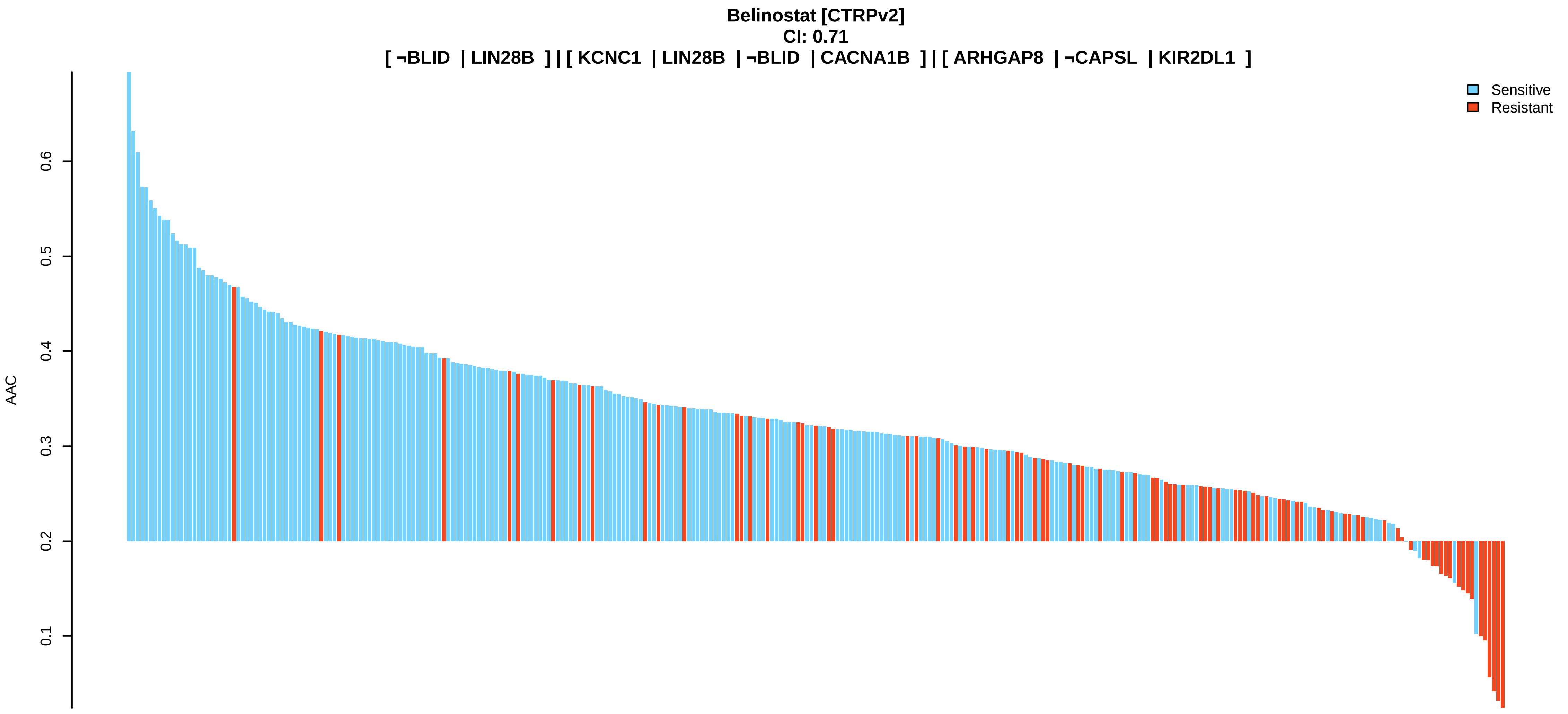
Belinostat [CTRPv2]
CI: 0.71
[ ¬BLID | LIN28B ] | [ KCNC1 | LIN28B | ¬BLID | CACNA1B ] | [ ARHGAP8 | ¬CAPSL | KIR2DL1 ]
AAC
0.1
0.2
0.3
0.4
0.5
0.6
Sensitive
Resistant

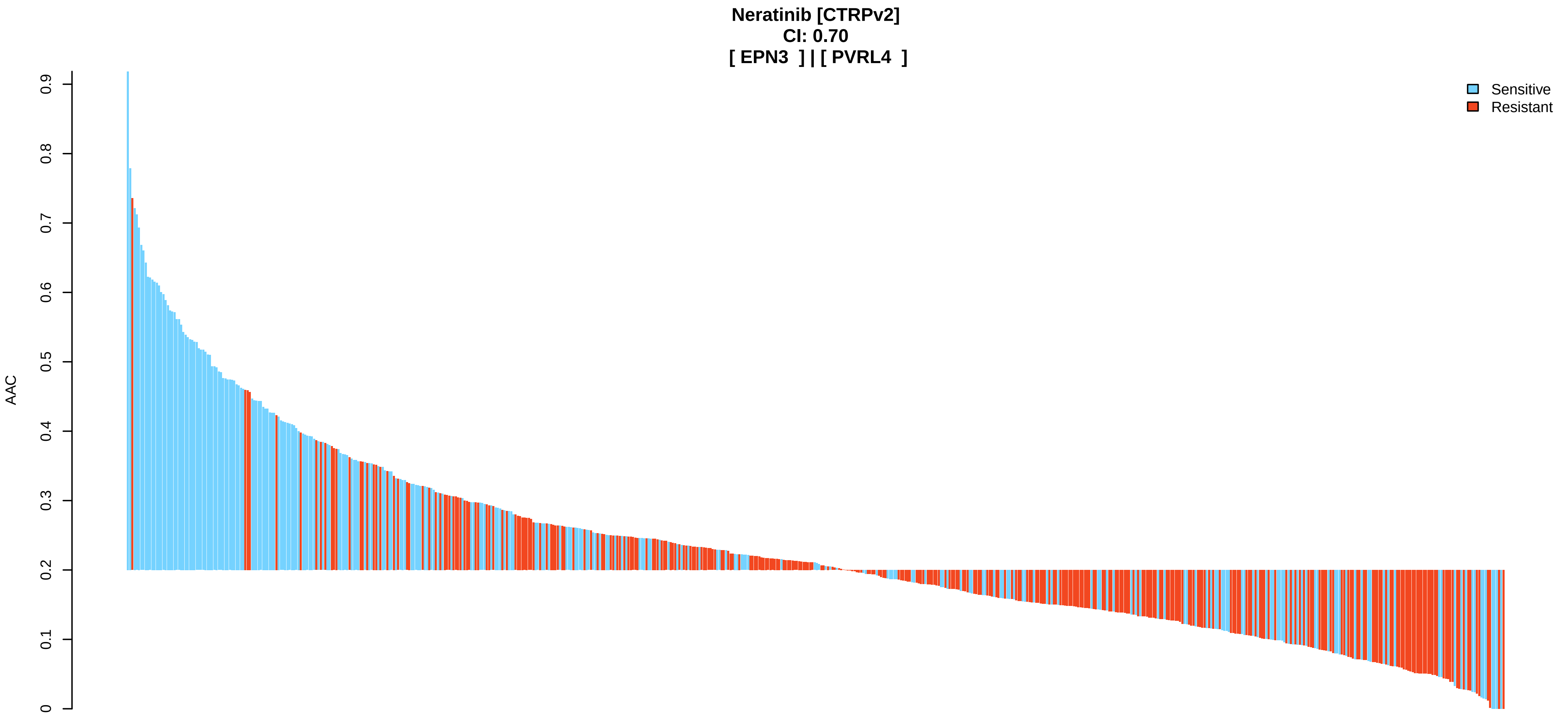
Neratinib [CTRPv2]
CI: 0.70
[ EPN3 ] | [ PVRL4 ]
AAC
0
0.1
0.2
0.3
0.4
0.5
0.6
0.7
0.8
0.9
Sensitive
Resistant

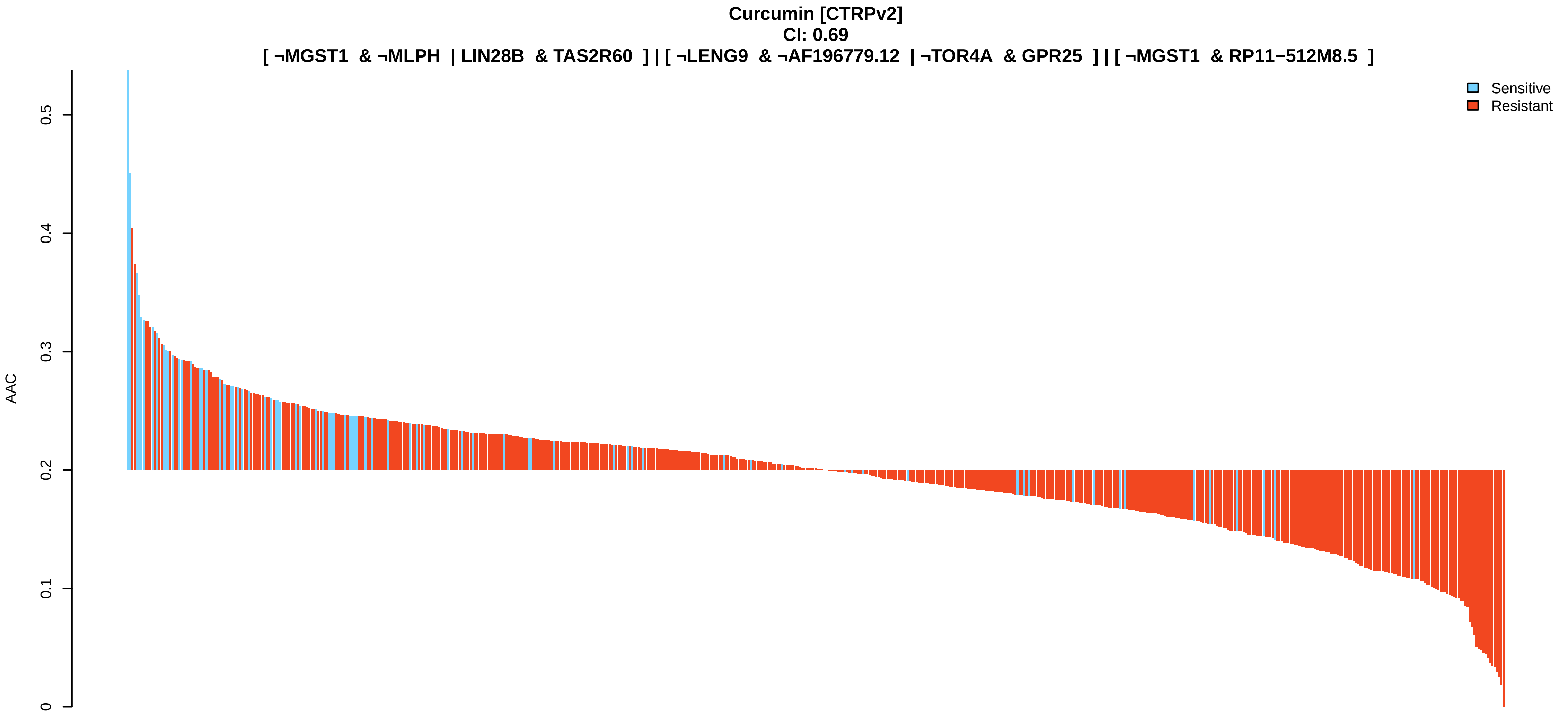
Curcumin [CTRPv2]
CI: 0.69
[ ¬MGST1 & ¬MLPH | LIN28B & TAS2R60 ] | [ ¬LENG9 & ¬AF196779.12 | ¬TOR4A & GPR25 ] | [ ¬MGST1 & RP11−512M8.5 ]
AAC
0
0.1
0.2
0.3
0.4
0.5
Sensitive
Resistant

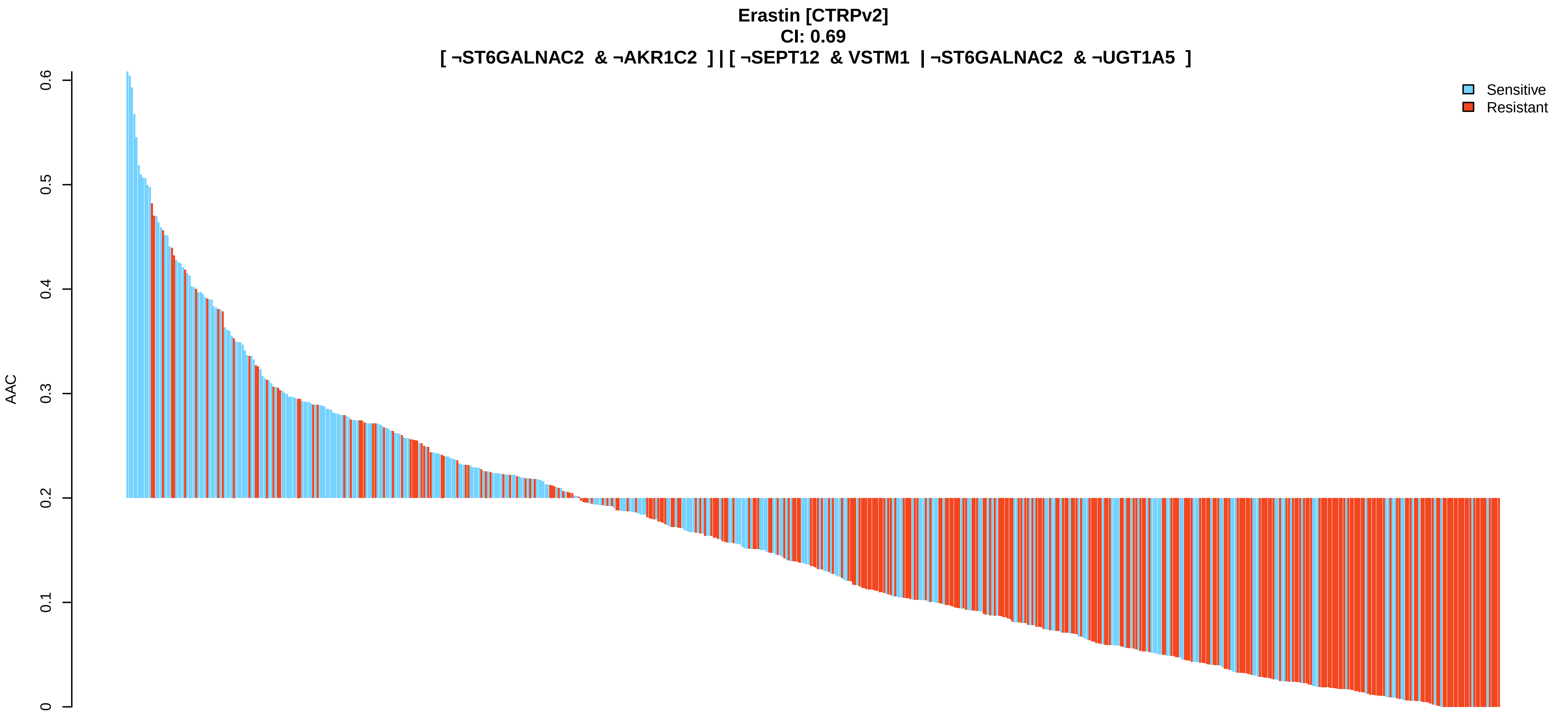

Erastin [CTRPv2]
CI: 0.69
[ ¬ST6GALNAC2 & ¬AKR1C2 ] | [ ¬SEPT12 & VSTM1 | ¬ST6GALNAC2 & ¬UGT1A5 ]
AAC
0
0.1
0.2
0.3
0.4
0.5
0.6
Sensitive
Resistant


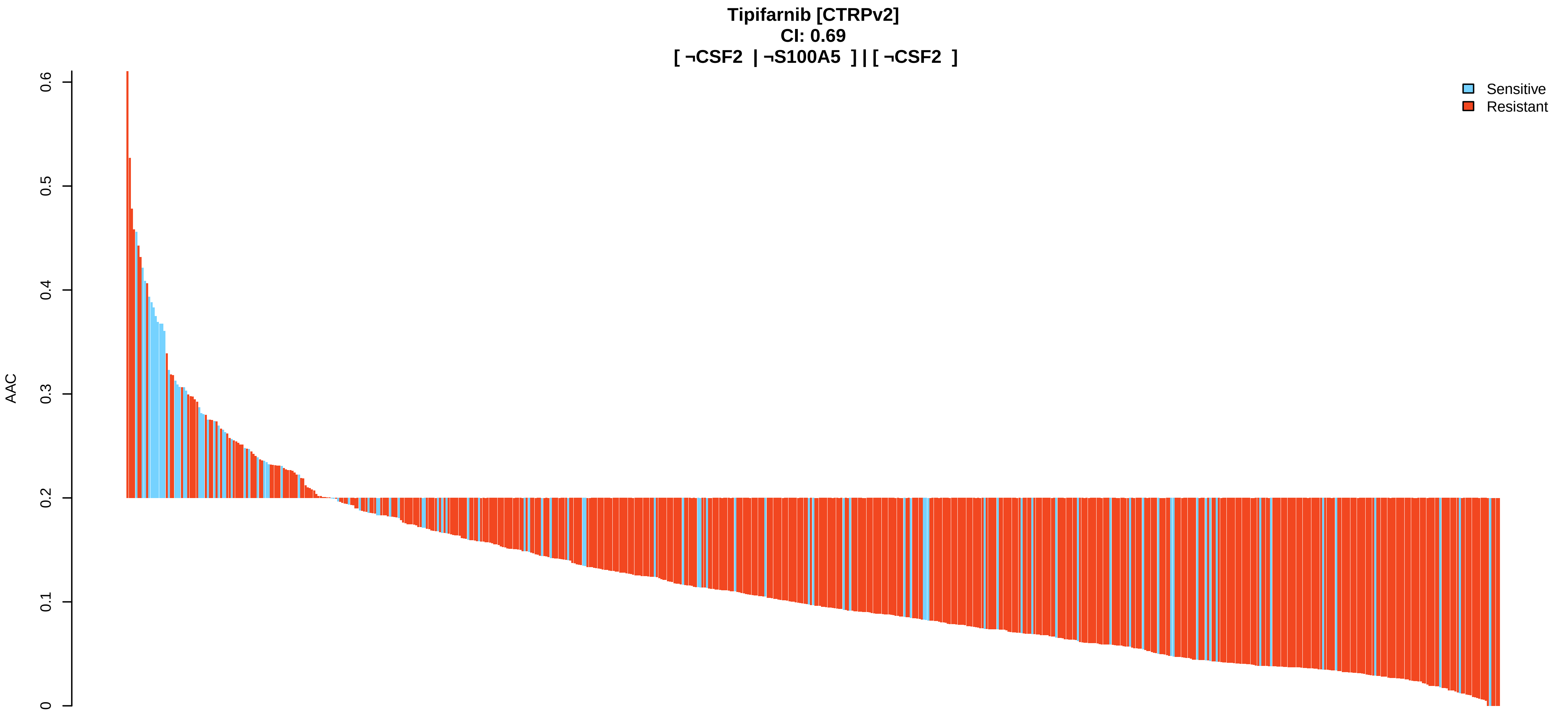

Tipifarnib [CTRPv2]
CI: 0.69
[ ¬CSF2 | ¬S100A5 ] | [ ¬CSF2 ]
AAC
0
0.1
0.2
0.3
0.4
0.5
0.6
Sensitive
Resistant


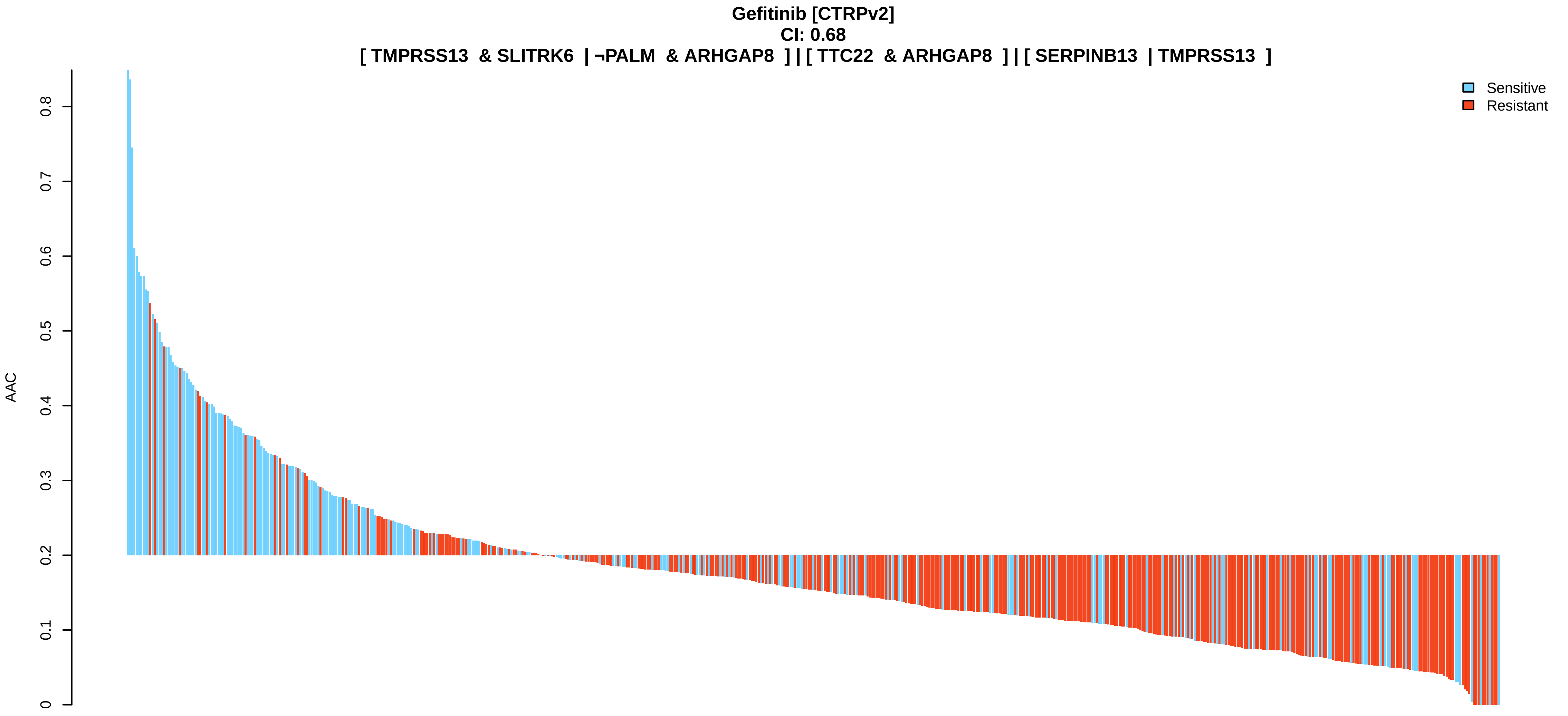

Gefitinib [CTRPv2]
CI: 0.68
[ TMPRSS13 & SLITRK6 | ¬PALM & ARHGAP8 ] | [ TTC22 & ARHGAP8 ] | [ SERPINB13 | TMPRSS13 ]
AAC
0
0.1
0.2
0.3
0.4
0.5
0.6
0.7
0.8
Sensitive
Resistant

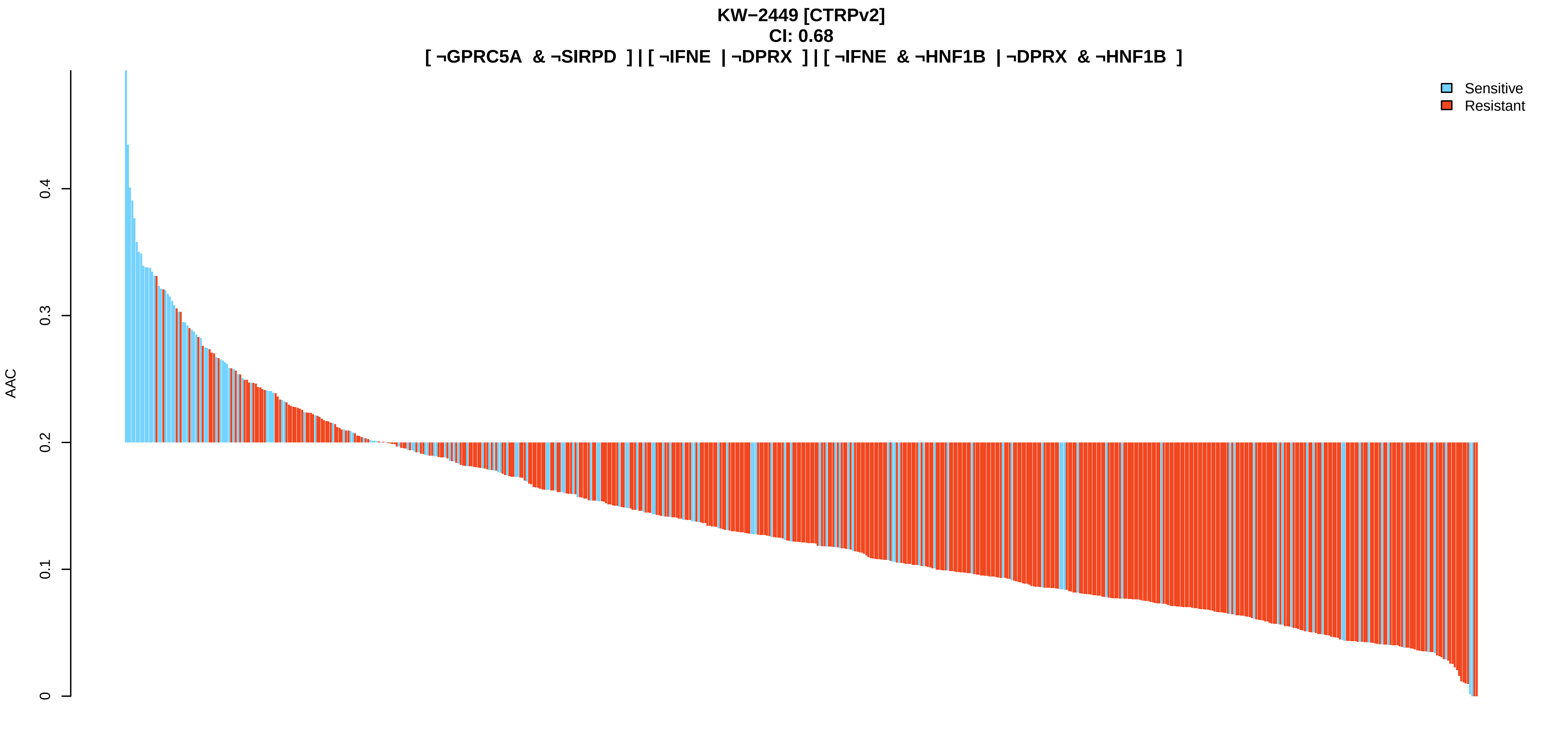
KW−2449 [CTRPv2]
CI: 0.68
[ ¬GPRC5A & ¬SIRPD ] | [ ¬IFNE | ¬DPRX ] | [ ¬IFNE & ¬HNF1B | ¬DPRX & ¬HNF1B ]
Sensitive
Resistant
AAC
0
0.1
0.2
0.3
0.4

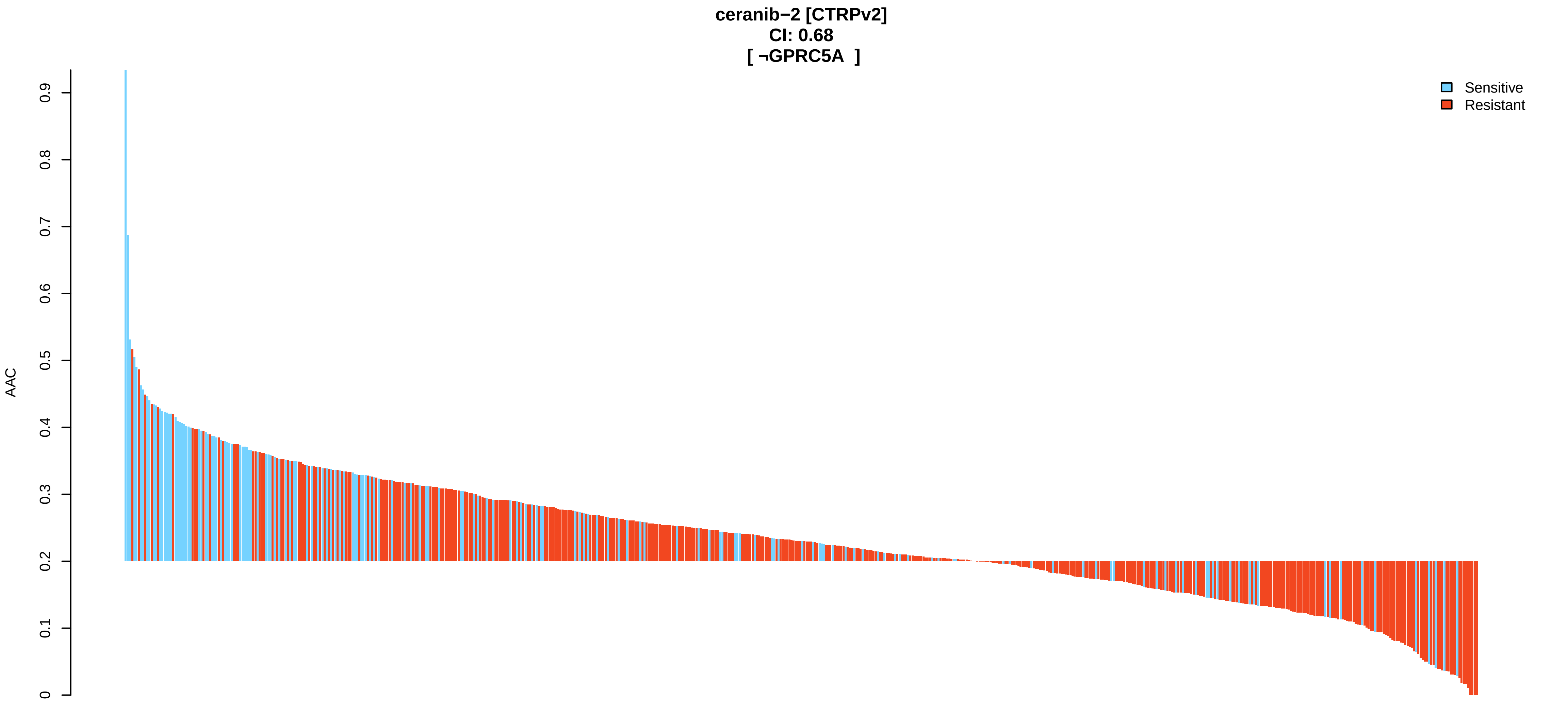
ceranib−2 [CTRPv2]
CI: 0.68
[ ¬GPRC5A ]
Sensitive
Resistant
AAC
0
0.1
0.2
0.3
0.4
0.5
0.6
0.7
0.8
0.9

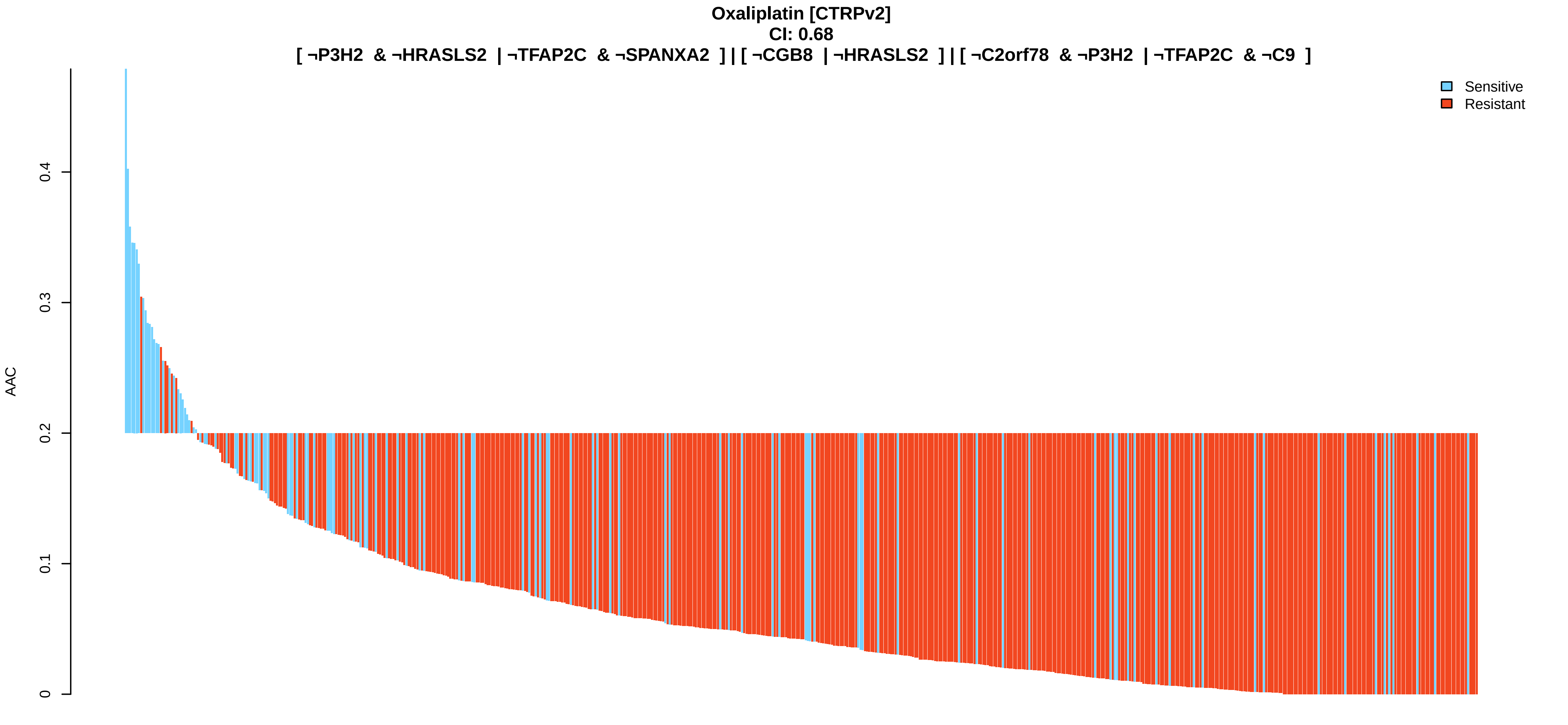
Oxaliplatin [CTRPv2]
CI: 0.68
[ ¬P3H2 & ¬HRASLS2 | ¬TFAP2C & ¬SPANXA2 ] | [ ¬CGB8 | ¬HRASLS2 ] | [ ¬C2orf78 & ¬P3H2 | ¬TFAP2C & ¬C9 ]
Sensitive
Resistant
AAC
0
0.1
0.2
0.3
0.4

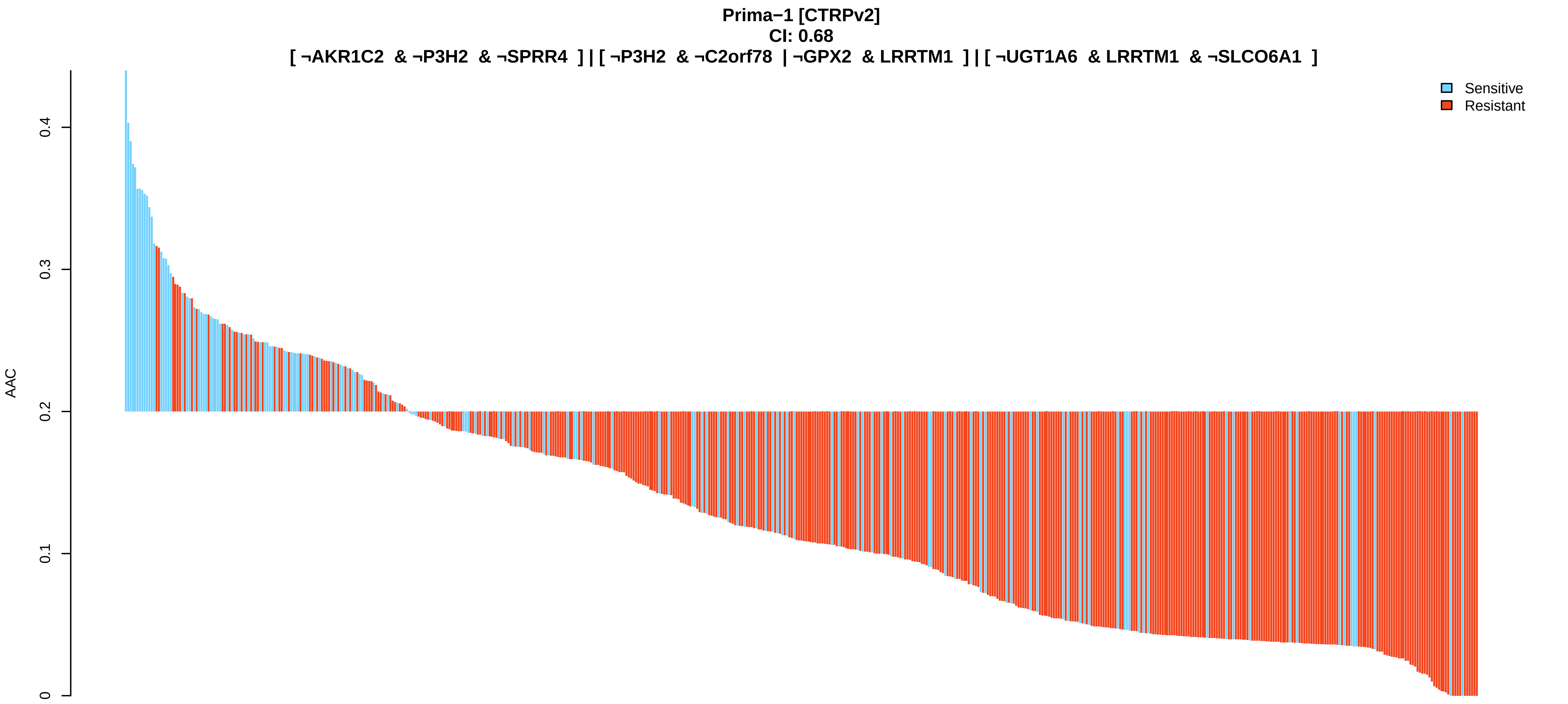

Prima−1 [CTRPv2]
CI: 0.68
[ ¬AKR1C2 & ¬P3H2 & ¬SPRR4 ] | [ ¬P3H2 & ¬C2orf78 | ¬GPX2 & LRRTM1 ] | [ ¬UGT1A6 & LRRTM1 & ¬SLCO6A1 ]
AAC
0
0.1
0.2
0.3
0.4
Sensitive
Resistant


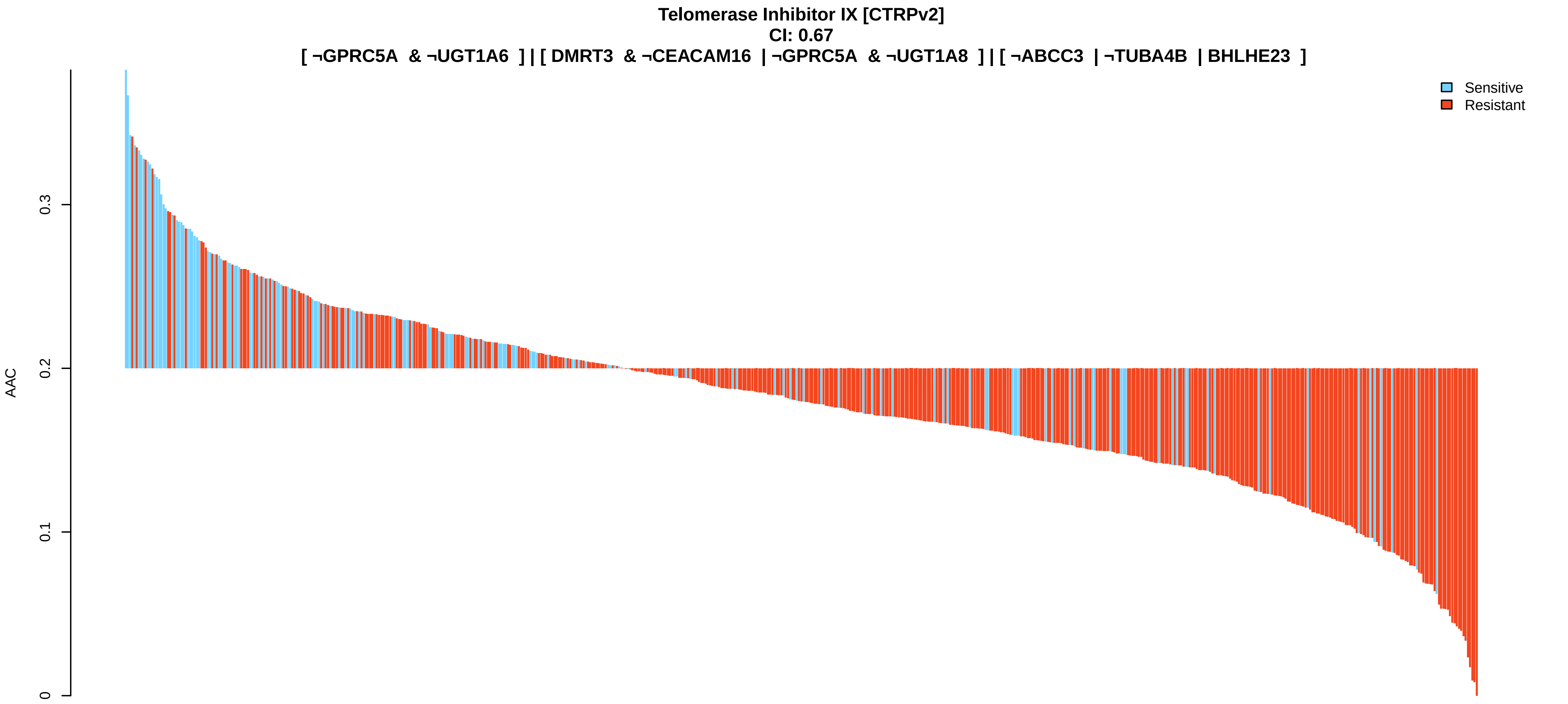

Telomerase Inhibitor IX [CTRPv2]
CI: 0.67
[ ¬GPRC5A & ¬UGT1A6 ] | [ DMRT3 & ¬CEACAM16 | ¬GPRC5A & ¬UGT1A8 ] | [ ¬ABCC3 | ¬TUBA4B | BHLHE23 ]
AAC
0
0.1
0.2
0.3
Sensitive
Resistant


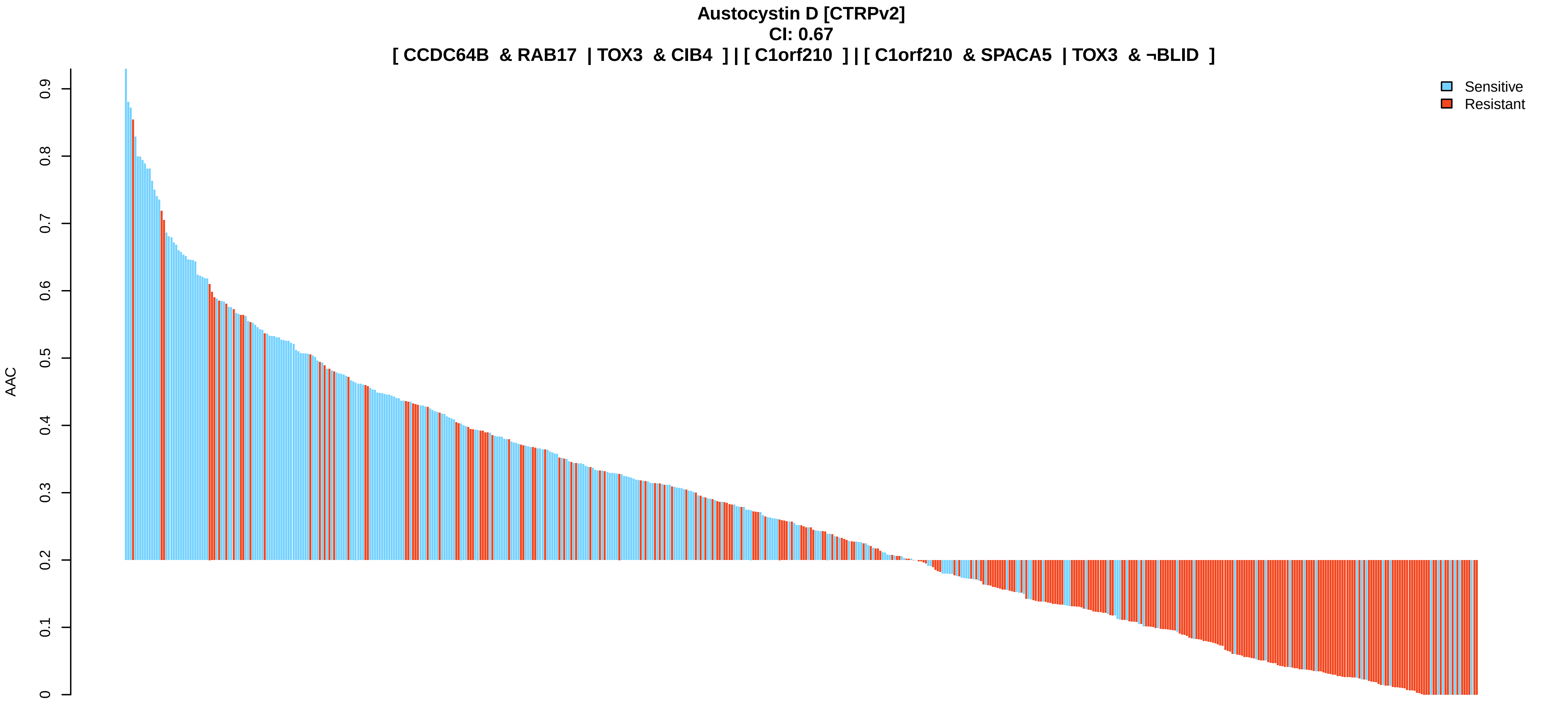

Austocystin D [CTRPv2]
CI: 0.67
[ CCDC64B & RAB17 | TOX3 & CIB4 ] | [ C1orf210 ] | [ C1orf210 & SPACA5 | TOX3 & ¬BLID ]
AAC
0
0.1
0.2
0.3
0.4
0.5
0.6
0.7
0.8
0.9
Sensitive
Resistant

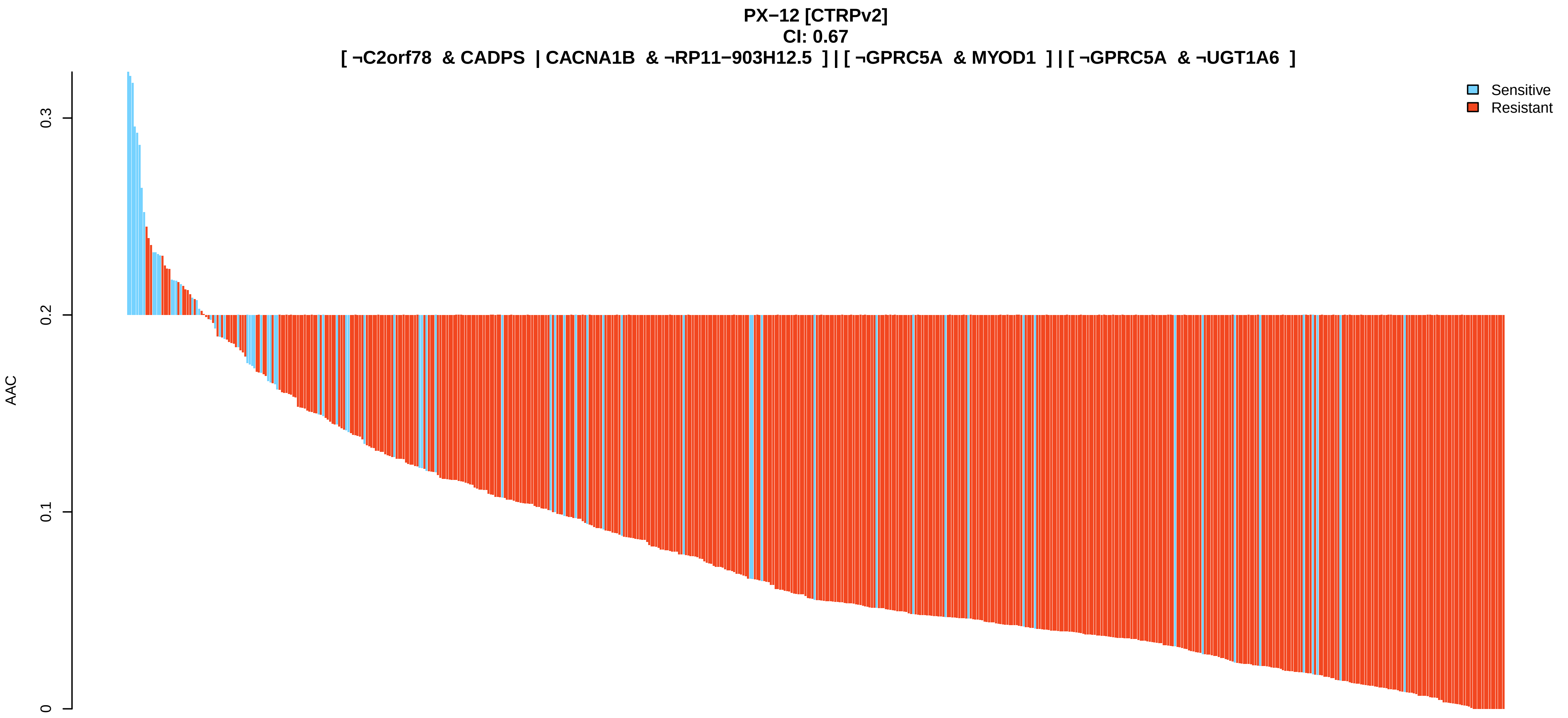
PX−12 [CTRPv2]
CI: 0.67
[ ¬C2orf78 & CADPS | CACNA1B & ¬RP11−903H12.5 ] | [ ¬GPRC5A & MYOD1 ] | [ ¬GPRC5A & ¬UGT1A6 ]
AAC
0
0.1
0.2
0.3
Sensitive
Resistant

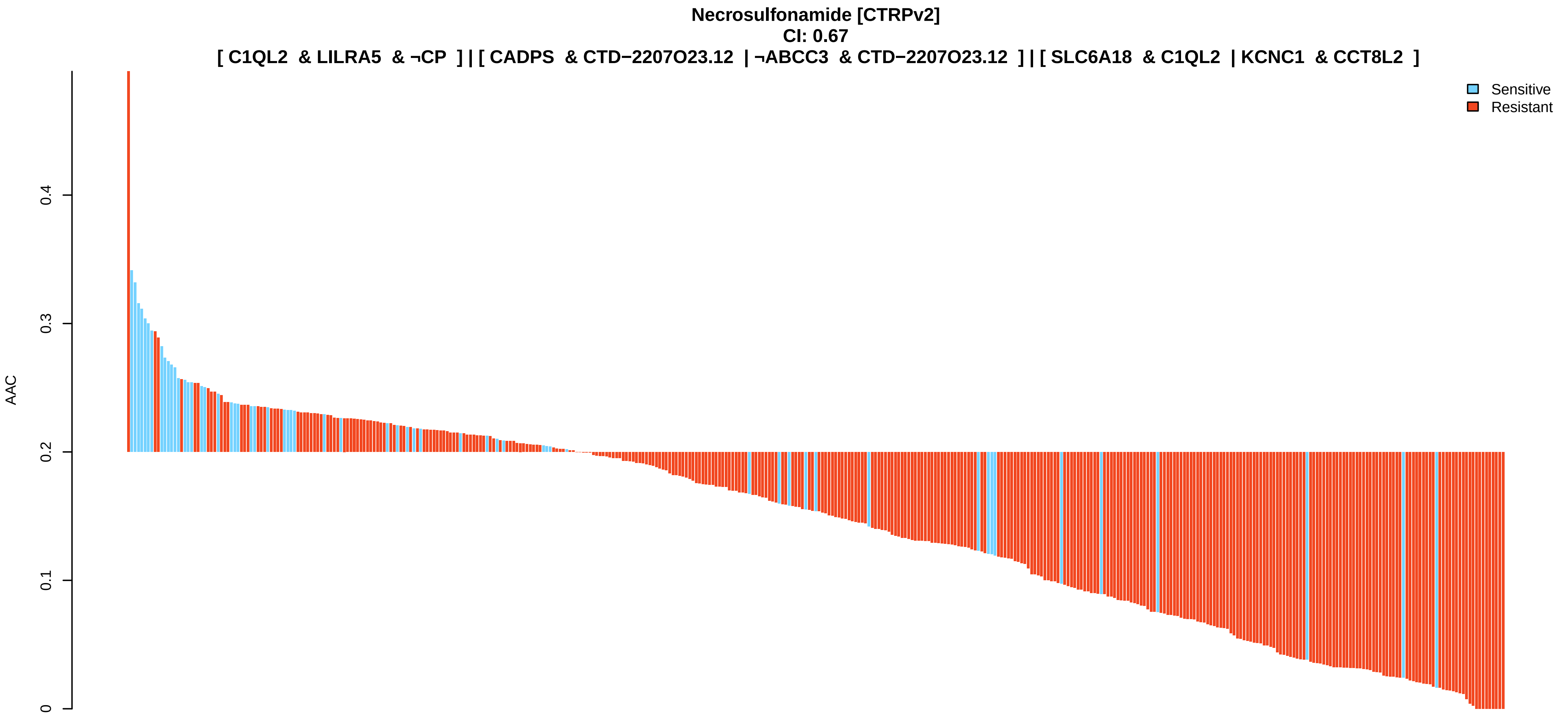
Necrosulfonamide [CTRPv2]
CI: 0.67
[ C1QL2 & LILRA5 & ¬CP ] | [ CADPS & CTD−2207O23.12 | ¬ABCC3 & CTD−2207O23.12 ] | [ SLC6A18 & C1QL2 | KCNC1 & CCT8L2 ]
AAC
0
0.1
0.2
0.3
0.4
Sensitive
Resistant

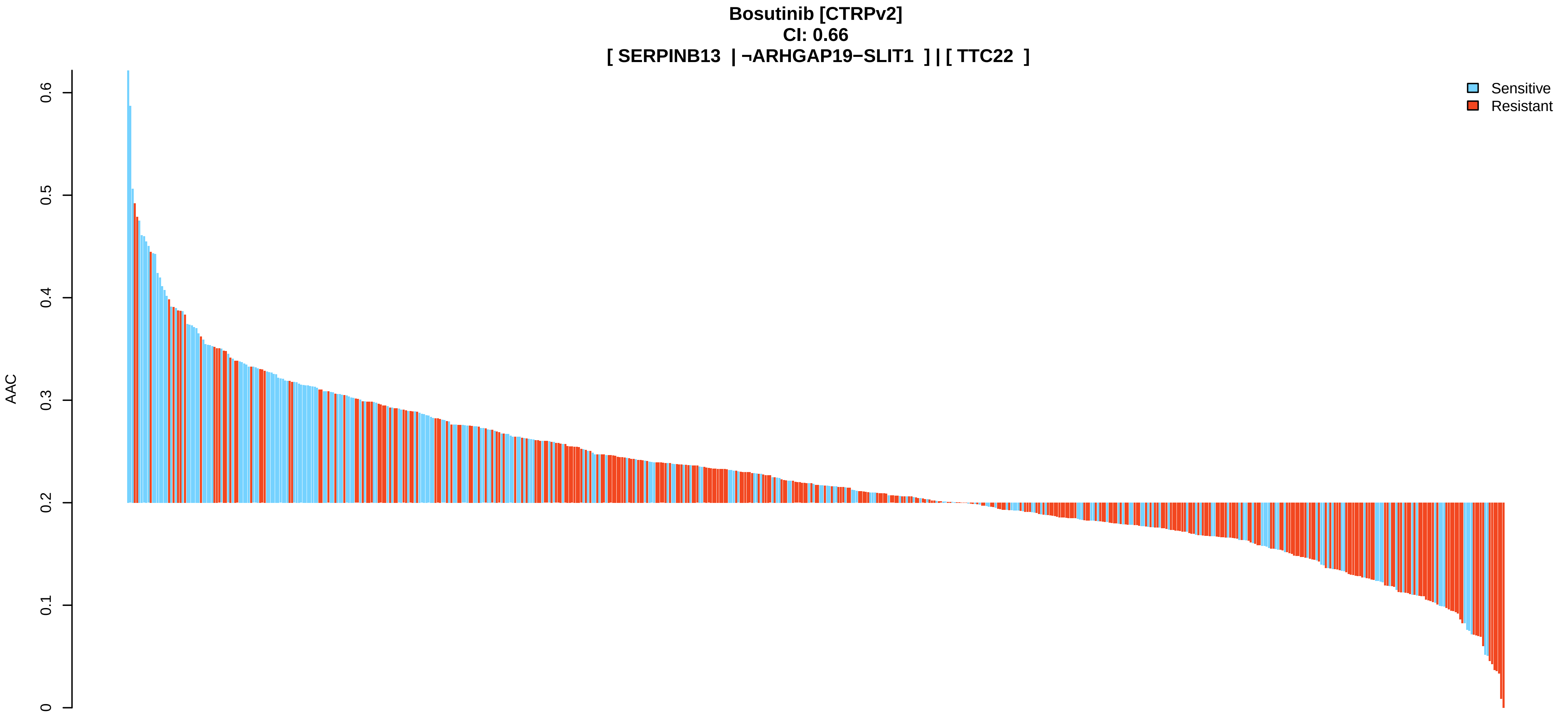
Bosutinib [CTRPv2]
CI: 0.66
[ SERPINB13 | ¬ARHGAP19−SLIT1 ] | [ TTC22 ]
AAC
0
0.1
0.2
0.3
0.4
0.5
0.6
Sensitive
Resistant

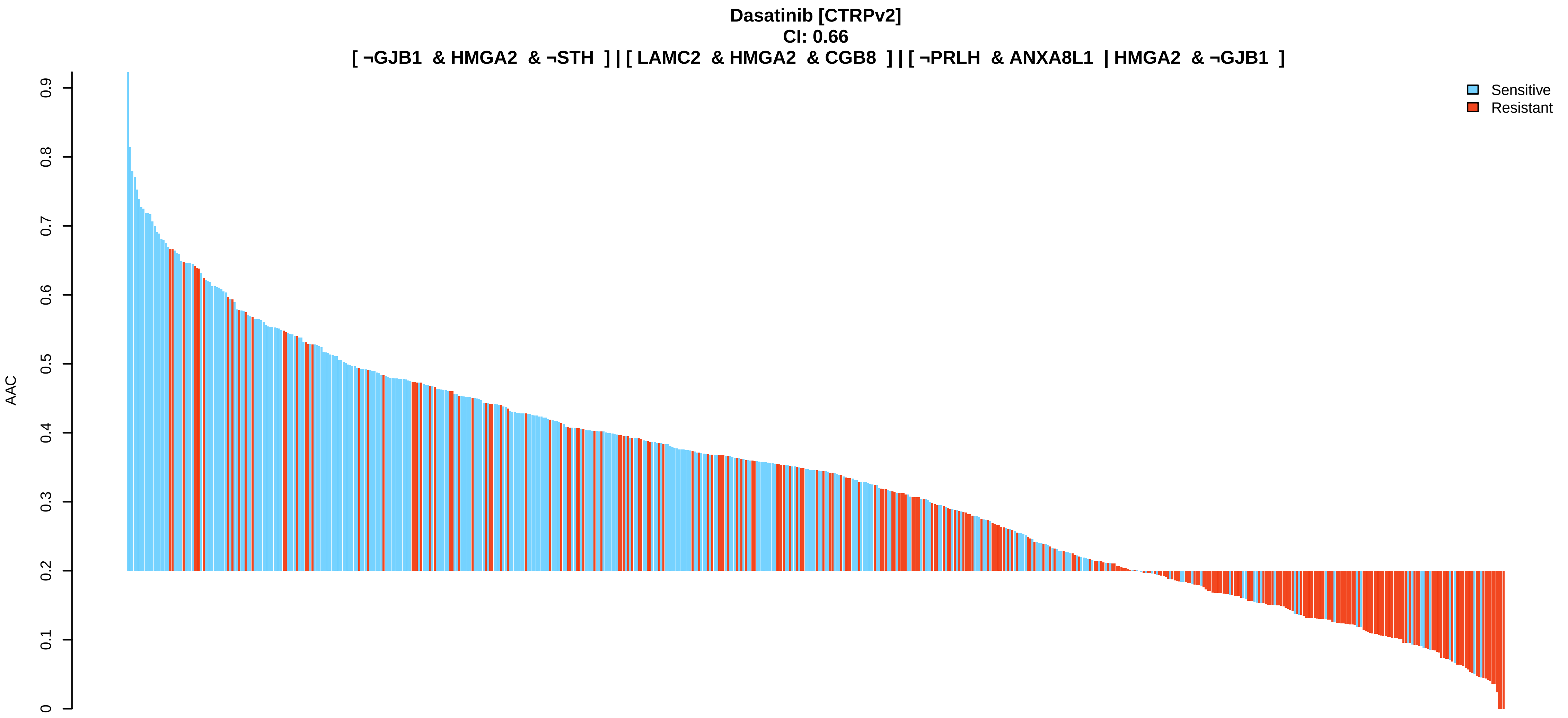
Dasatinib [CTRPv2]
CI: 0.66
[ ¬GJB1 & HMGA2 & ¬STH ] | [ LAMC2 & HMGA2 & CGB8 ] | [ ¬PRLH & ANXA8L1 | HMGA2 & ¬GJB1 ]
AAC
0
0.1
0.2
0.3
0.4
0.5
0.6
0.7
0.8
0.9
Sensitive
Resistant

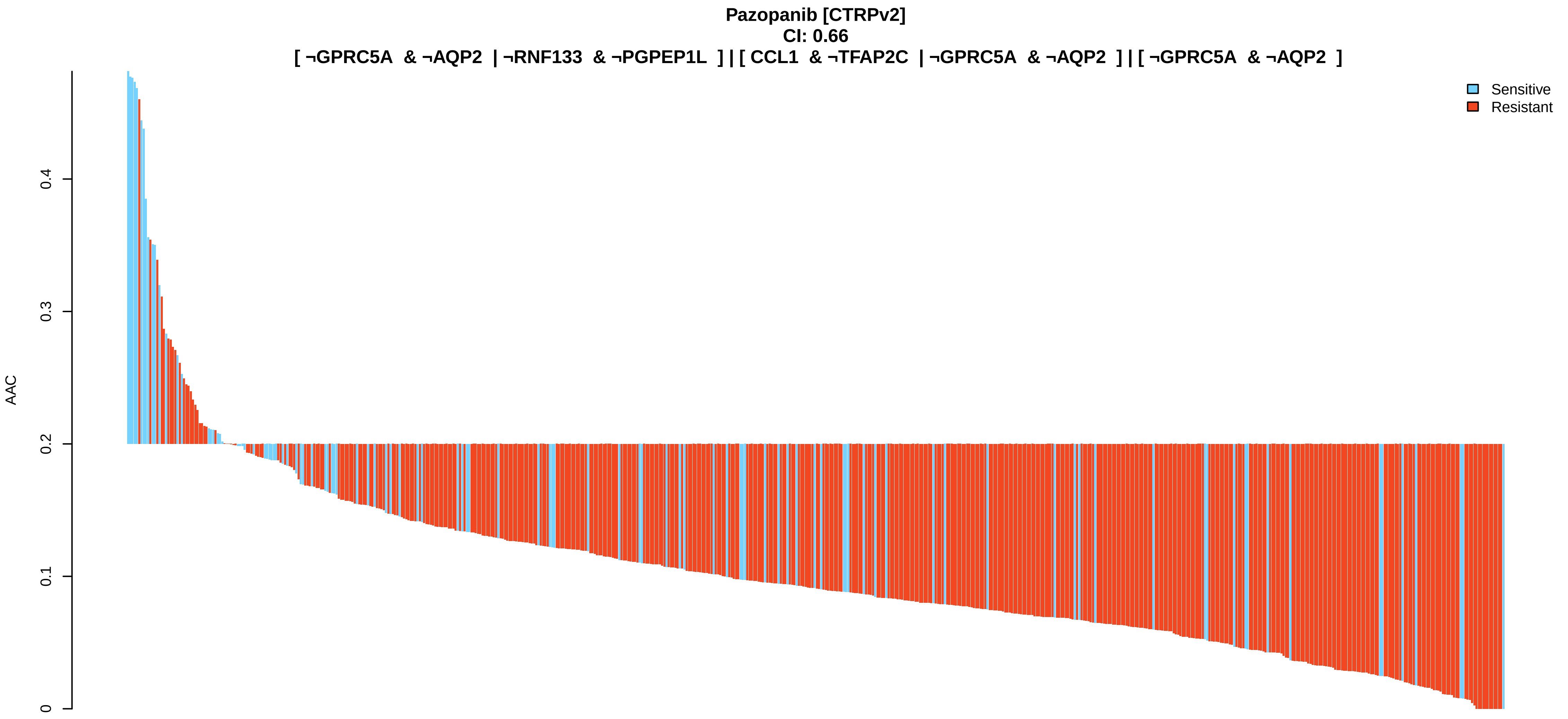
Pazopanib [CTRPv2]
CI: 0.66
[ ¬GPRC5A & ¬AQP2 | ¬RNF133 & ¬PGPEP1L ] | [ CCL1 & ¬TFAP2C | ¬GPRC5A & ¬AQP2 ] | [ ¬GPRC5A & ¬AQP2 ]
AAC
0
0.1
0.2
0.3
0.4
Sensitive
Resistant

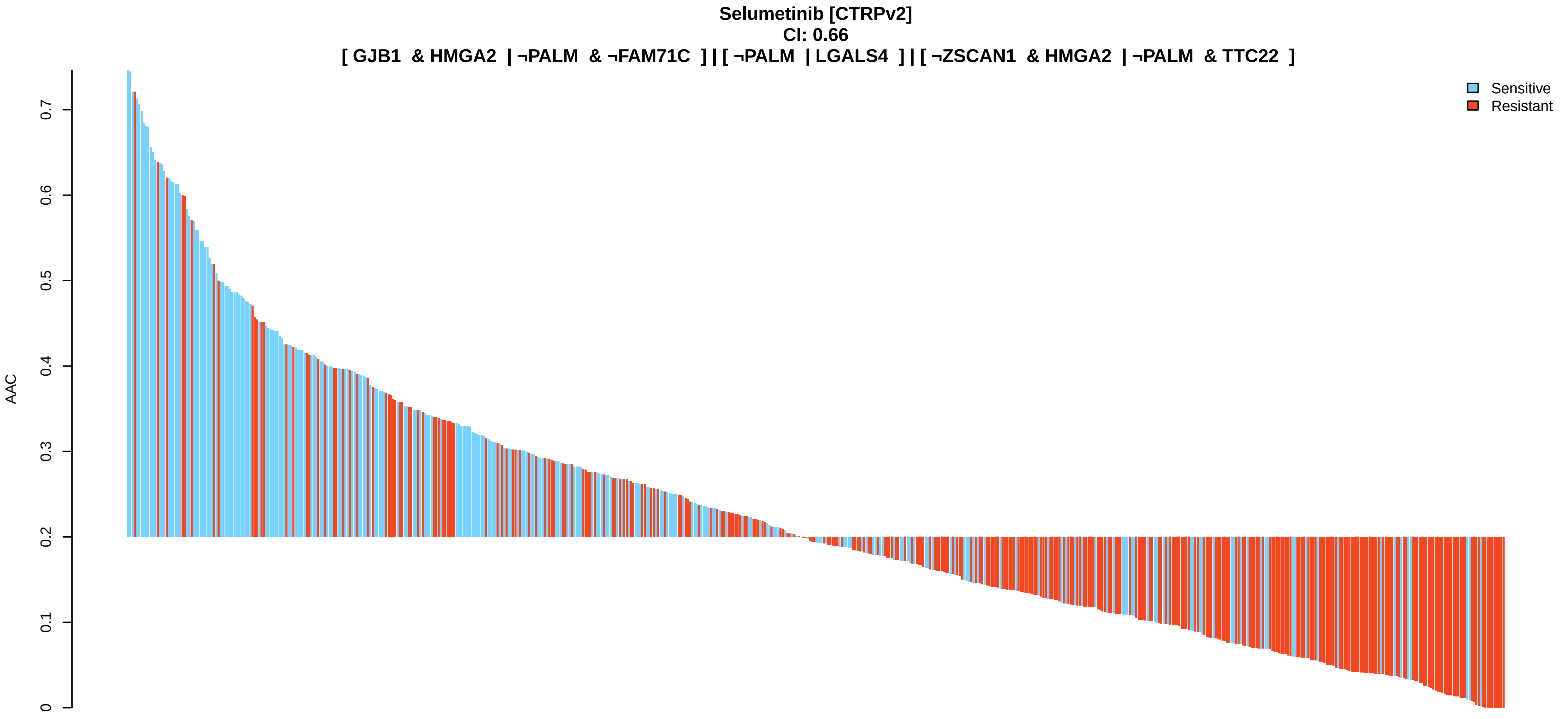
Selumetinib [CTRPv2]
CI: 0.66
[ GJB1 & HMGA2 | ¬PALM & ¬FAM71C ] | [ ¬PALM | LGALS4 ] | [ ¬ZSCAN1 & HMGA2 | ¬PALM & TTC22 ]
AAC
0
0.1
0.2
0.3
0.4
0.5
0.6
0.7
Sensitive
Resistant

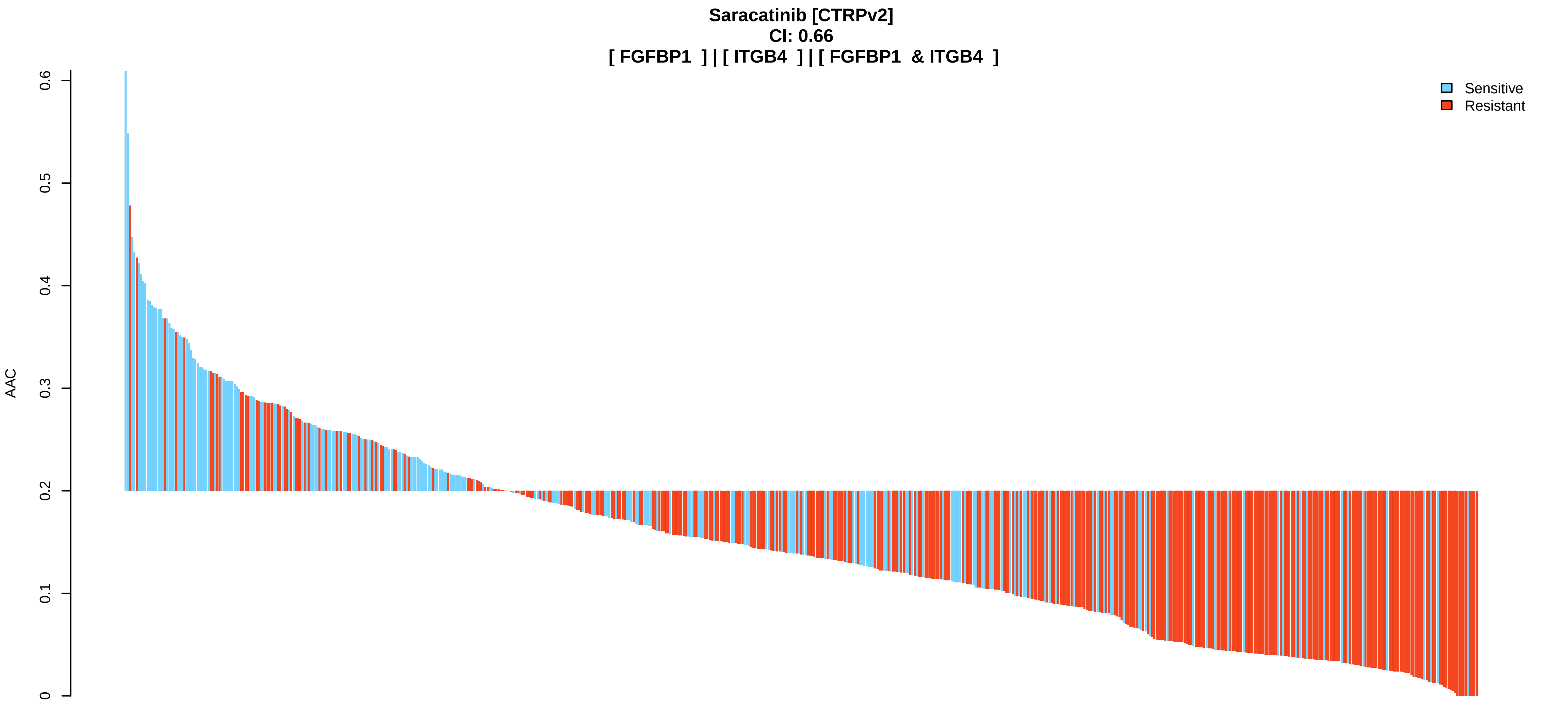
Saracatinib [CTRPv2]
CI: 0.66
[ FGFBP1 ] | [ ITGB4 ] | [ FGFBP1 & ITGB4 ]
AAC
0
0.1
0.2
0.3
0.4
0.5
0.6
Sensitive
Resistant

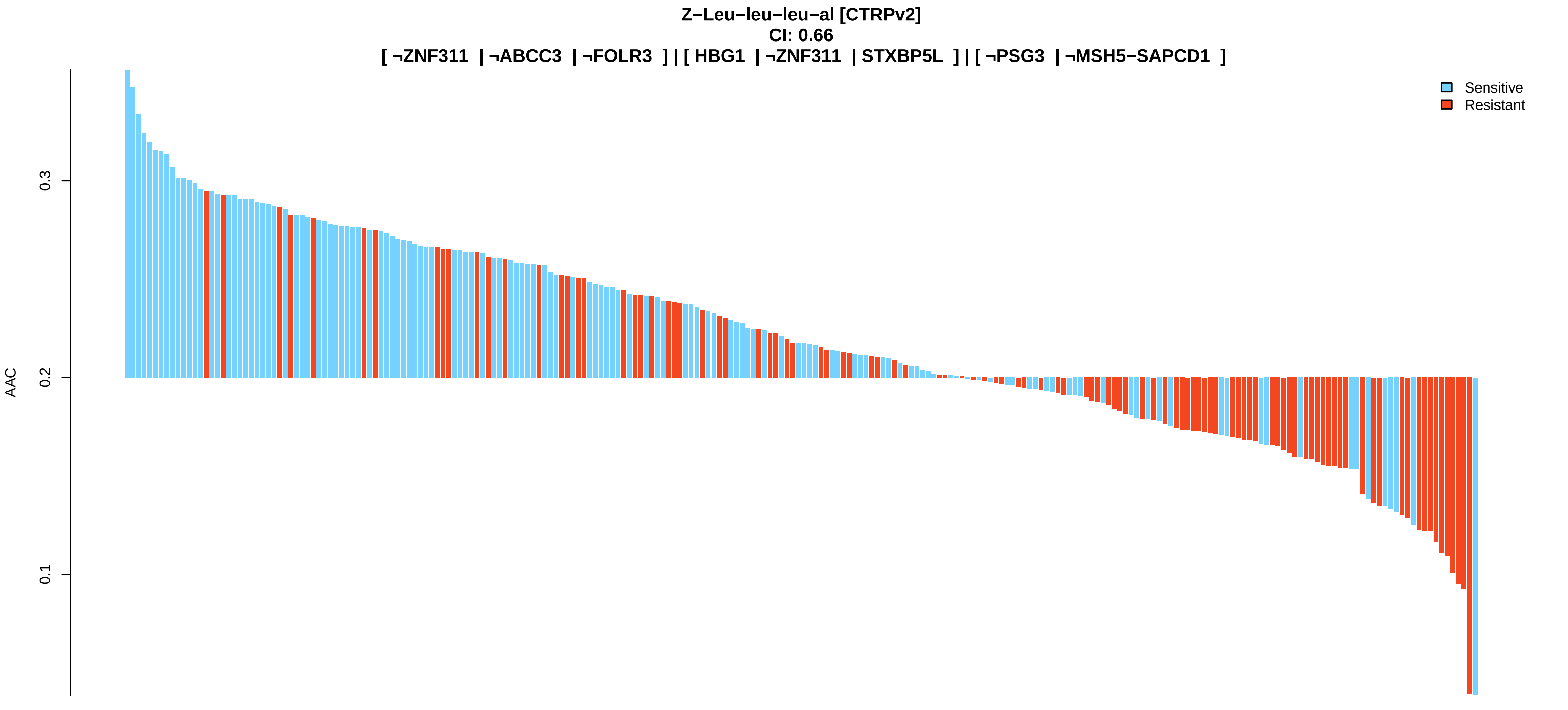
Z−Leu−leu−leu−al [CTRPv2]
CI: 0.66
[ ¬ZNF311 | ¬ABCC3 | ¬FOLR3 ] | [ HBG1 | ¬ZNF311 | STXBP5L ] | [ ¬PSG3 | ¬MSH5−SAPCD1 ]
AAC
0.1
0.2
0.3
Sensitive
Resistant

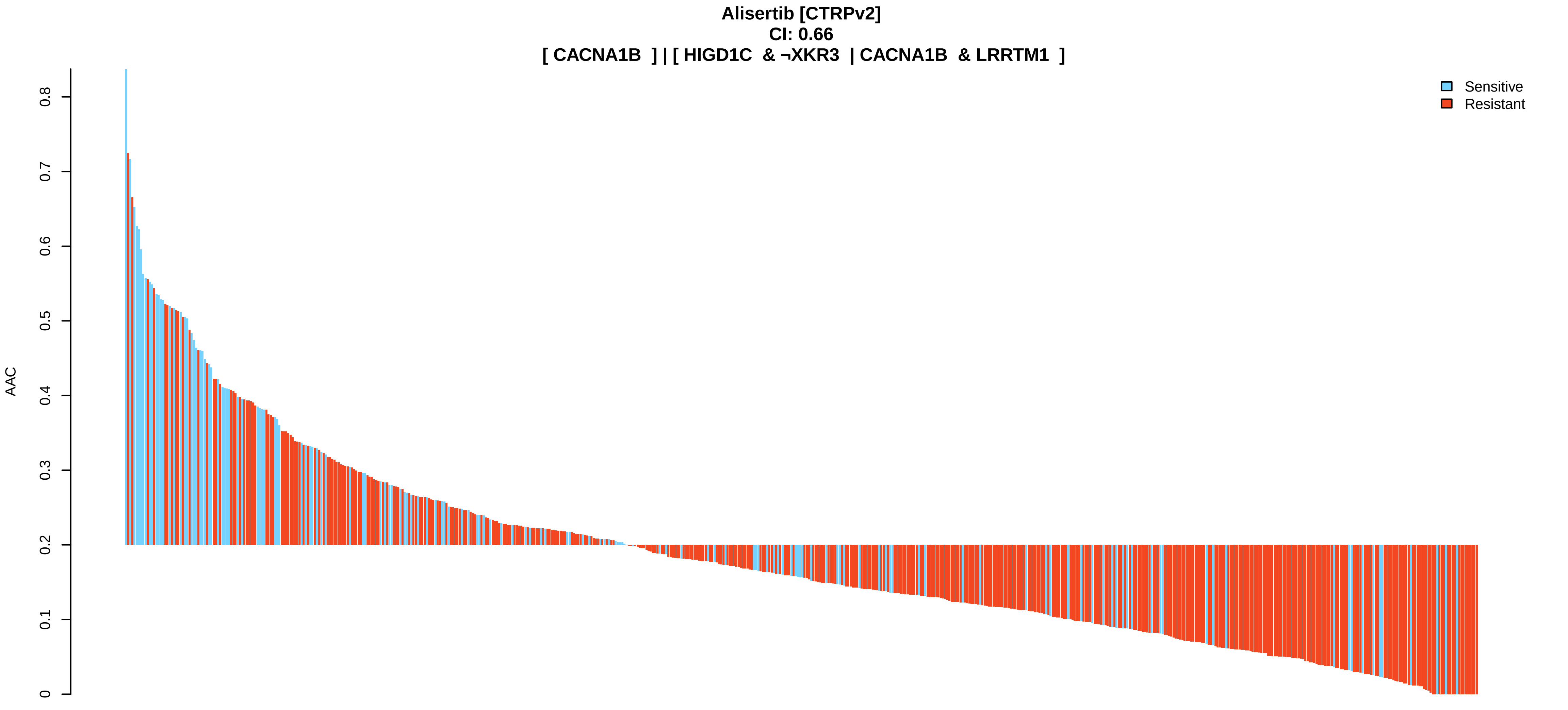
Alisertib [CTRPv2]
CI: 0.66
[ CACNA1B ] | [ HIGD1C & ¬XKR3 | CACNA1B & LRRTM1 ]
AAC
0
0.1
0.2
0.3
0.4
0.5
0.6
0.7
0.8
Sensitive
Resistant

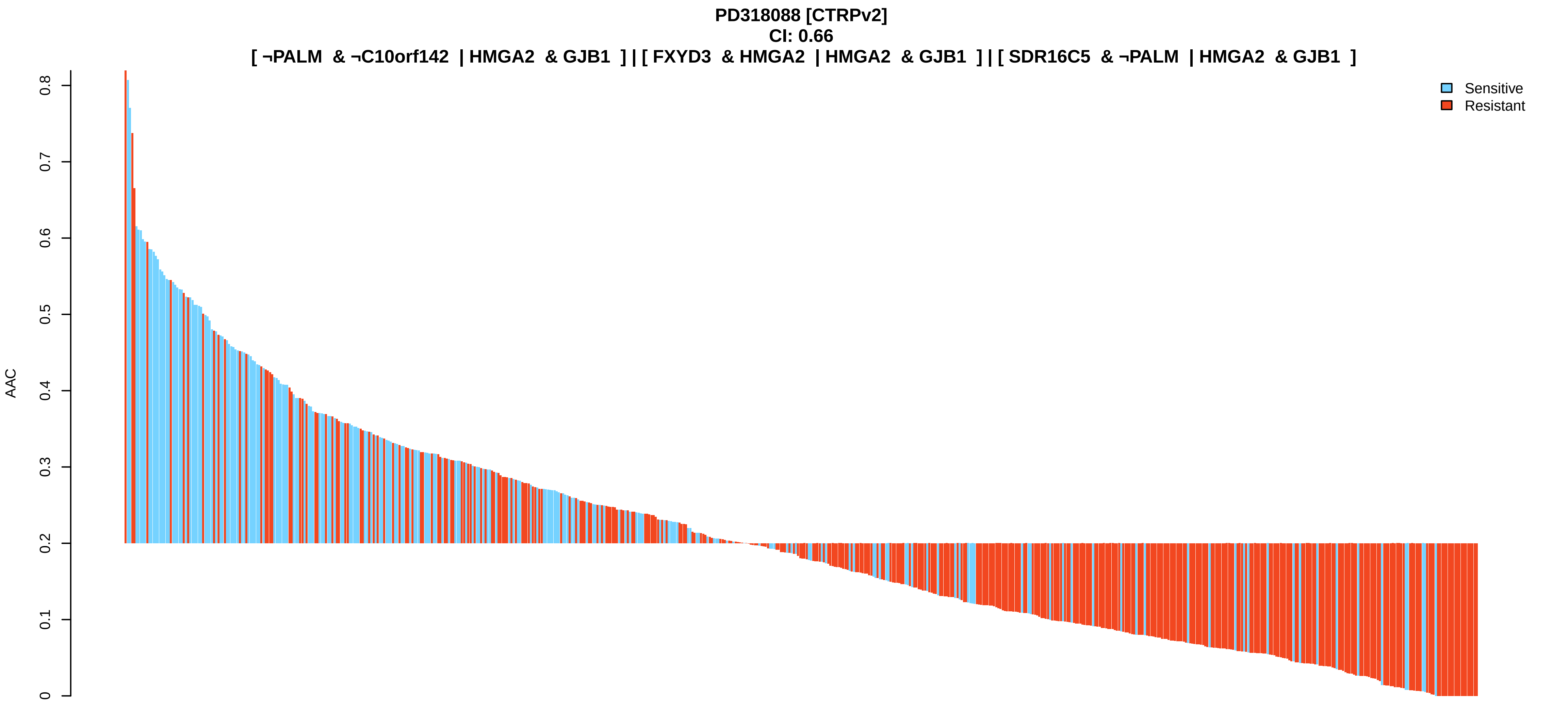
PD318088 [CTRPv2]
CI: 0.66
[ ¬PALM & ¬C10orf142 | HMGA2 & GJB1 ] | [ FXYD3 & HMGA2 | HMGA2 & GJB1 ] | [ SDR16C5 & ¬PALM | HMGA2 & GJB1 ]
AAC
0
0.1
0.2
0.3
0.4
0.5
0.6
0.7
0.8
Sensitive
Resistant

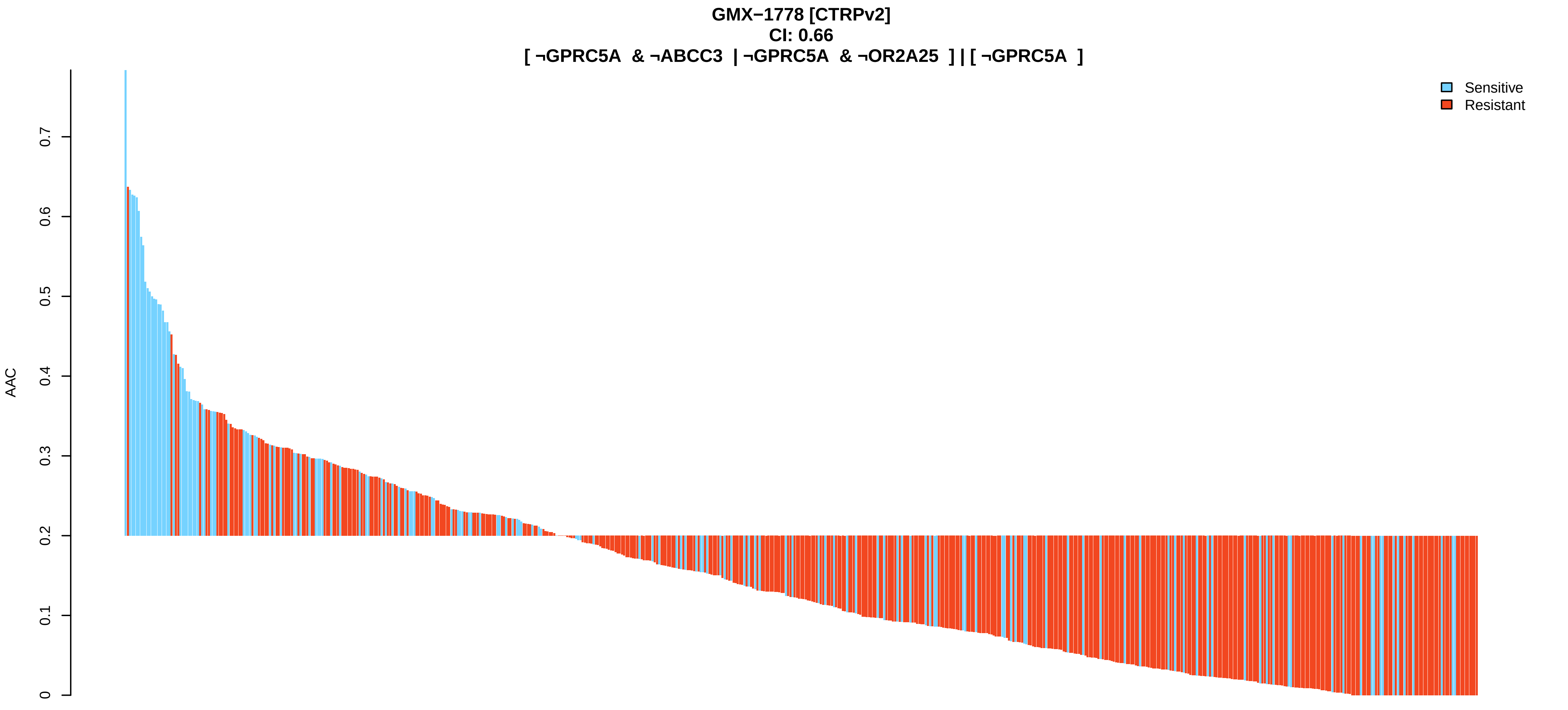
GMX−1778 [CTRPv2]
CI: 0.66
[ ¬GPRC5A & ¬ABCC3 | ¬GPRC5A & ¬OR2A25 ] | [ ¬GPRC5A ]
AAC
0
0.1
0.2
0.3
0.4
0.5
0.6
0.7
Sensitive
Resistant

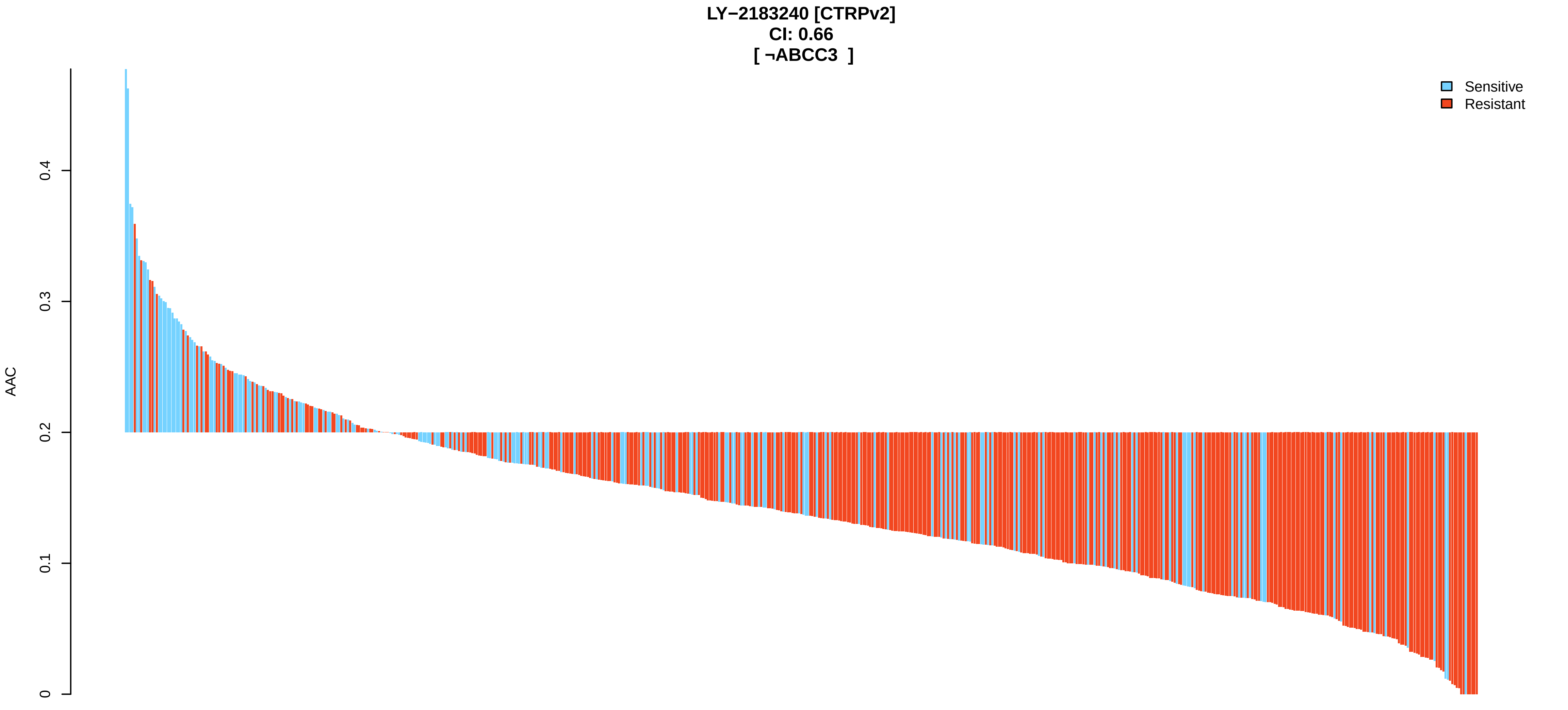
LY−2183240 [CTRPv2]
CI: 0.66
[ ¬ABCC3 ]
AAC
0
0.1
0.2
0.3
0.4
Sensitive
Resistant

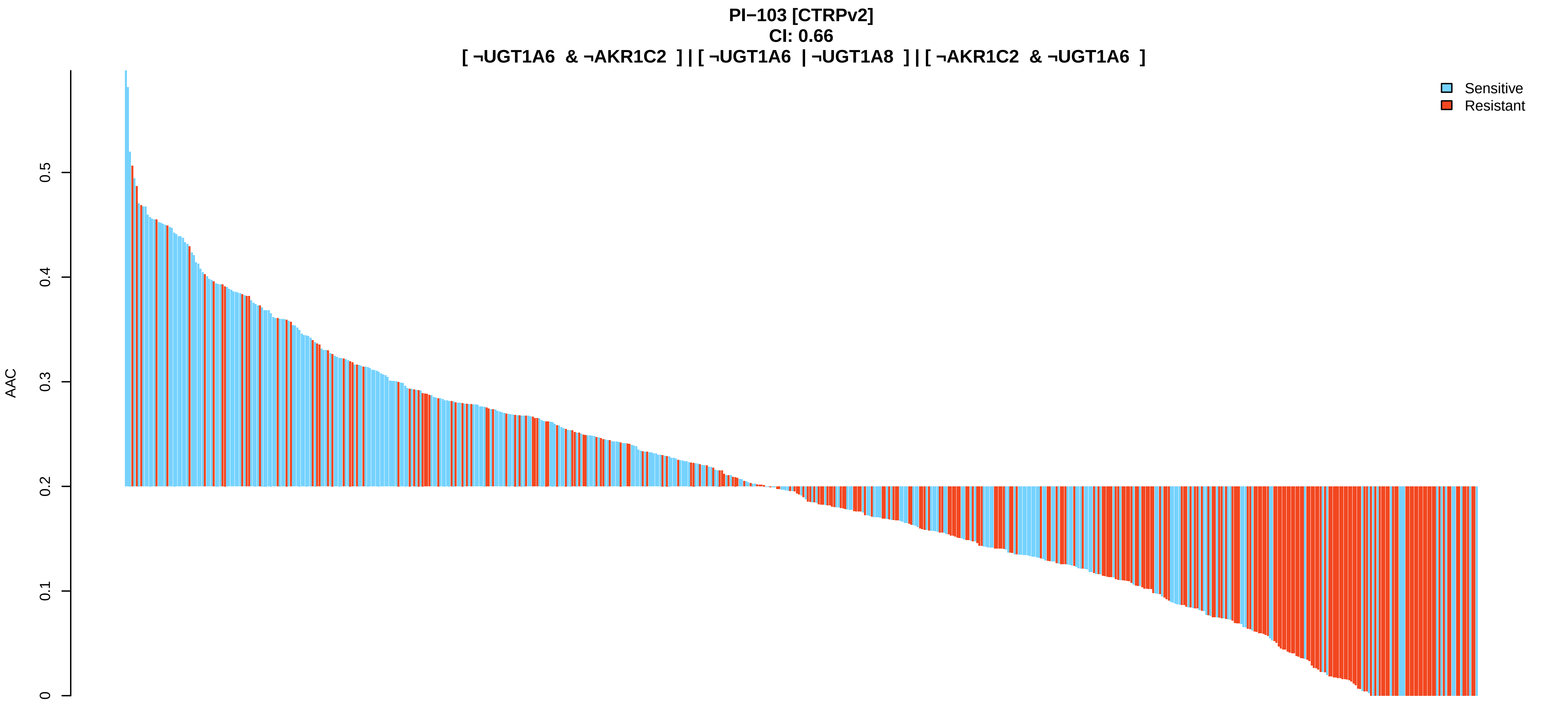
PI−103 [CTRPv2]
CI: 0.66
[ ¬UGT1A6 & ¬AKR1C2 ] | [ ¬UGT1A6 | ¬UGT1A8 ] | [ ¬AKR1C2 & ¬UGT1A6 ]
Sensitive
Resistant
AAC
0
0.1
0.2
0.3
0.4
0.5

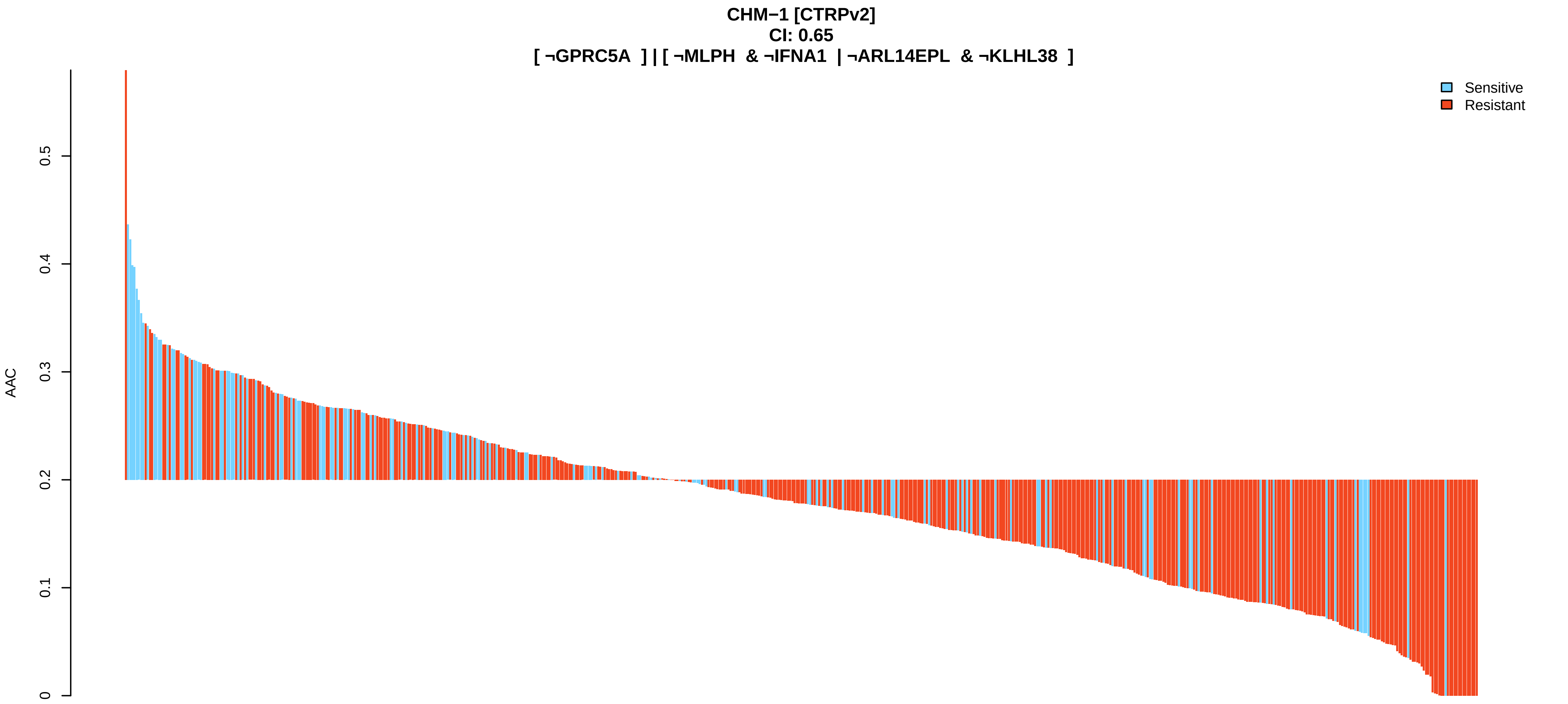
CHM−1 [CTRPv2]
CI: 0.65
[ ¬GPRC5A ] | [ ¬MLPH & ¬IFNA1 | ¬ARL14EPL & ¬KLHL38 ]
Sensitive
Resistant
AAC
0
0.1
0.2
0.3
0.4
0.5

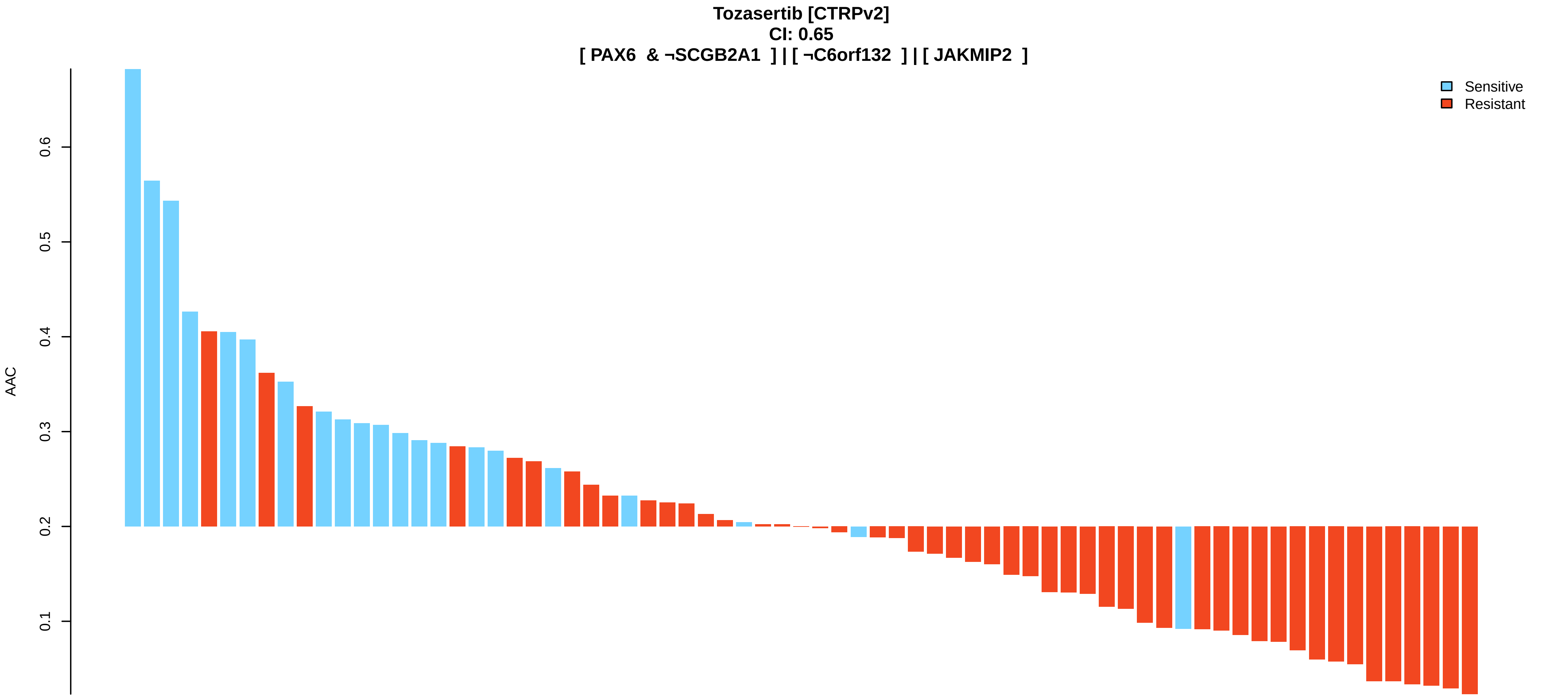
Tozasertib [CTRPv2]
CI: 0.65
[ PAX6 & ¬SCGB2A1 ] | [ ¬C6orf132 ] | [ JAKMIP2 ]
Sensitive
Resistant
AAC
0.1
0.2
0.3
0.4
0.5
0.6

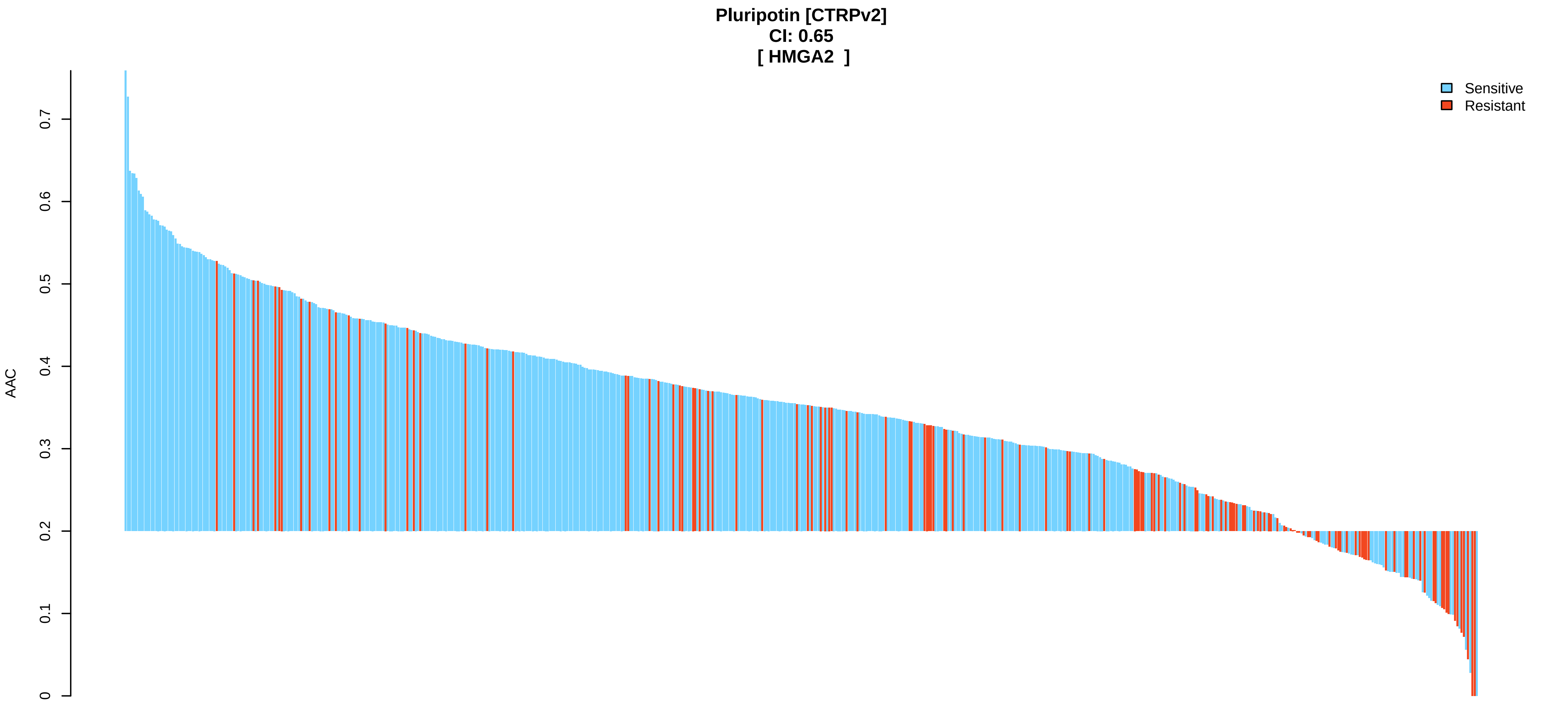
Pluripotin [CTRPv2]
CI: 0.65
[ HMGA2 ]
Sensitive
Resistant
AAC
0
0.1
0.2
0.3
0.4
0.5
0.6
0.7

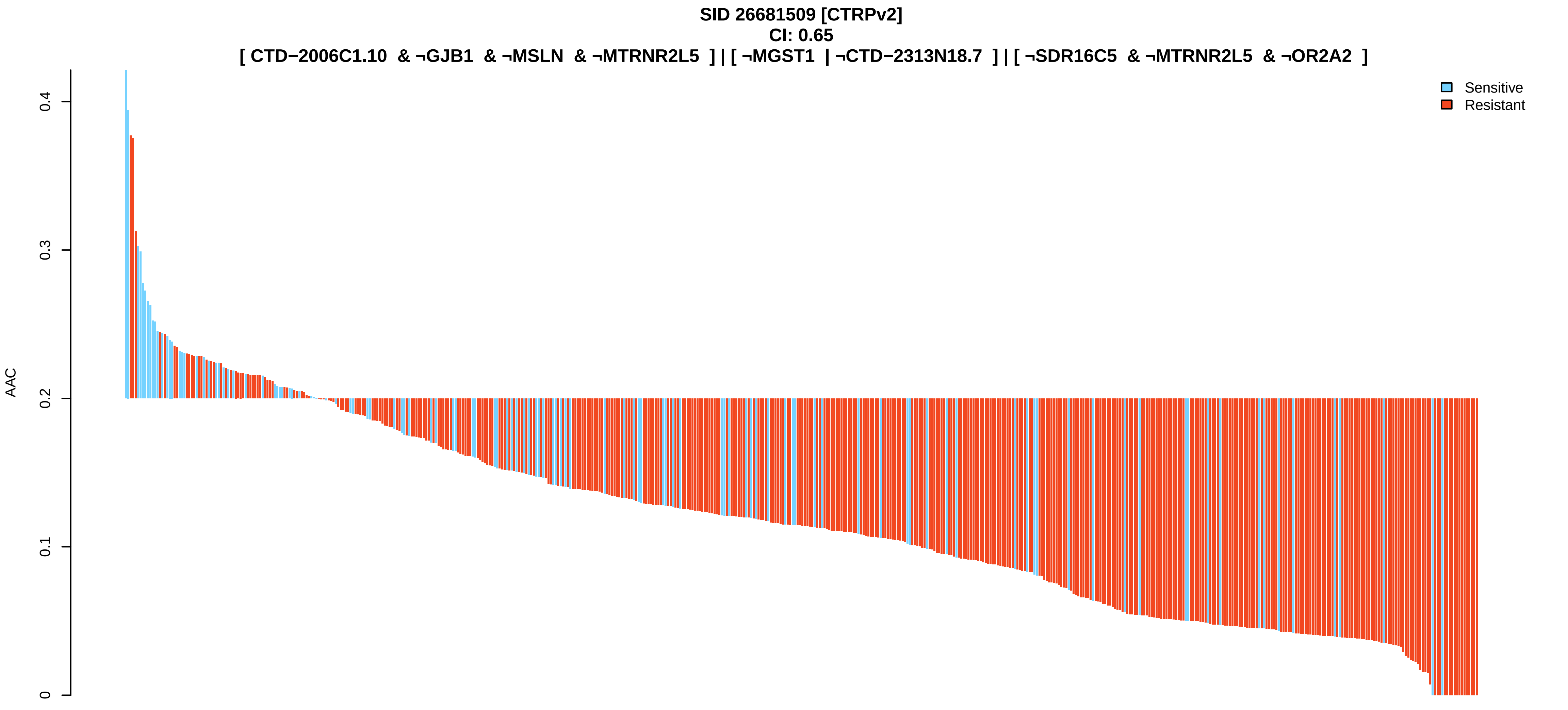
SID 26681509 [CTRPv2]
CI: 0.65
[ CTD−2006C1.10 & ¬GJB1 & ¬MSLN & ¬MTRNR2L5 ] | [ ¬MGST1 | ¬CTD−2313N18.7 ] | [ ¬SDR16C5 & ¬MTRNR2L5 & ¬OR2A2 ]
Sensitive
Resistant
AAC
0
0.1
0.2
0.3
0.4

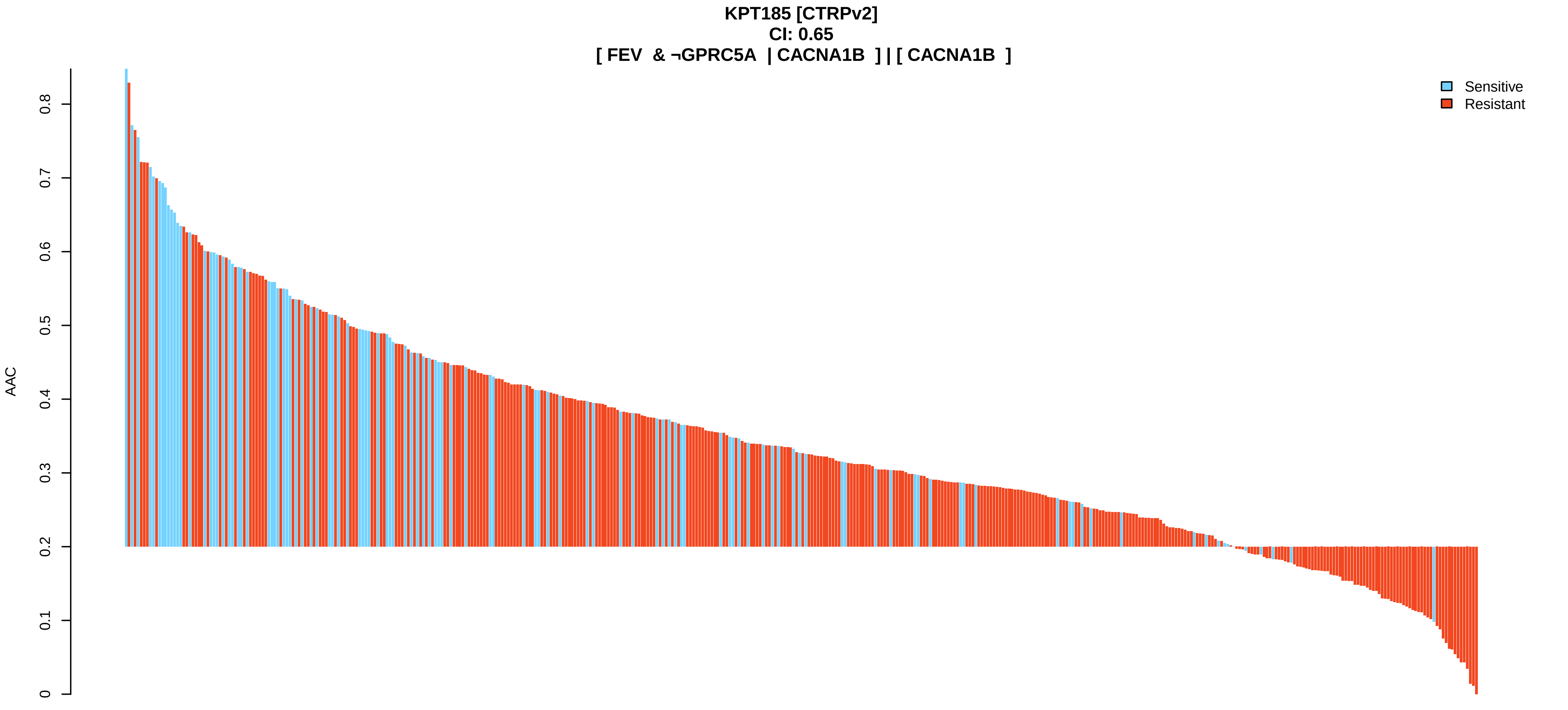
KPT185 [CTRPv2]
CI: 0.65
[ FEV & ¬GPRC5A | CACNA1B ] | [ CACNA1B ]
Sensitive
Resistant
AAC
0
0.1
0.2
0.3
0.4
0.5
0.6
0.7
0.8

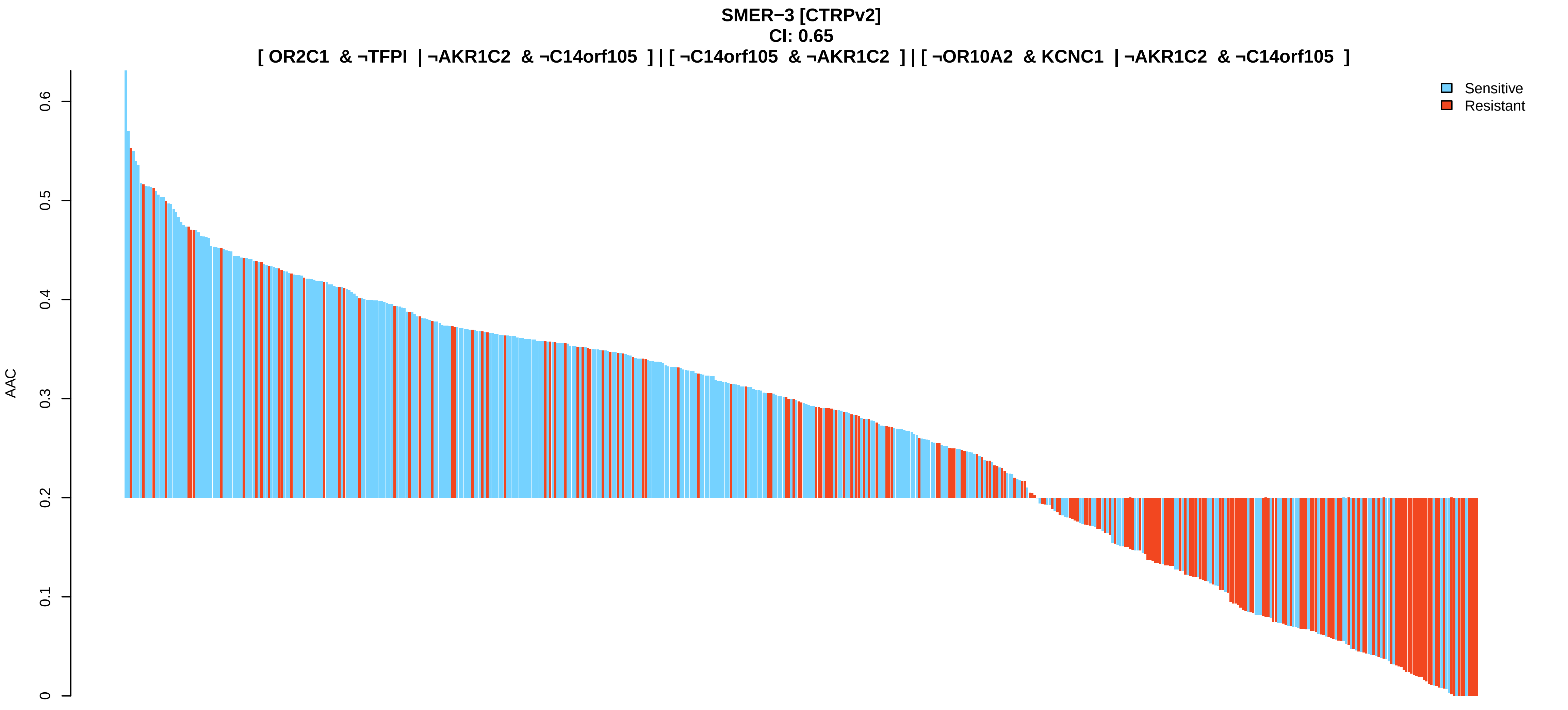

SMER−3 [CTRPv2]
CI: 0.65
[ OR2C1 & ¬TFPI | ¬AKR1C2 & ¬C14orf105 ] | [ ¬C14orf105 & ¬AKR1C2 ] | [ ¬OR10A2 & KCNC1 | ¬AKR1C2 & ¬C14orf105 ]
AAC
0
0.1
0.2
0.3
0.4
0.5
0.6
Sensitive
Resistant


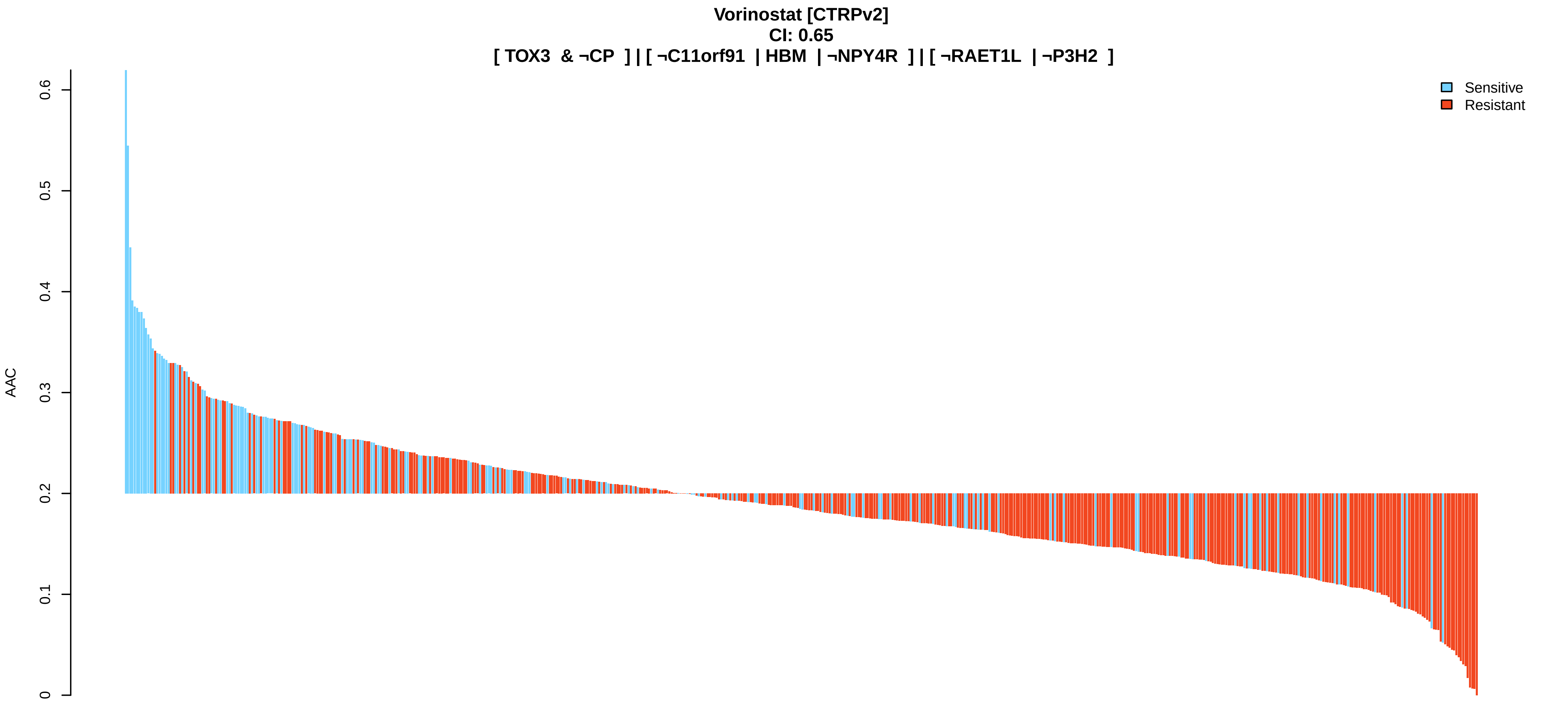

Vorinostat [CTRPv2]
CI: 0.65
[ TOX3 & ¬CP ] | [ ¬C11orf91 | HBM | ¬NPY4R ] | [ ¬RAET1L | ¬P3H2 ]
AAC
0
0.1
0.2
0.3
0.4
0.5
0.6
Sensitive
Resistant


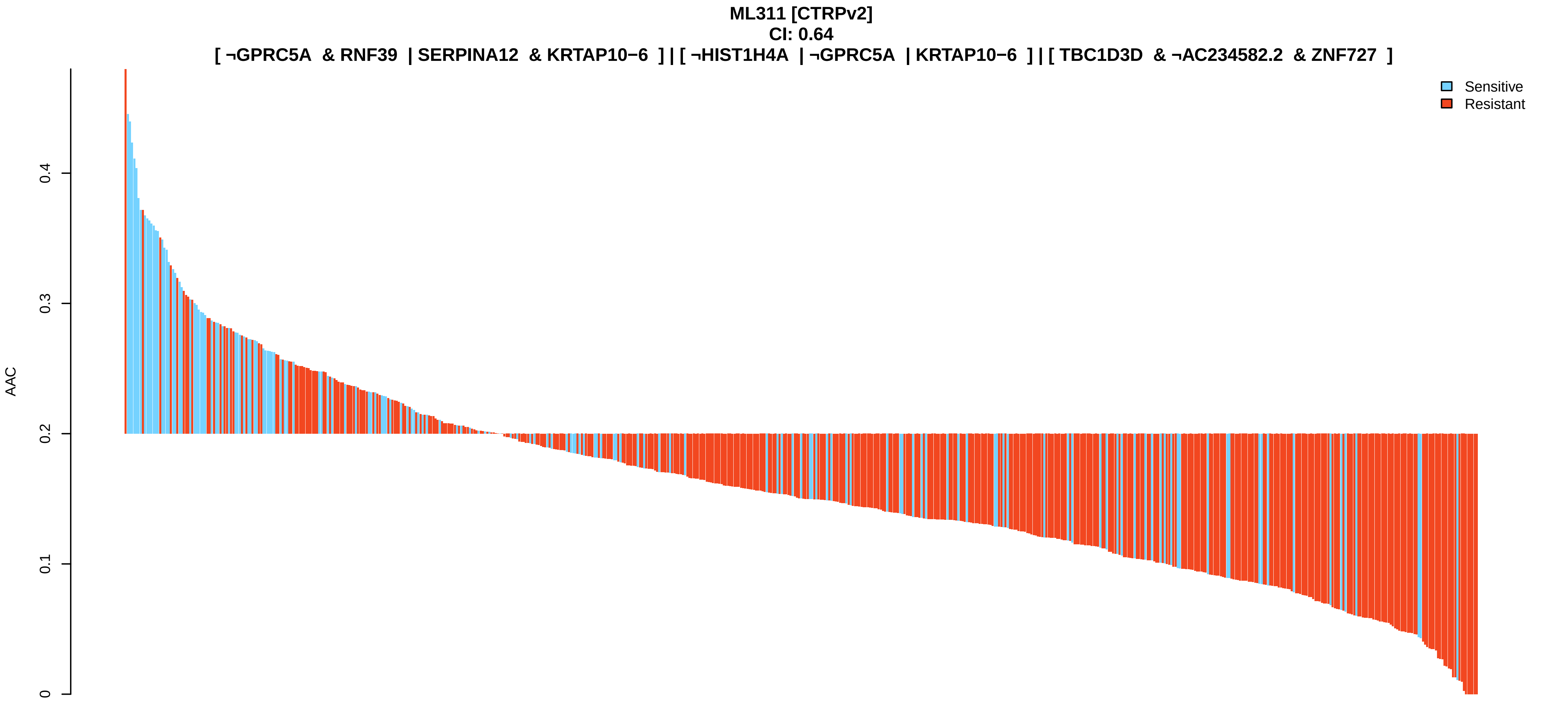

ML311 [CTRPv2]
CI: 0.64
[ ¬GPRC5A & RNF39 | SERPINA12 & KRTAP10−6 ] | [ ¬HIST1H4A | ¬GPRC5A | KRTAP10−6 ] | [ TBC1D3D & ¬AC234582.2 & ZNF727 ]
AAC
0
0.1
0.2
0.3
0.4
Sensitive
Resistant

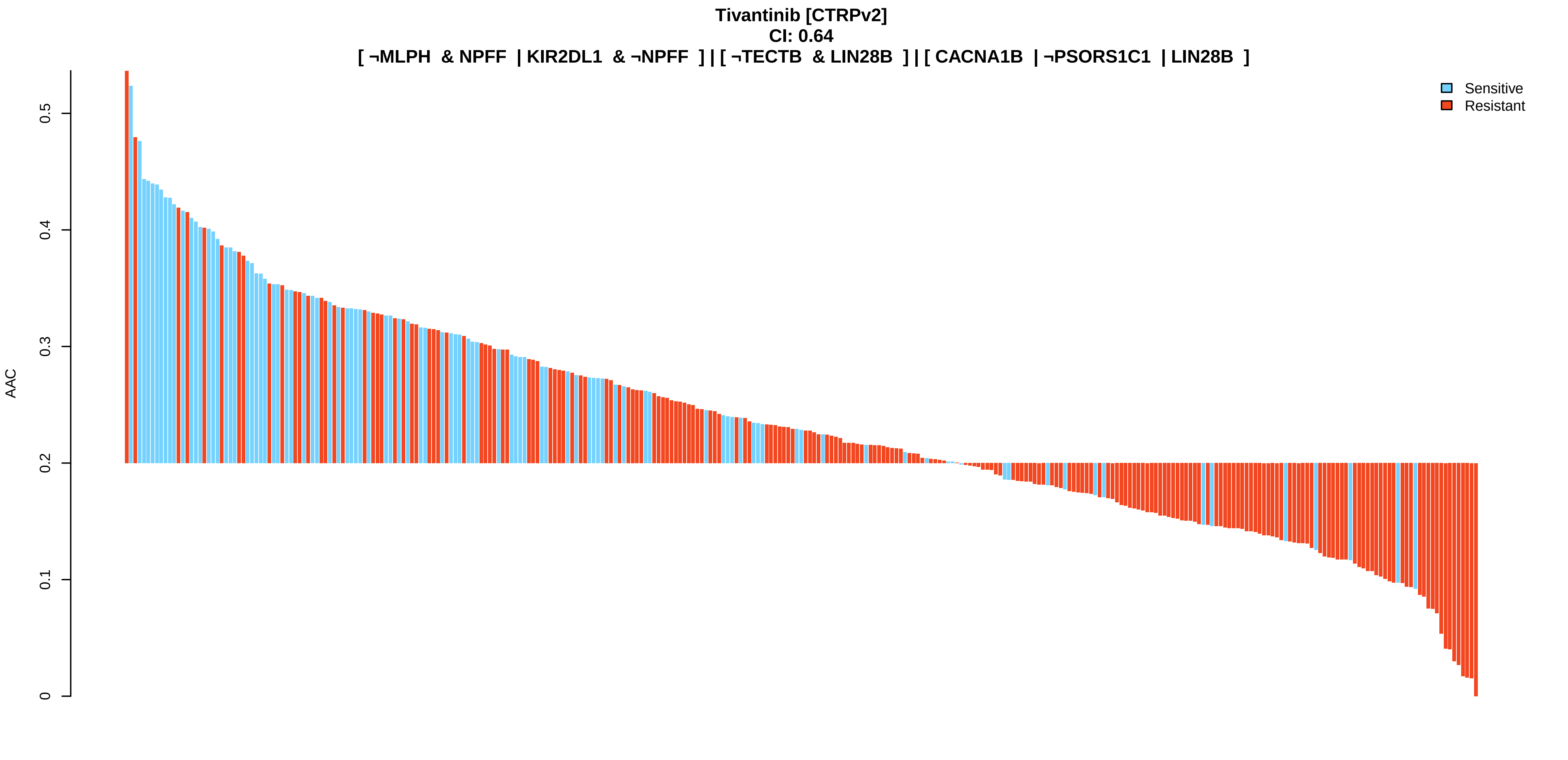
Tivantinib [CTRPv2]
CI: 0.64
[ ¬MLPH & NPFF | KIR2DL1 & ¬NPFF ] | [ ¬TECTB & LIN28B ] | [ CACNA1B | ¬PSORS1C1 | LIN28B ]
AAC
0
0.1
0.2
0.3
0.4
0.5
Sensitive
Resistant

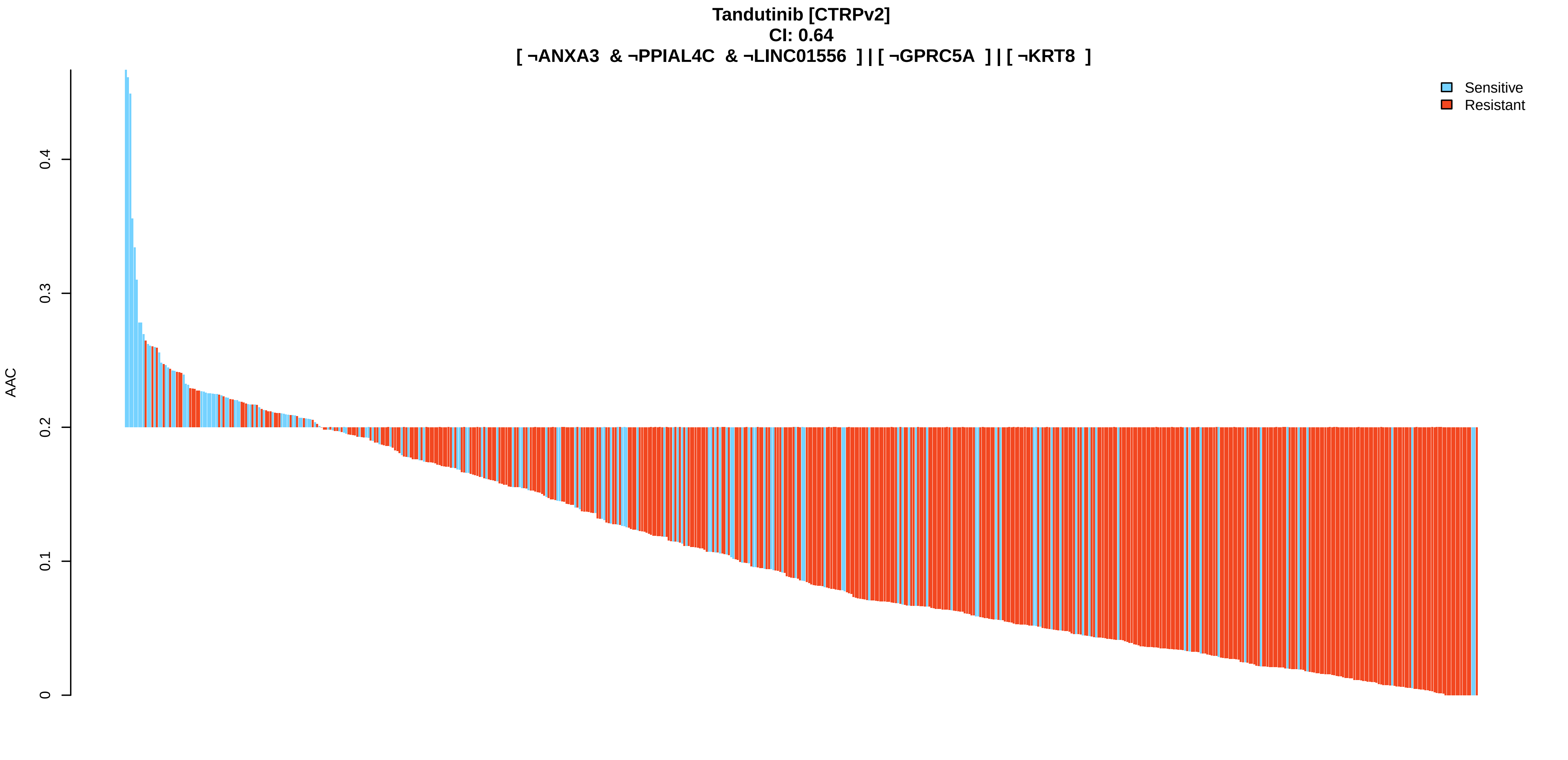
Tandutinib [CTRPv2]
CI: 0.64
[ ¬ANXA3 & ¬PPIAL4C & ¬LINC01556 ] | [ ¬GPRC5A ] | [ ¬KRT8 ]
AAC
0
0.1
0.2
0.3
0.4
Sensitive
Resistant

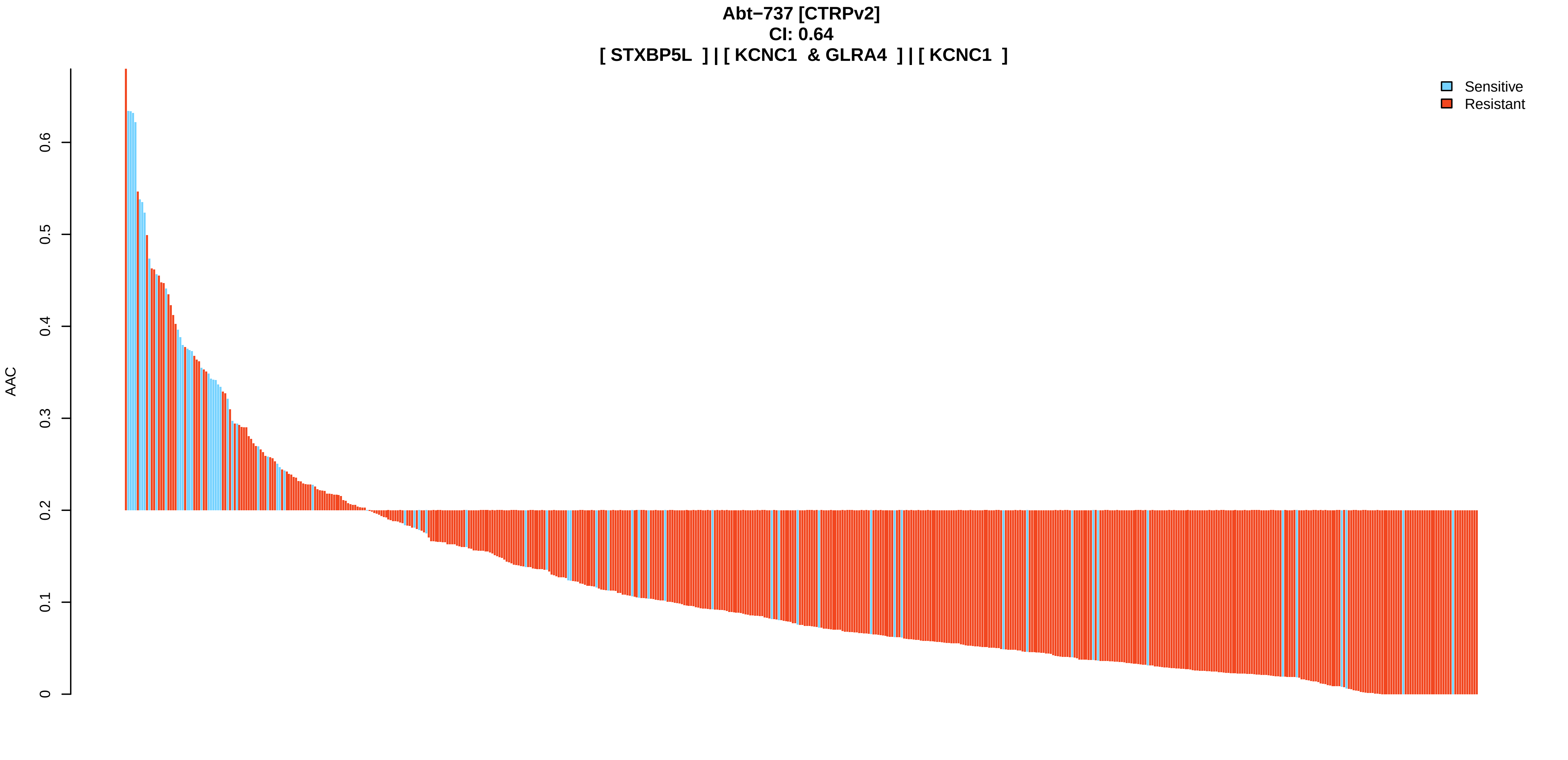
Abt−737 [CTRPv2]
CI: 0.64
[ STXBP5L ] | [ KCNC1 & GLRA4 ] | [ KCNC1 ]
AAC
0
0.1
0.2
0.3
0.4
0.5
0.6
Sensitive
Resistant

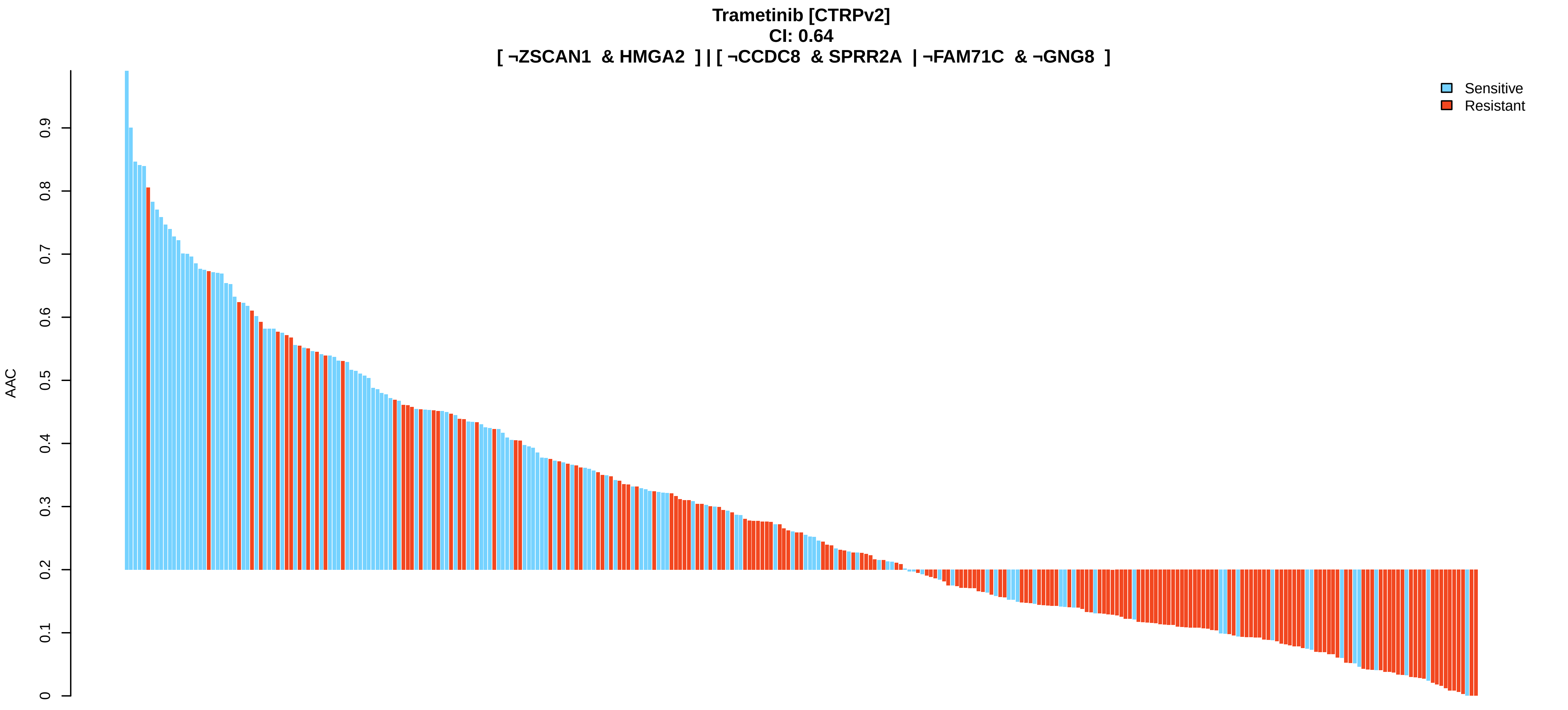
Trametinib [CTRPv2]
CI: 0.64
[ ¬ZSCAN1 & HMGA2 ] | [ ¬CCDC8 & SPRR2A | ¬FAM71C & ¬GNG8 ]
AAC
0
0.1
0.2
0.3
0.4
0.5
0.6
0.7
0.8
0.9
Sensitive
Resistant

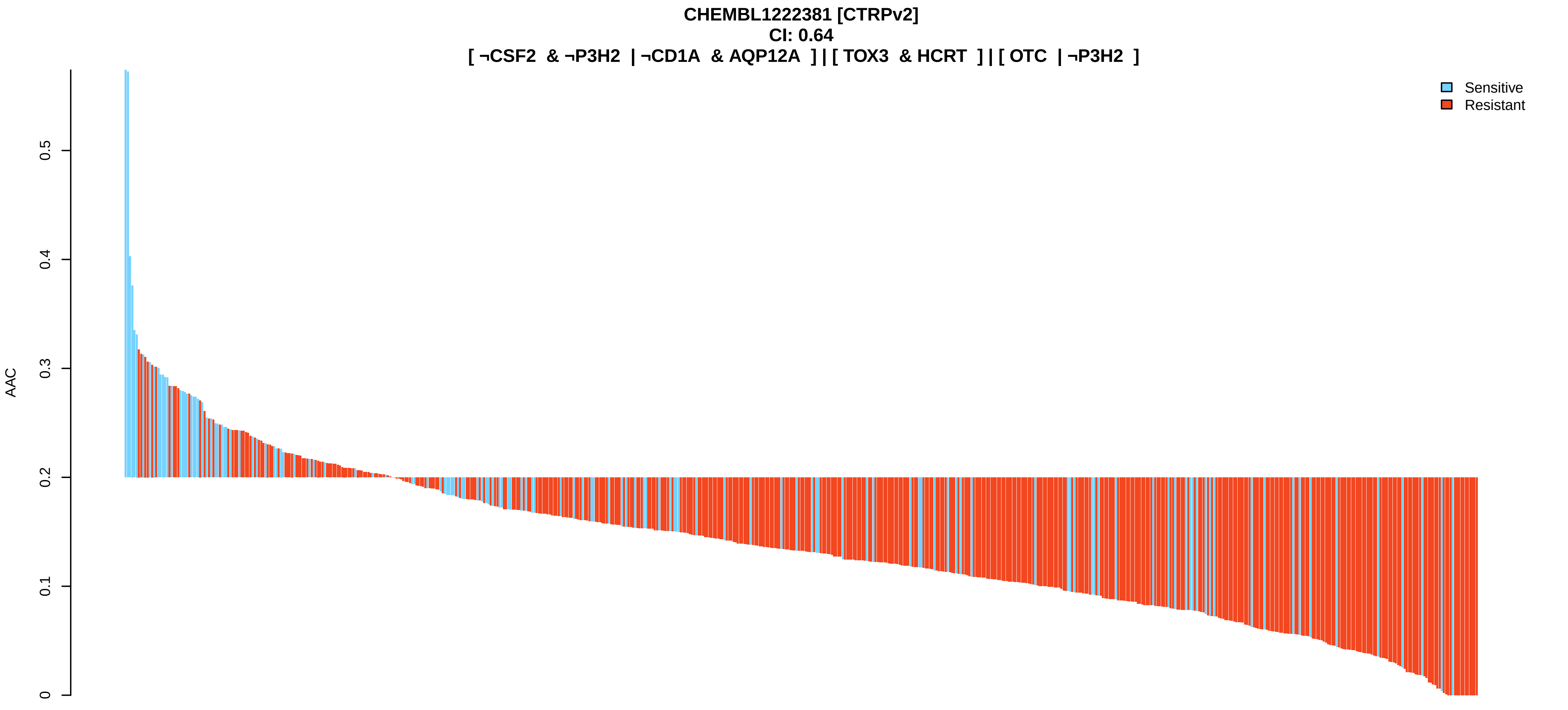
CHEMBL1222381 [CTRPv2]
CI: 0.64
[ ¬CSF2 & ¬P3H2 | ¬CD1A & AQP12A ] | [ TOX3 & HCRT ] | [ OTC | ¬P3H2 ]
AAC
0
0.1
0.2
0.3
0.4
0.5
Sensitive
Resistant

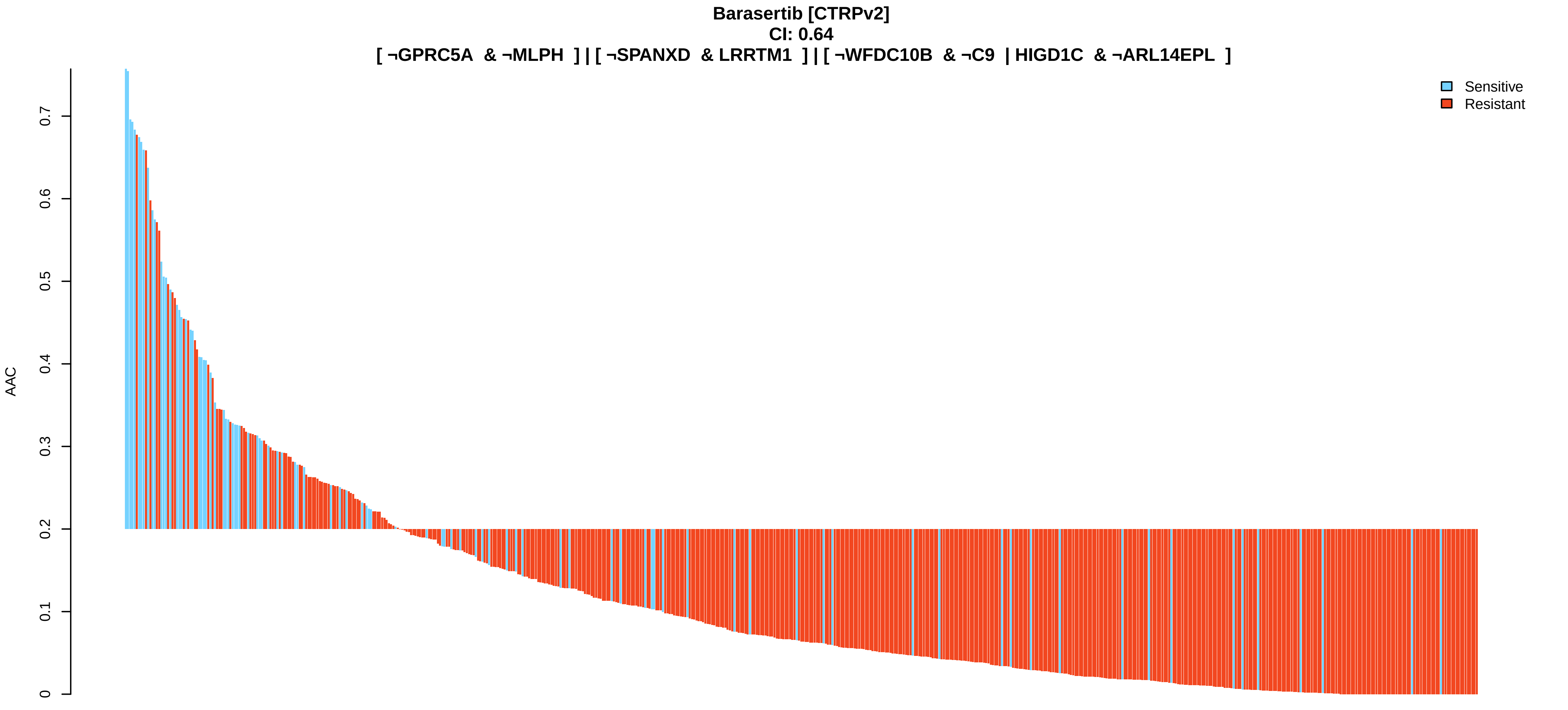
Barasertib [CTRPv2]
CI: 0.64
[ ¬GPRC5A & ¬MLPH ] | [ ¬SPANXD & LRRTM1 ] | [ ¬WFDC10B & ¬C9 | HIGD1C & ¬ARL14EPL ]
AAC
0
0.1
0.2
0.3
0.4
0.5
0.6
0.7
Sensitive
Resistant

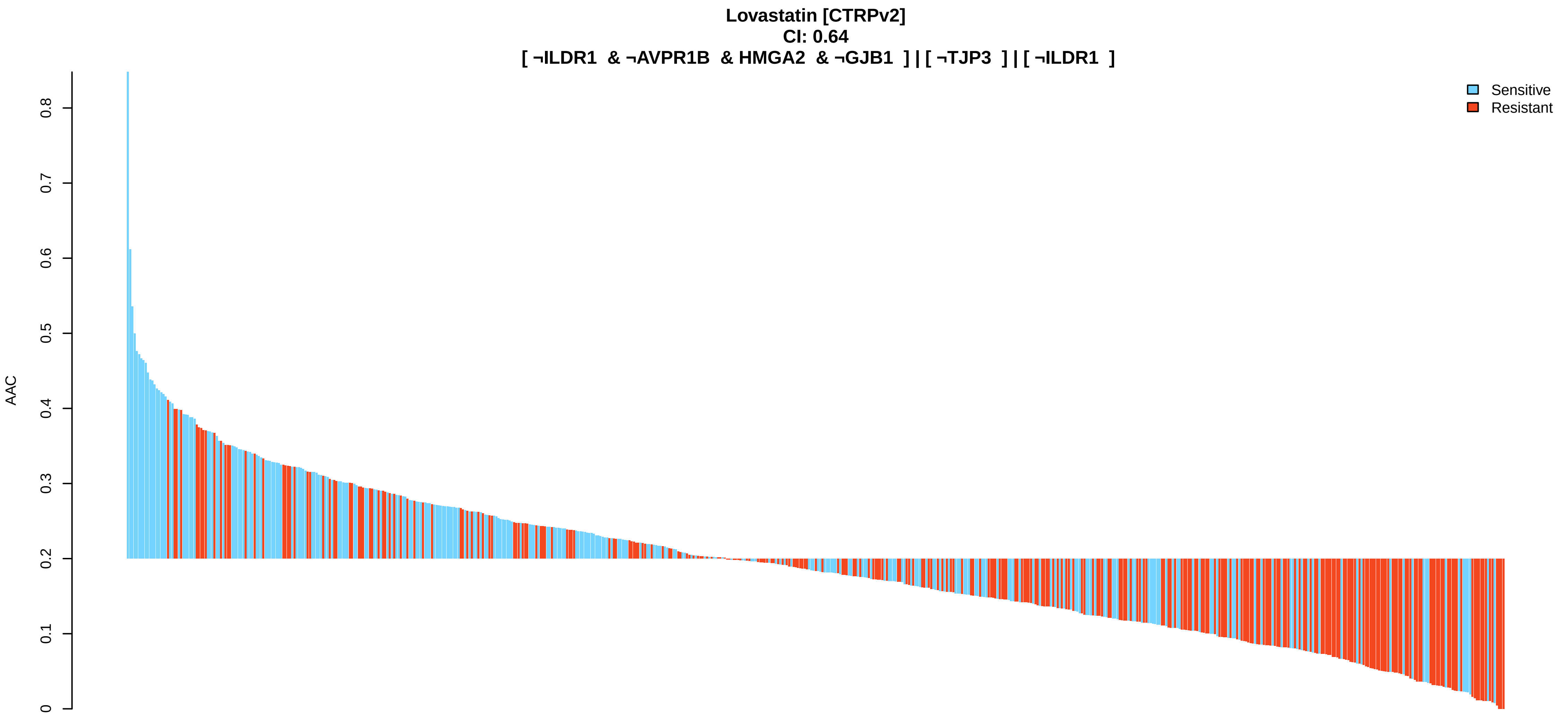
Lovastatin [CTRPv2]
CI: 0.64
[ ¬ILDR1 & ¬AVPR1B & HMGA2 & ¬GJB1 ] | [ ¬TJP3 ] | [ ¬ILDR1 ]
AAC
0
0.1
0.2
0.3
0.4
0.5
0.6
0.7
0.8
Sensitive
Resistant

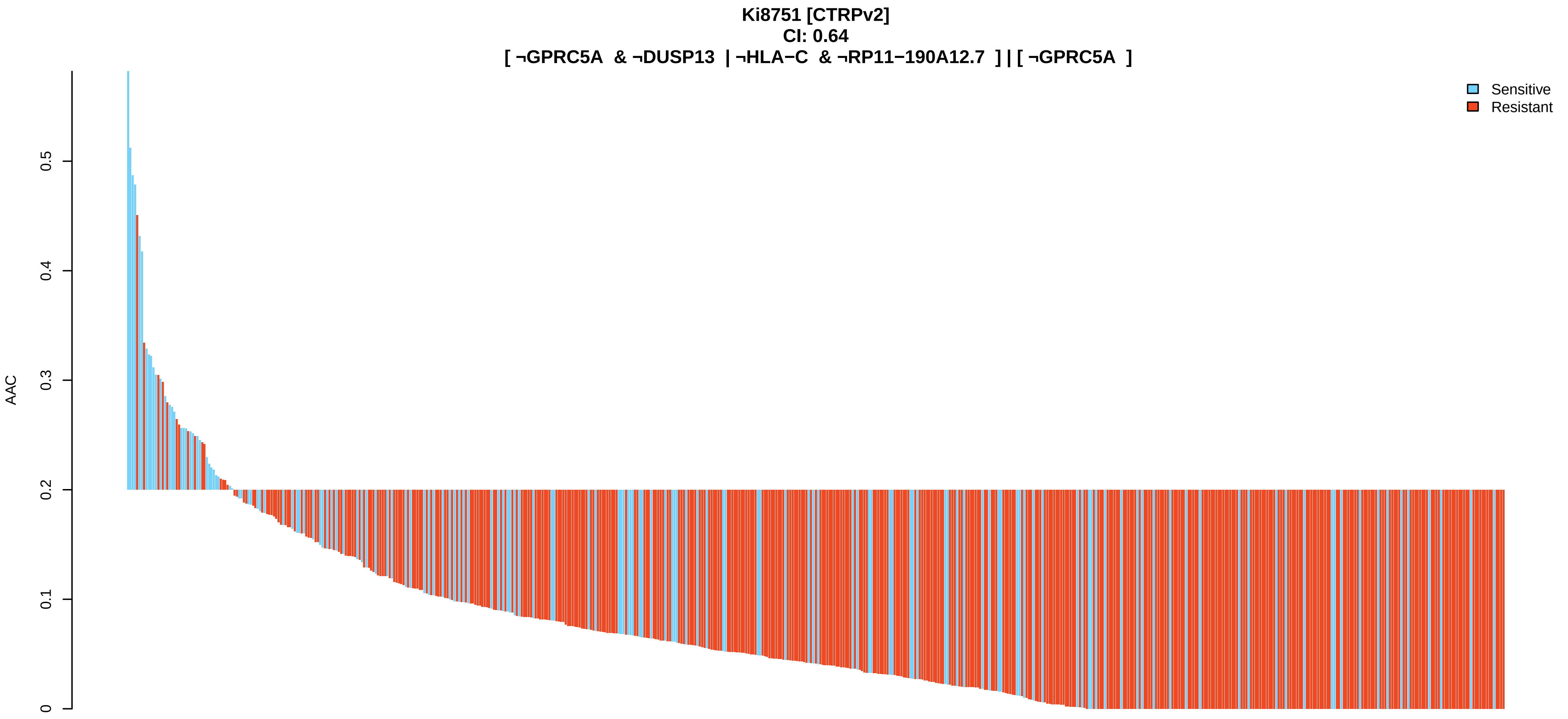
Ki8751 [CTRPv2]
CI: 0.64
[ ¬GPRC5A & ¬DUSP13 | ¬HLA−C & ¬RP11−190A12.7 ] | [ ¬GPRC5A ]
AAC
0
0.1
0.2
0.3
0.4
0.5
Sensitive
Resistant

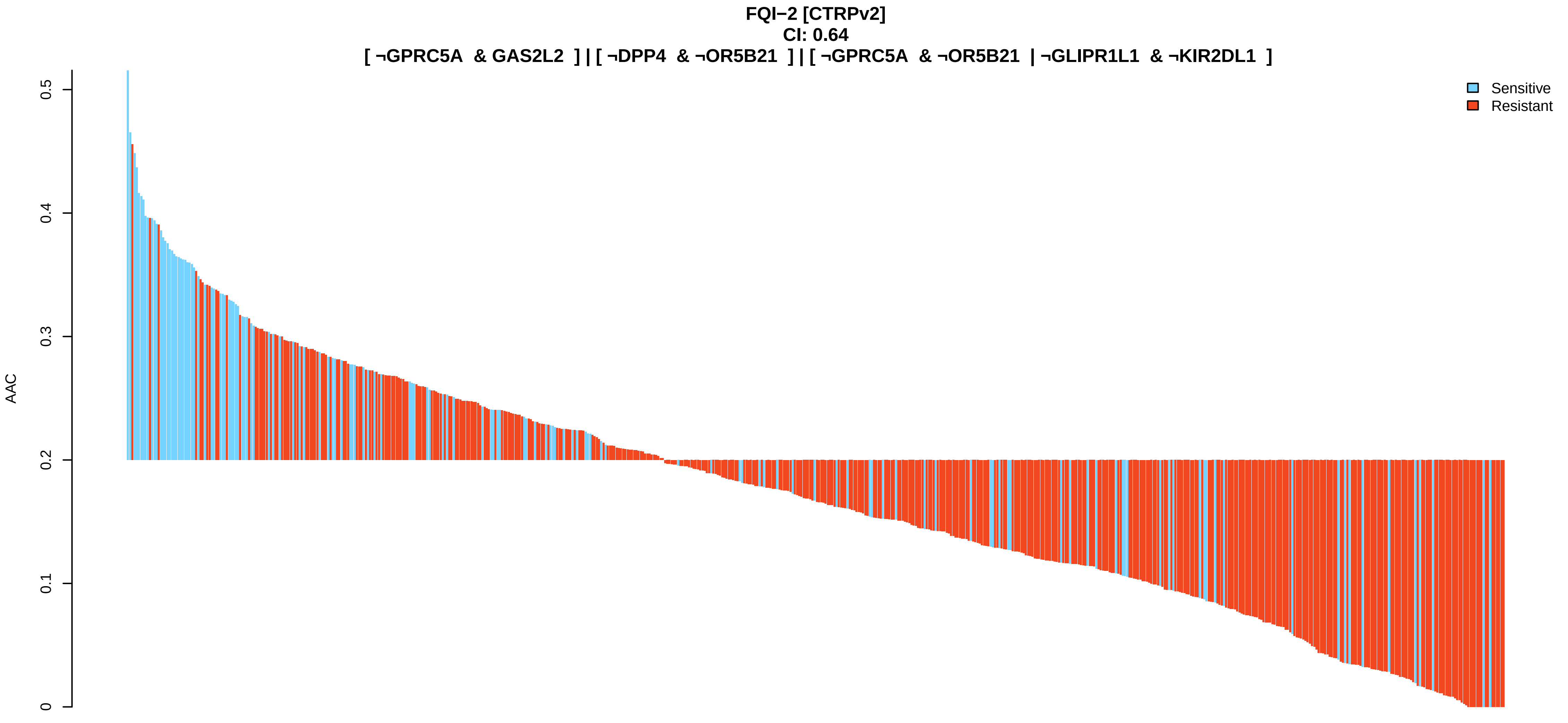
FQI−2 [CTRPv2]
CI: 0.64
[ ¬GPRC5A & GAS2L2 ] | [ ¬DPP4 & ¬OR5B21 ] | [ ¬GPRC5A & ¬OR5B21 | ¬GLIPR1L1 & ¬KIR2DL1 ]
AAC
0
0.1
0.2
0.3
0.4
0.5
Sensitive
Resistant

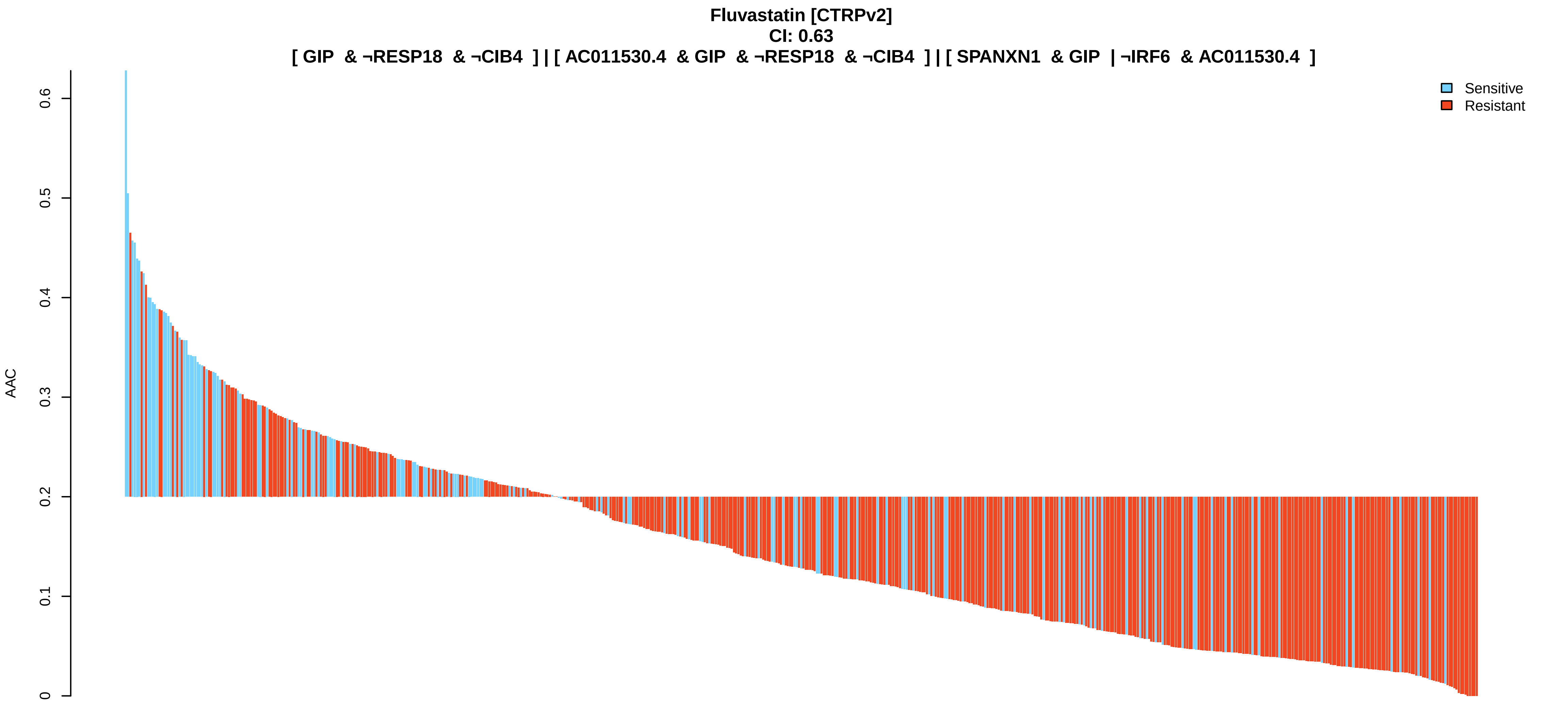
Fluvastatin [CTRPv2]
CI: 0.63
[ GIP & ¬RESP18 & ¬CIB4 ] | [ AC011530.4 & GIP & ¬RESP18 & ¬CIB4 ] | [ SPANXN1 & GIP | ¬IRF6 & AC011530.4 ]
Sensitive
Resistant
AAC
0
0.1
0.2
0.3
0.4
0.5
0.6

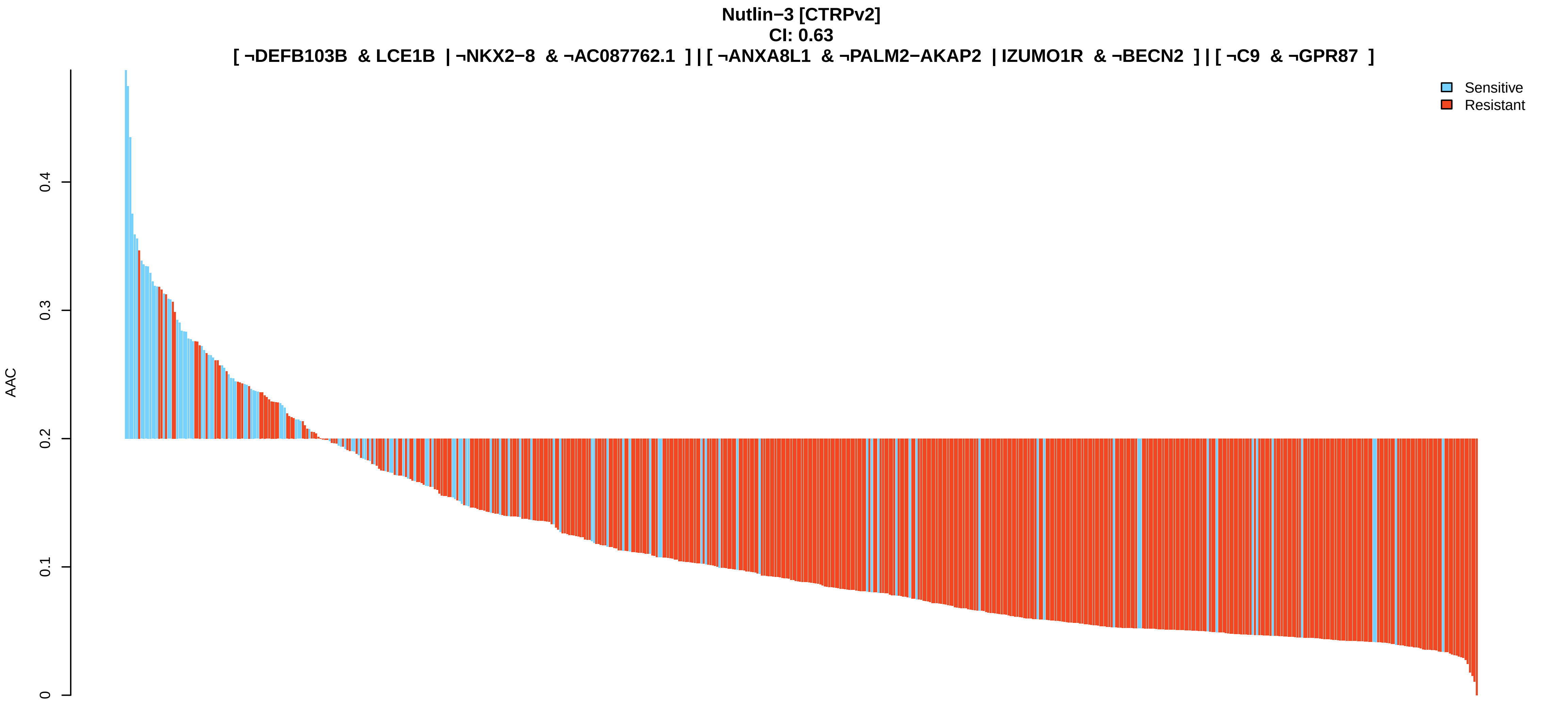
Nutlin−3 [CTRPv2]
CI: 0.63
[ ¬DEFB103B & LCE1B | ¬NKX2−8 & ¬AC087762.1 ] | [ ¬ANXA8L1 & ¬PALM2−AKAP2 | IZUMO1R & ¬BECN2 ] | [ ¬C9 & ¬GPR87 ]
Sensitive
Resistant
AAC
0
0.1
0.2
0.3
0.4

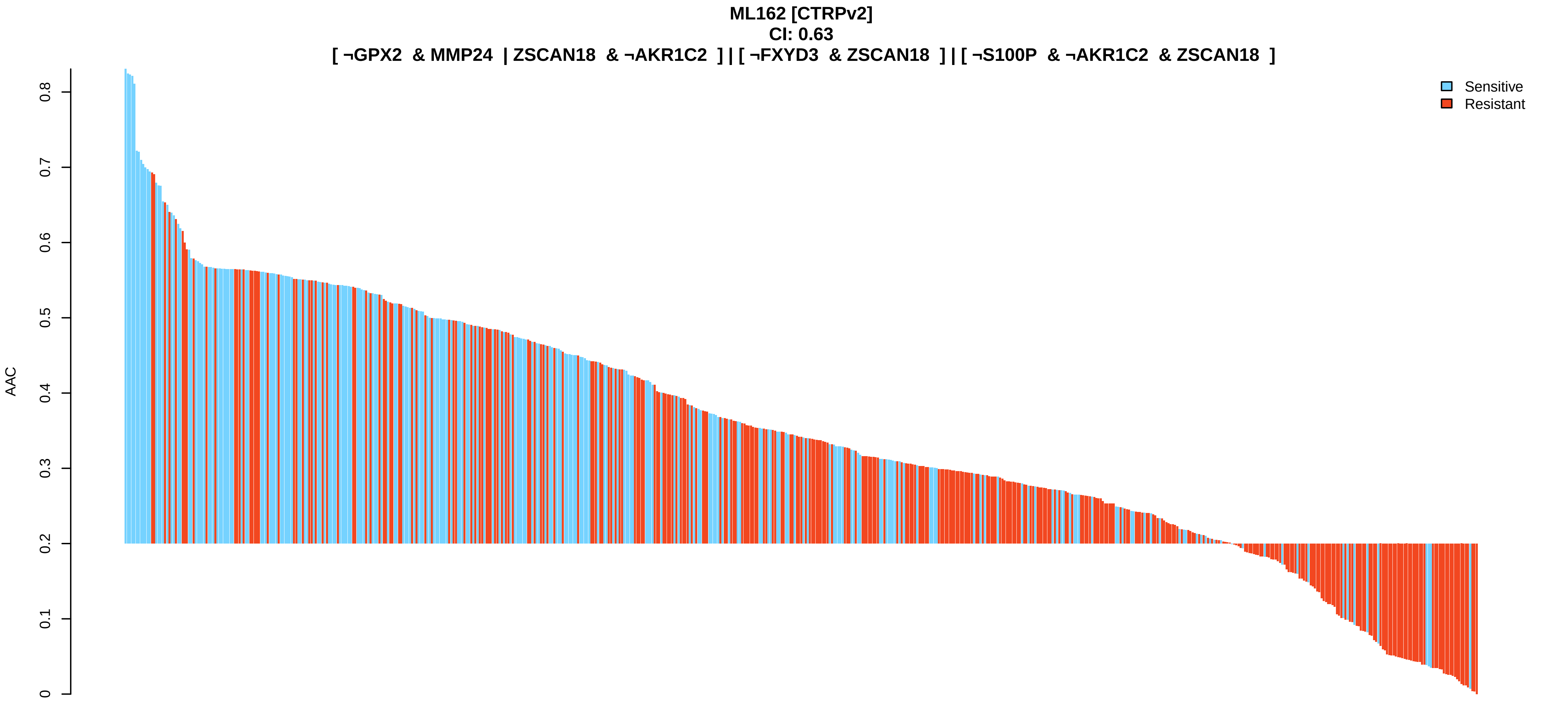
ML162 [CTRPv2]
CI: 0.63
[ ¬GPX2 & MMP24 | ZSCAN18 & ¬AKR1C2 ] | [ ¬FXYD3 & ZSCAN18 ] | [ ¬S100P & ¬AKR1C2 & ZSCAN18 ]
Sensitive
Resistant
AAC
0
0.1
0.2
0.3
0.4
0.5
0.6
0.7
0.8

**BRD1812 [CTRPv2]**
**CI: 0.63**
**[ ¬ABCC3 & LIN28B | ¬CTRB1 & ¬ARL14EPL ] | [ ¬ABCC3 ] | [ ¬ABCC3 & ¬ARL14EPL | ¬ABCC3 & ¬PSG11 ]**

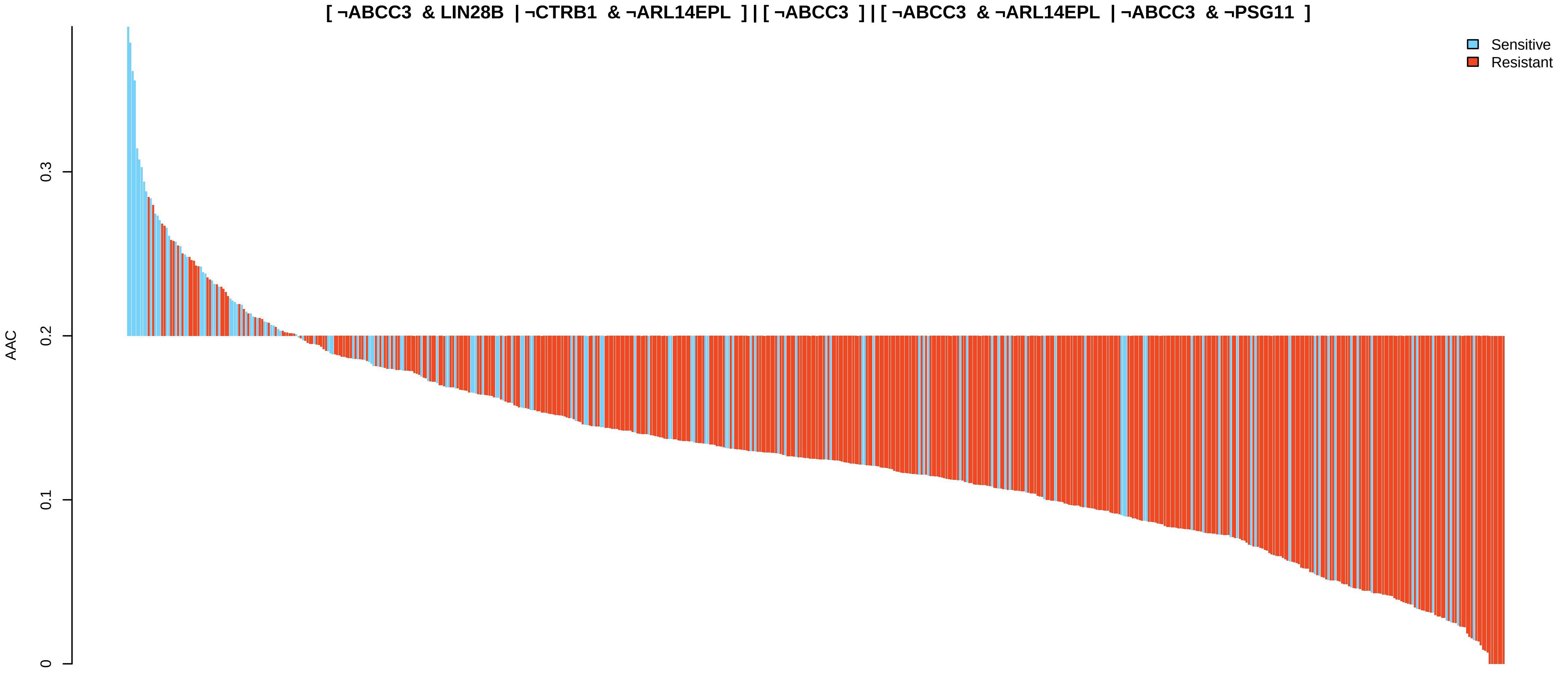


**YK 4−279 [CTRPv2]**
**CI: 0.63**
**[ ¬GPRC5A & ¬C9 | HES3 & ¬MLPH ] | [ ¬GPRC5A & ¬C9 | ¬TFPI & ¬SLC34A1 ] | [ HES3 & ¬MLPH | CACNA1B & ¬PLAC8 ]**

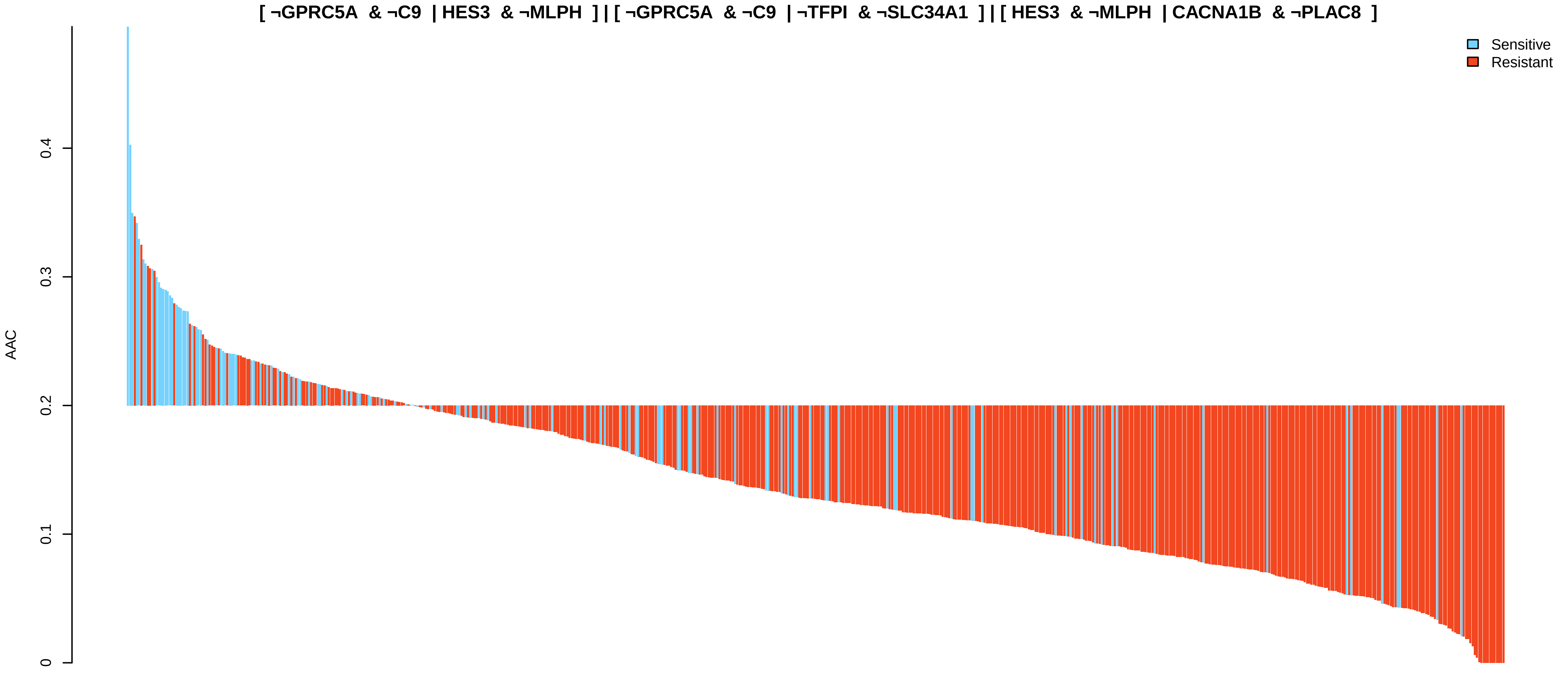


**Sepantronium bromide [CTRPv2]**
**CI: 0.63**
**[ OR2B2 & ¬AC011513.3 | LIN28B & KLF14 ] | [ OR2B2 & ¬AC011513.3 | CCDC63 & ¬TMIGD1 ] | [ ¬AC011513.3 & CTD−2561J22.3 | ¬AKR1C2 & LIN28B ]**

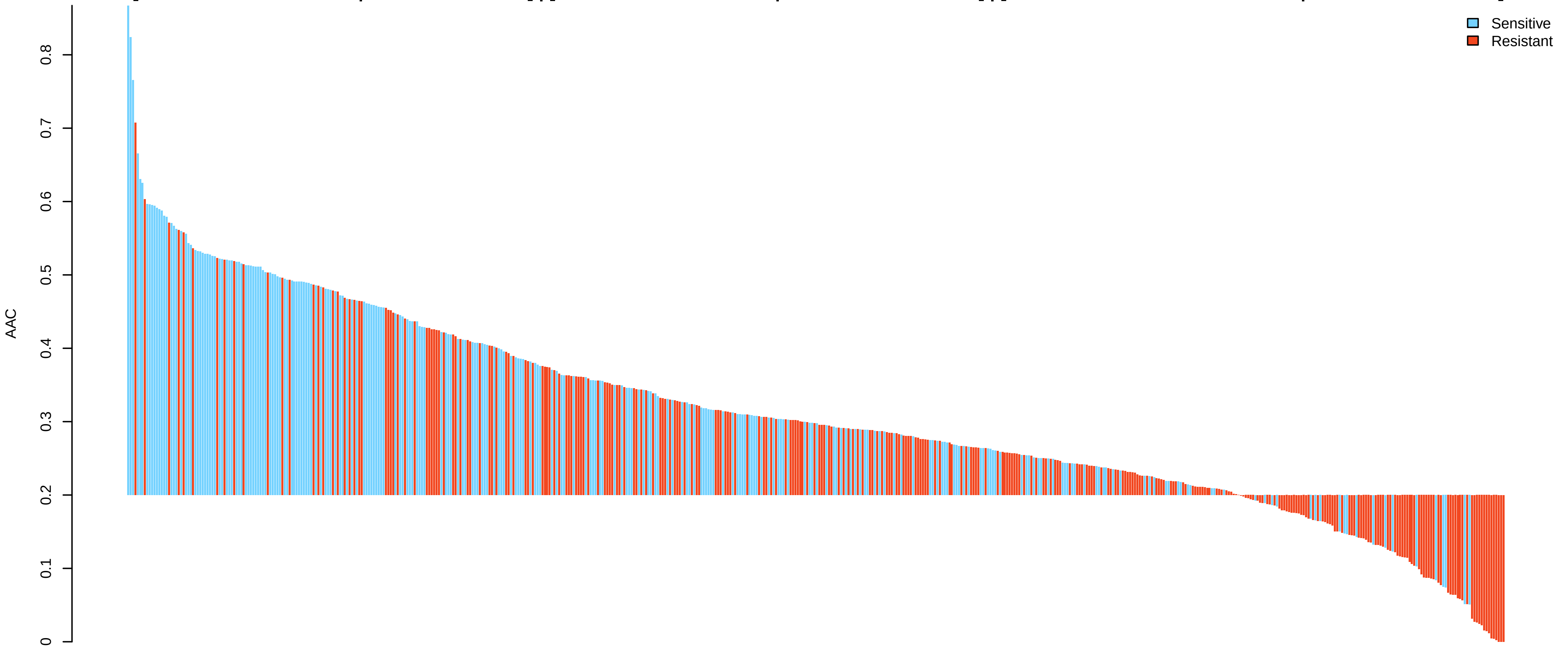

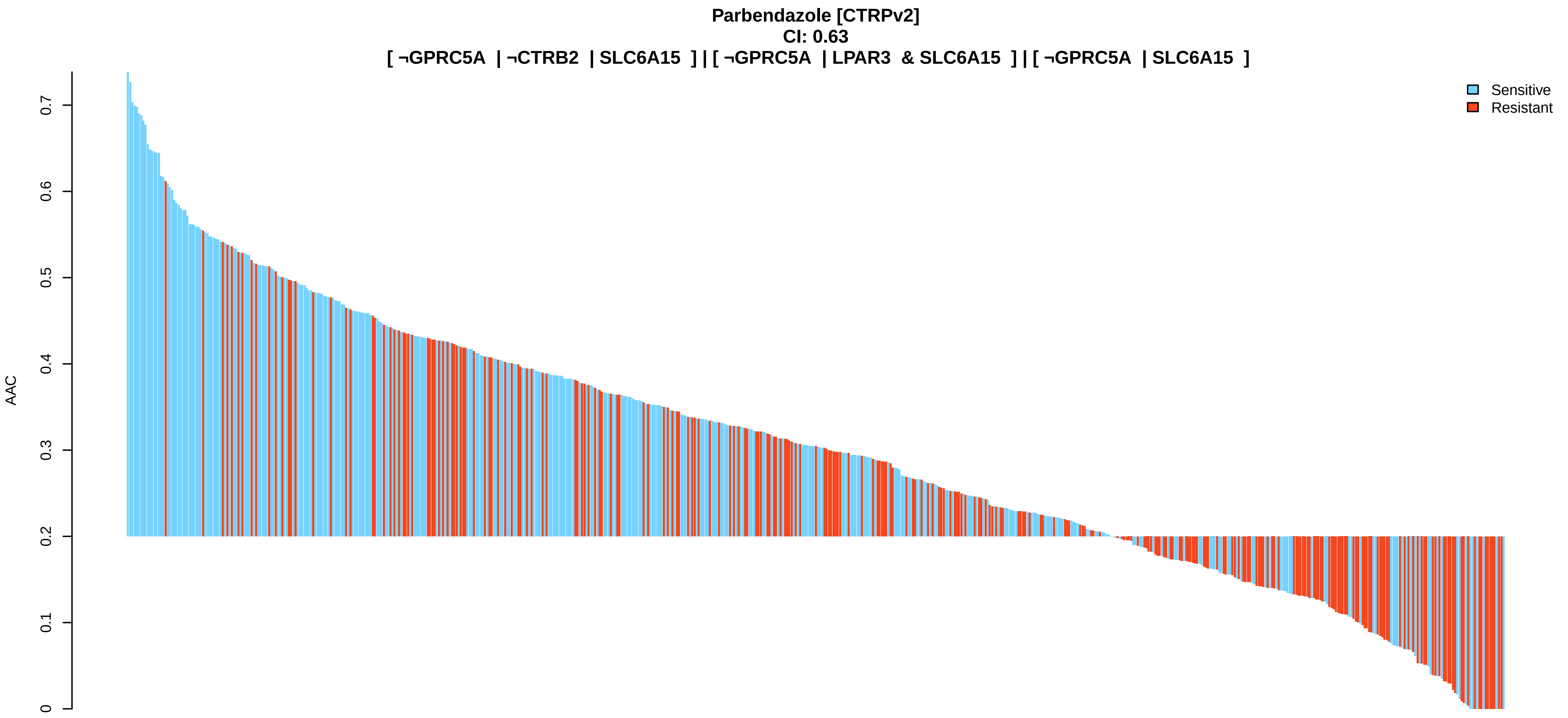
Parbendazole [CTRPv2]
CI: 0.63
[ ¬GPRC5A | ¬CTRB2 | SLC6A15 ] | [ ¬GPRC5A | LPAR3 & SLC6A15 ] | [ ¬GPRC5A | SLC6A15 ]
Sensitive
Resistant
AAC
0
0.1
0.2
0.3
0.4
0.5
0.6
0.7

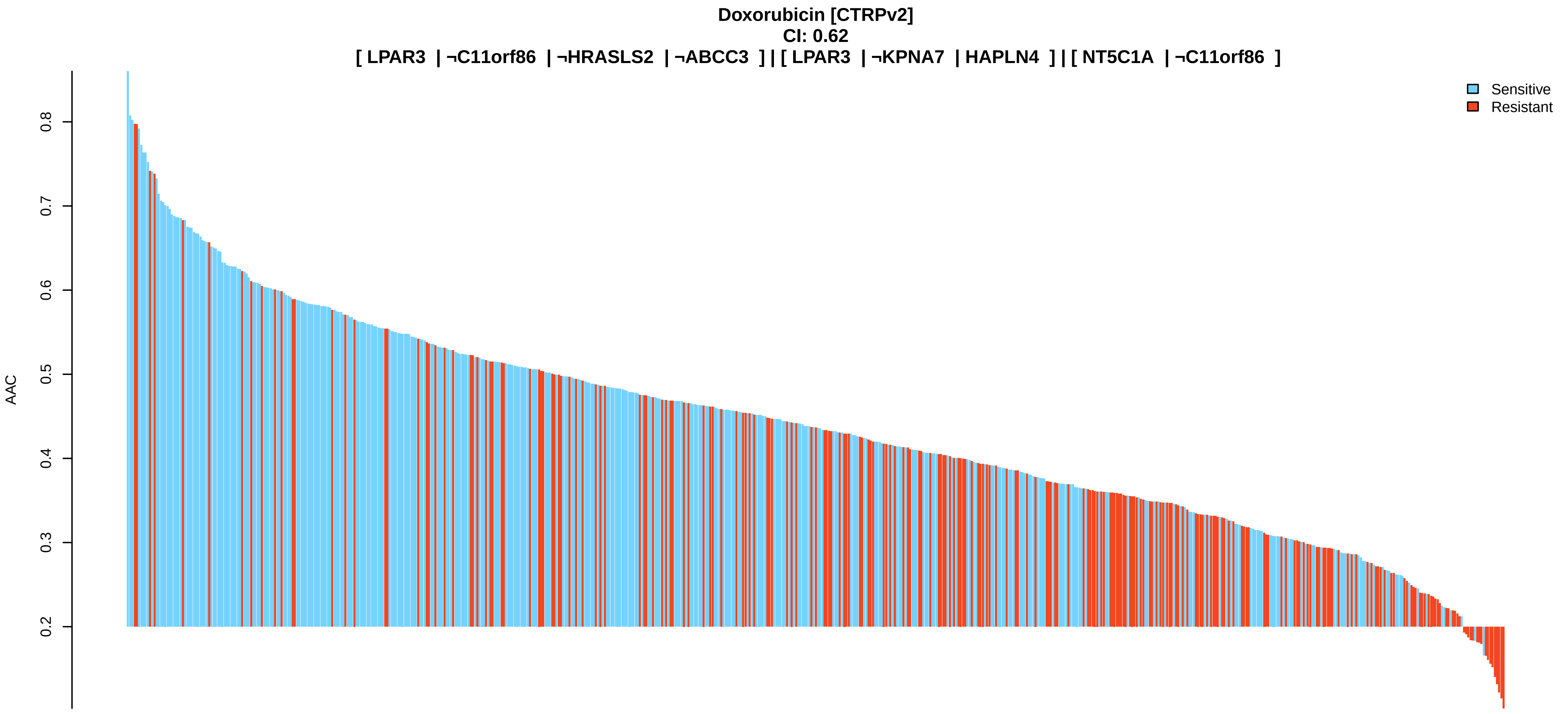
Doxorubicin [CTRPv2]
CI: 0.62
[ LPAR3 | ¬C11orf86 | ¬HRASLS2 | ¬ABCC3 ] | [ LPAR3 | ¬KPNA7 | HAPLN4 ] | [ NT5C1A | ¬C11orf86 ]
Sensitive
Resistant
AAC
0.2
0.3
0.4
0.5
0.6
0.7
0.8

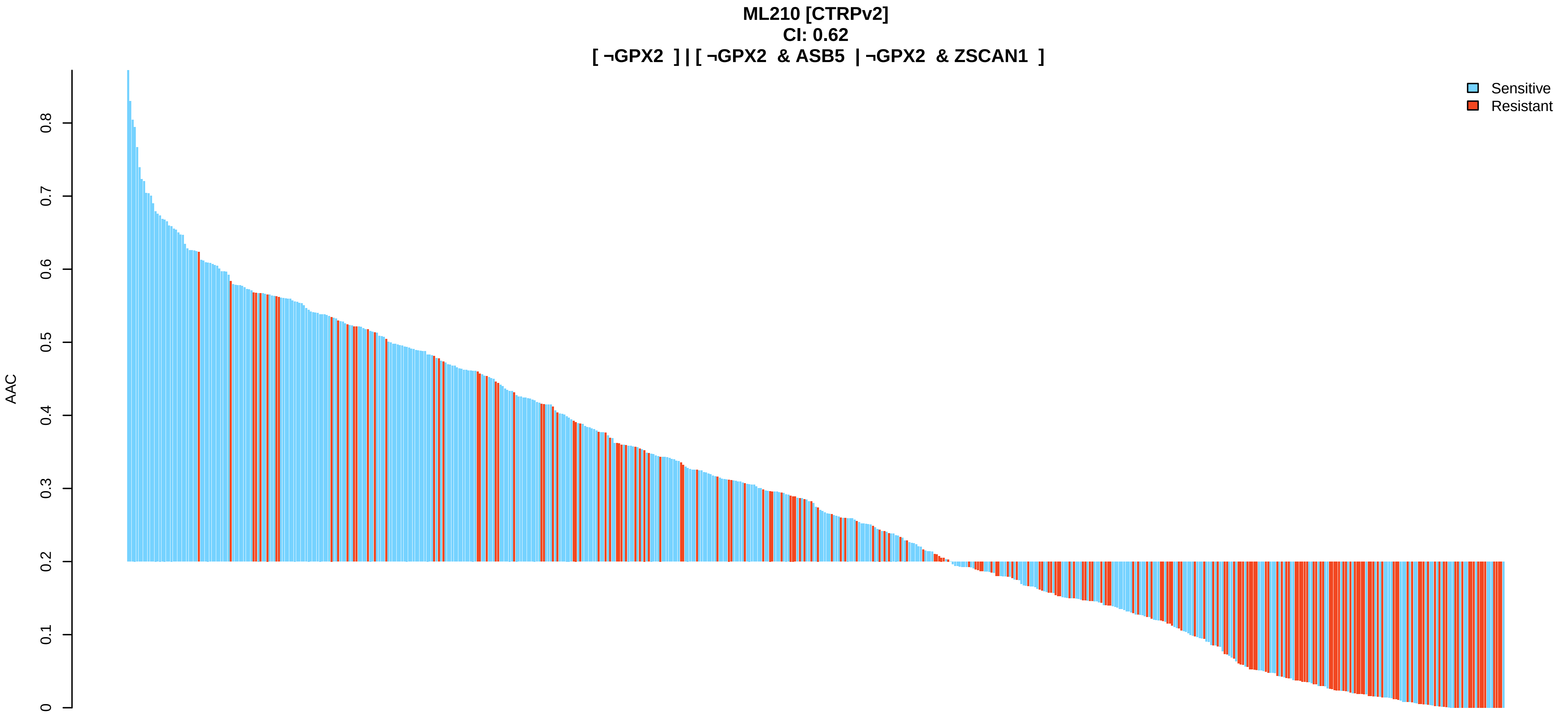
ML210 [CTRPv2]
CI: 0.62
[ ¬GPX2 ] | [ ¬GPX2 & ASB5 | ¬GPX2 & ZSCAN1 ]
Sensitive
Resistant
AAC
0
0.1
0.2
0.3
0.4
0.5
0.6
0.7
0.8

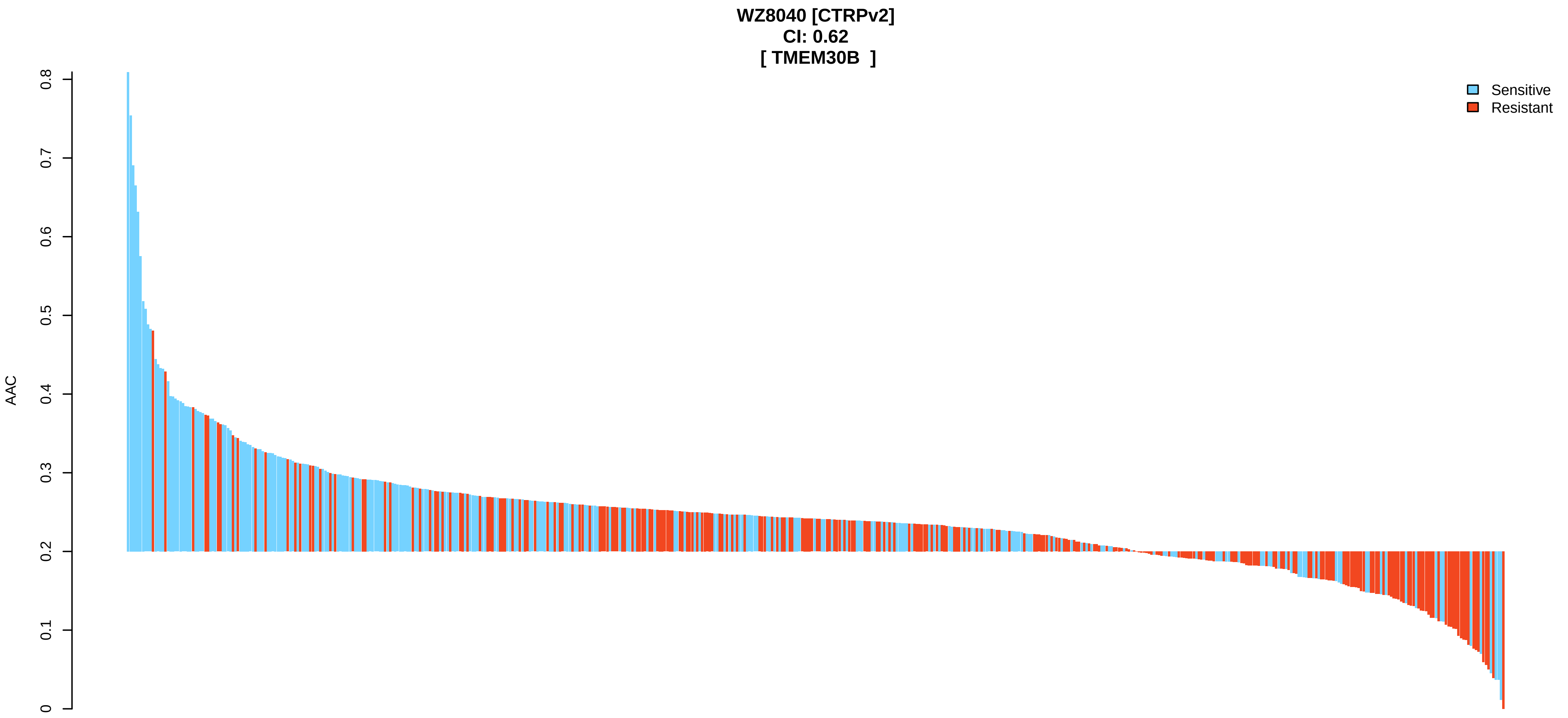
WZ8040 [CTRPv2]
CI: 0.62
[ TMEM30B ]
Sensitive
Resistant
AAC
0
0.1
0.2
0.3
0.4
0.5
0.6
0.7
0.8

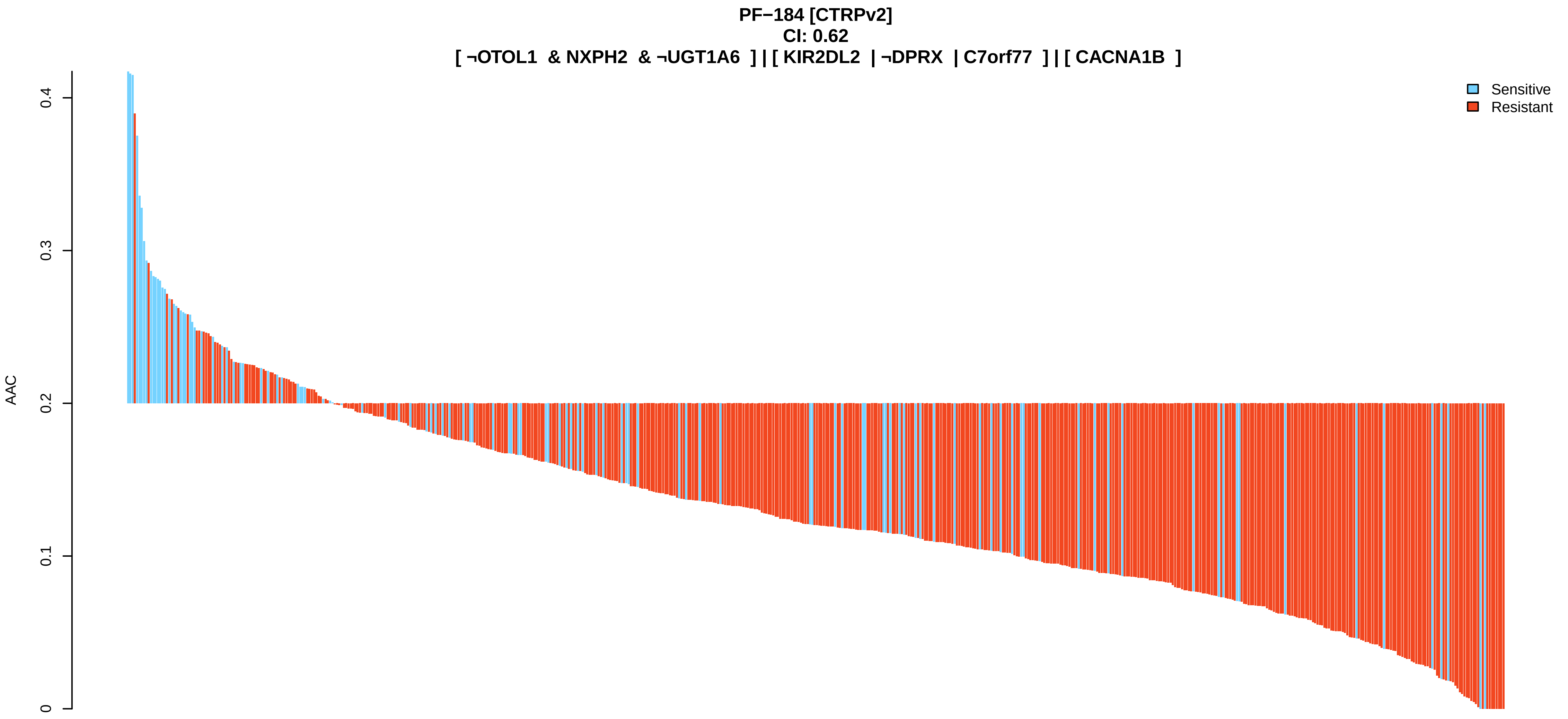
PF−184 [CTRPv2]
CI: 0.62
[ ¬OTOL1 & NXPH2 & ¬UGT1A6 ] | [ KIR2DL2 | ¬DPRX | C7orf77 ] | [ CACNA1B ]
Sensitive
Resistant
AAC
0
0.1
0.2
0.3
0.4

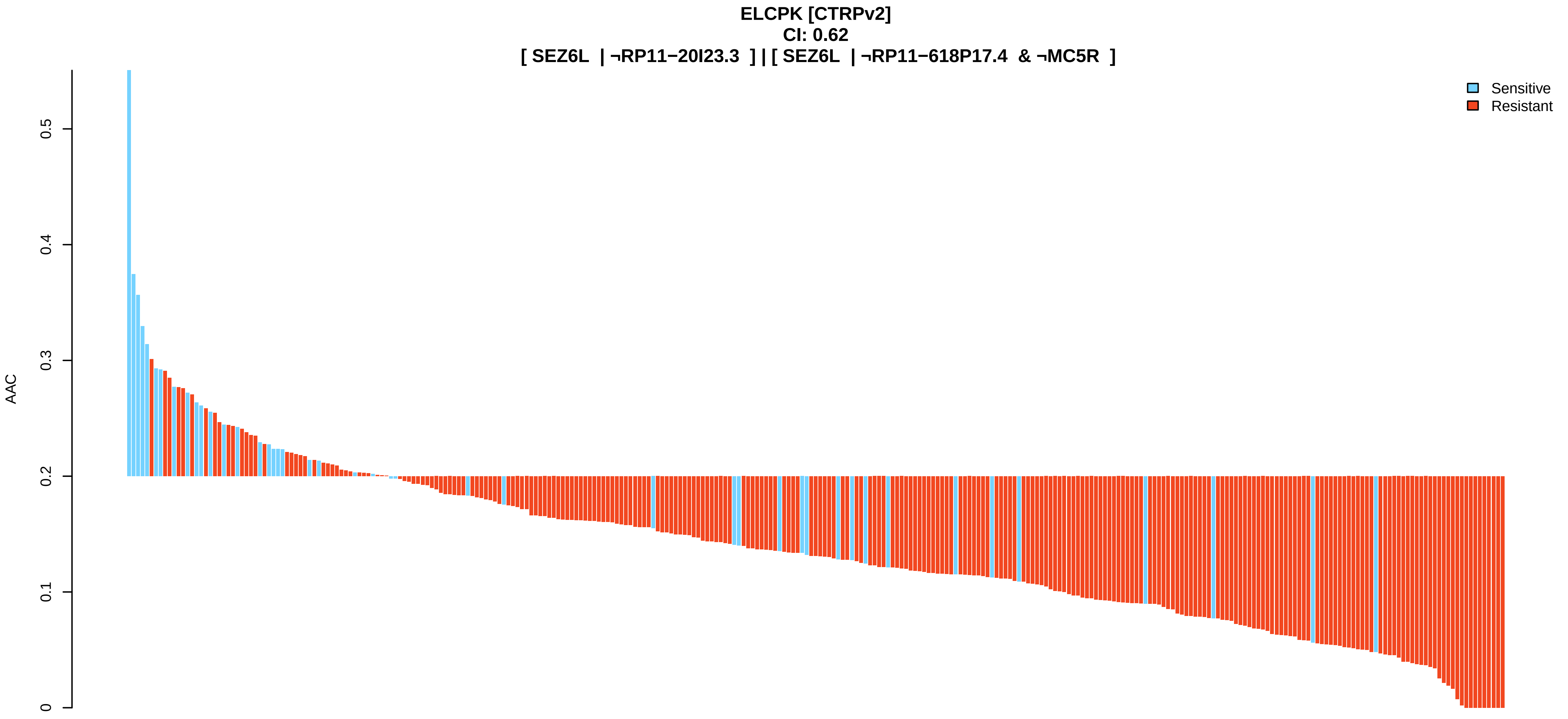
ELCPK [CTRPv2]
CI: 0.62
[ SEZ6L | ¬RP11−20I23.3 ] | [ SEZ6L | ¬RP11−618P17.4 & ¬MC5R ]
Sensitive
Resistant
AAC
0
0.1
0.2
0.3
0.4
0.5

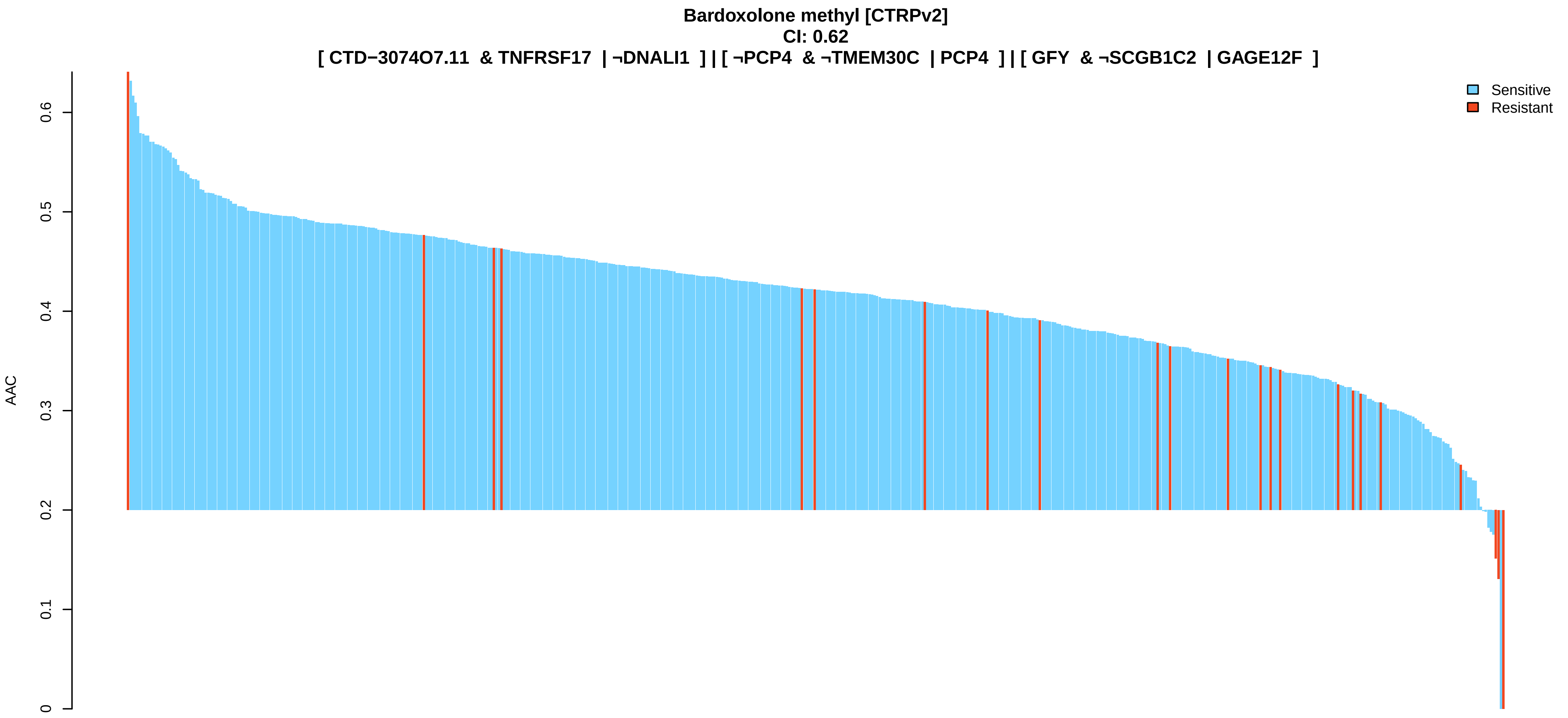

Bardoxolone methyl [CTRPv2]
CI: 0.62
[ CTD−3074O7.11 & TNFRSF17 | ¬DNALI1 ] | [ ¬PCP4 & ¬TMEM30C | PCP4 ] | [ GFY & ¬SCGB1C2 | GAGE12F ]
Sensitive
Resistant
AAC
0
0.1
0.2
0.3
0.4
0.5
0.6


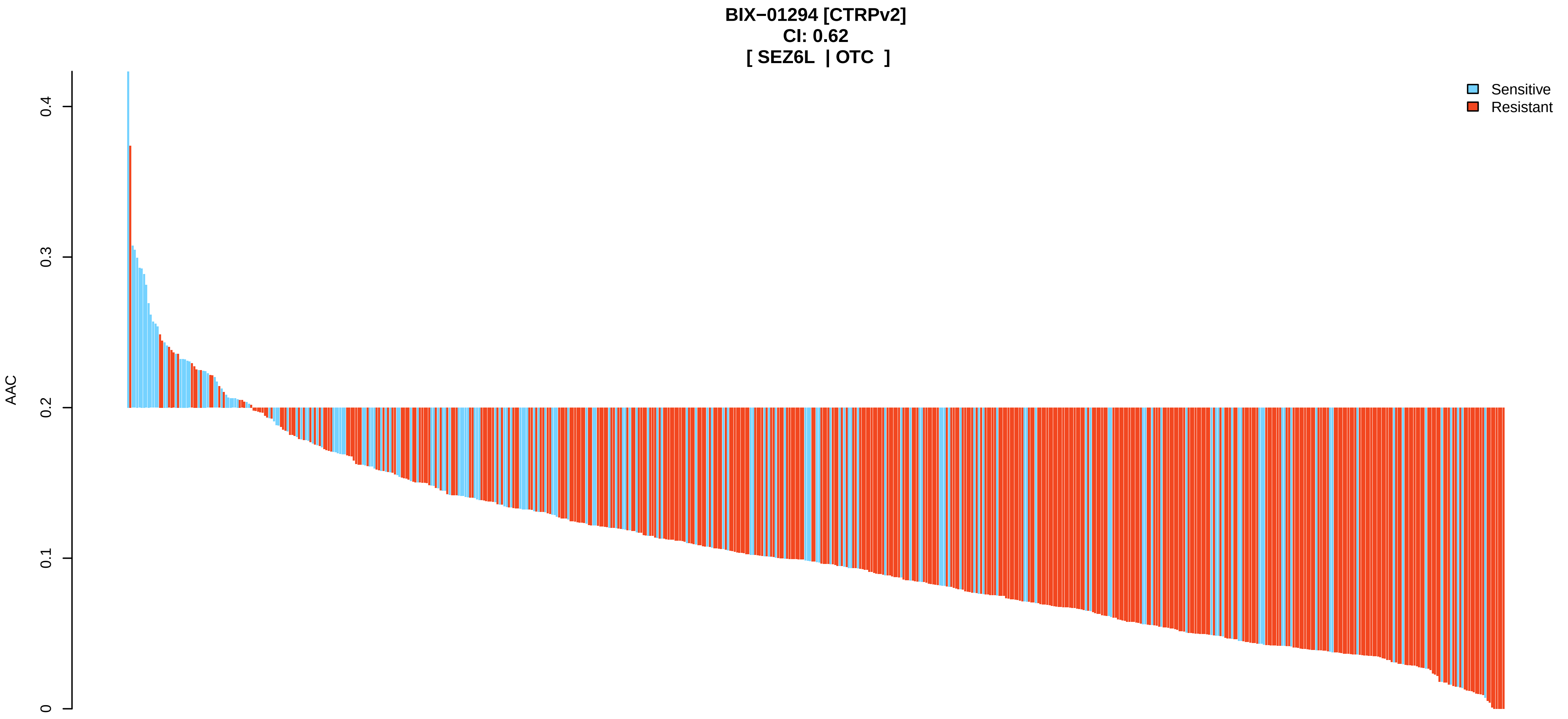

BIX−01294 [CTRPv2]
CI: 0.62
[ SEZ6L | OTC ]
Sensitive
Resistant
AAC
0
0.1
0.2
0.3
0.4


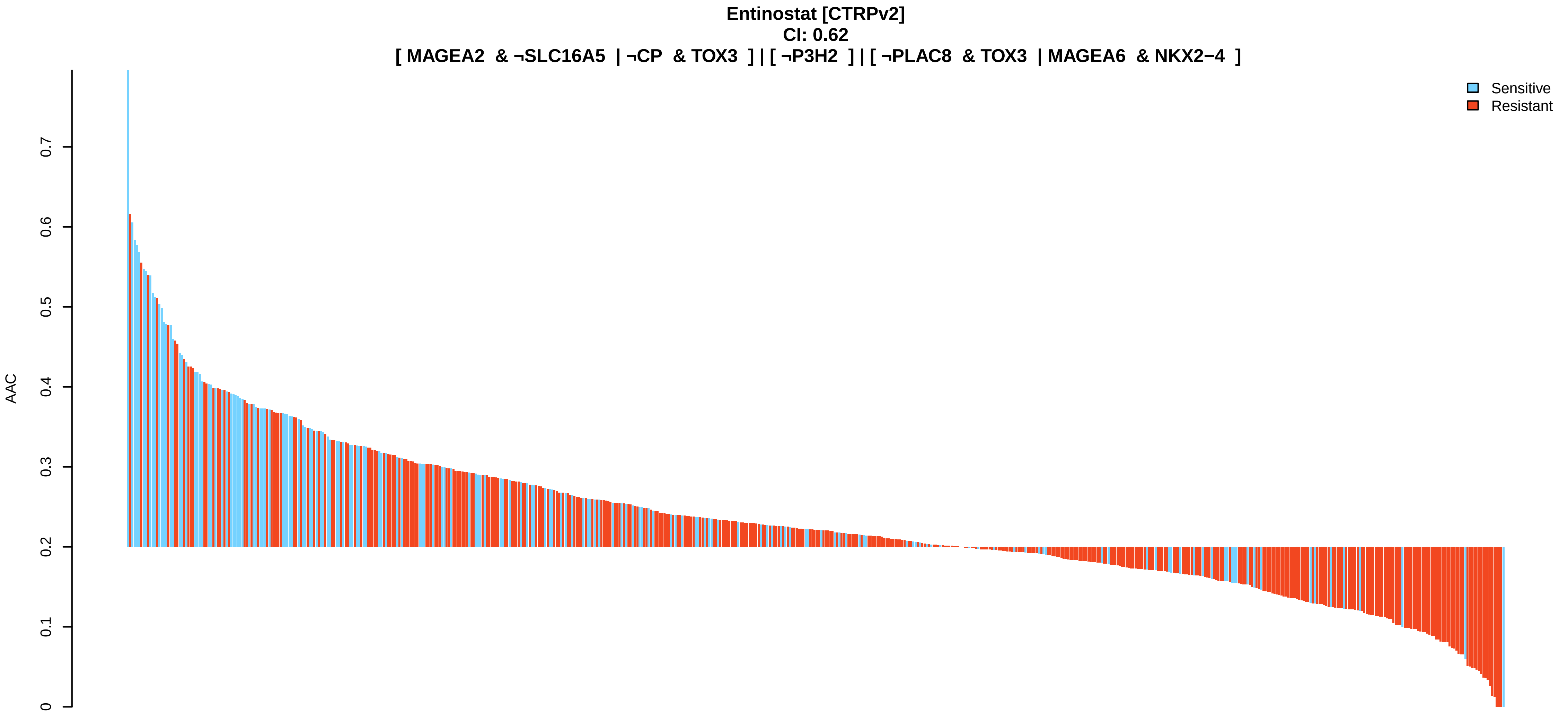

Entinostat [CTRPv2]
CI: 0.62
[ MAGEA2 & ¬SLC16A5 | ¬CP & TOX3 ] | [ ¬P3H2 ] | [ ¬PLAC8 & TOX3 | MAGEA6 & NKX2−4 ]
Sensitive
Resistant
AAC
0
0.1
0.2
0.3
0.4
0.5
0.6
0.7

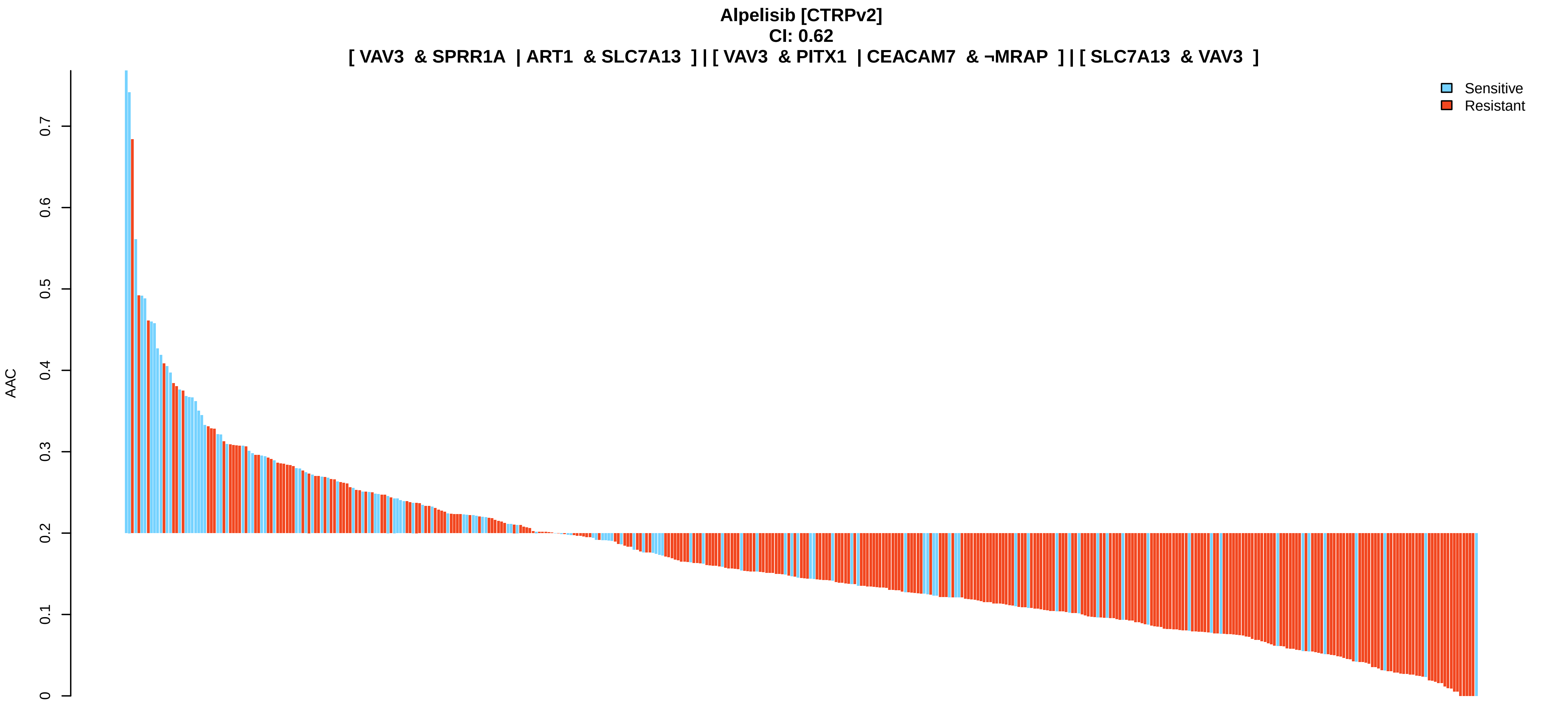
Alpelisib [CTRPv2]
CI: 0.62
[ VAV3 & SPRR1A | ART1 & SLC7A13 ] | [ VAV3 & PITX1 | CEACAM7 & ¬MRAP ] | [ SLC7A13 & VAV3 ]
Sensitive
Resistant
AAC
0
0.1
0.2
0.3
0.4
0.5
0.6
0.7

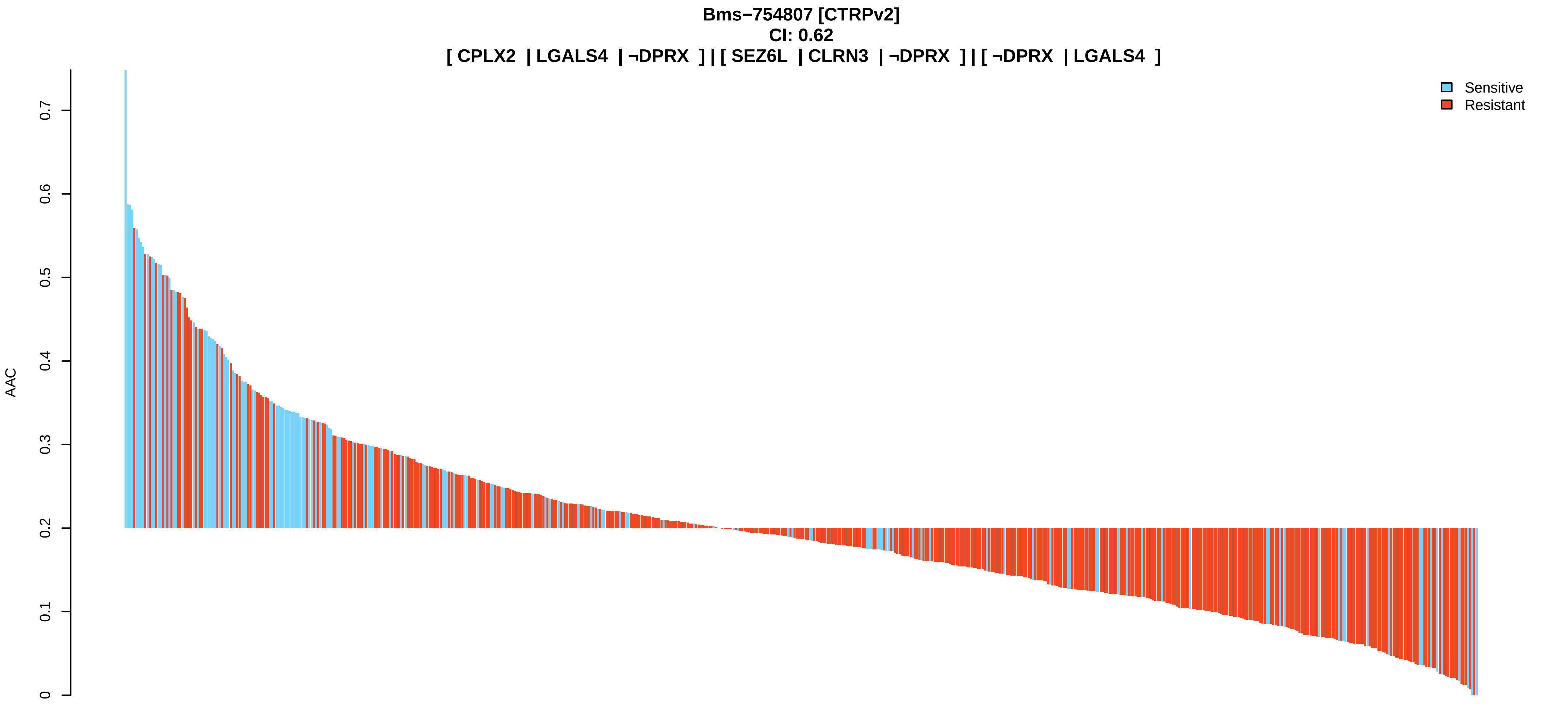
Bms−754807 [CTRPv2]
CI: 0.62
[ CPLX2 | LGALS4 | ¬DPRX ] | [ SEZ6L | CLRN3 | ¬DPRX ] | [ ¬DPRX | LGALS4 ]
Sensitive
Resistant
AAC
0
0.1
0.2
0.3
0.4
0.5
0.6
0.7

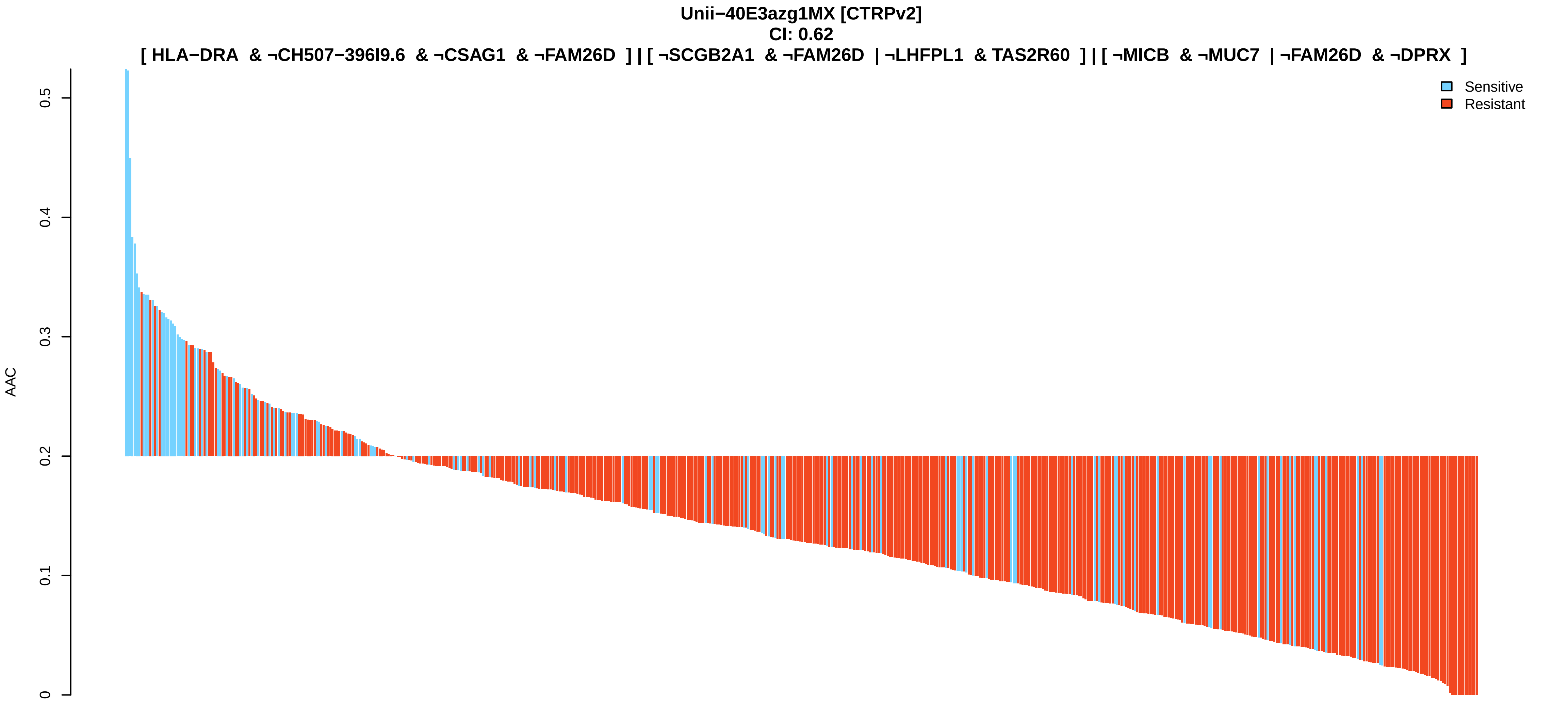
Unii−40E3azg1MX [CTRPv2]
CI: 0.62
[ HLA−DRA & ¬CH507−396I9.6 & ¬CSAG1 & ¬FAM26D ] | [ ¬SCGB2A1 & ¬FAM26D | ¬LHFPL1 & TAS2R60 ] | [ ¬MICB & ¬MUC7 | ¬FAM26D & ¬DPRX ]
Sensitive
Resistant
AAC
0
0.1
0.2
0.3
0.4
0.5

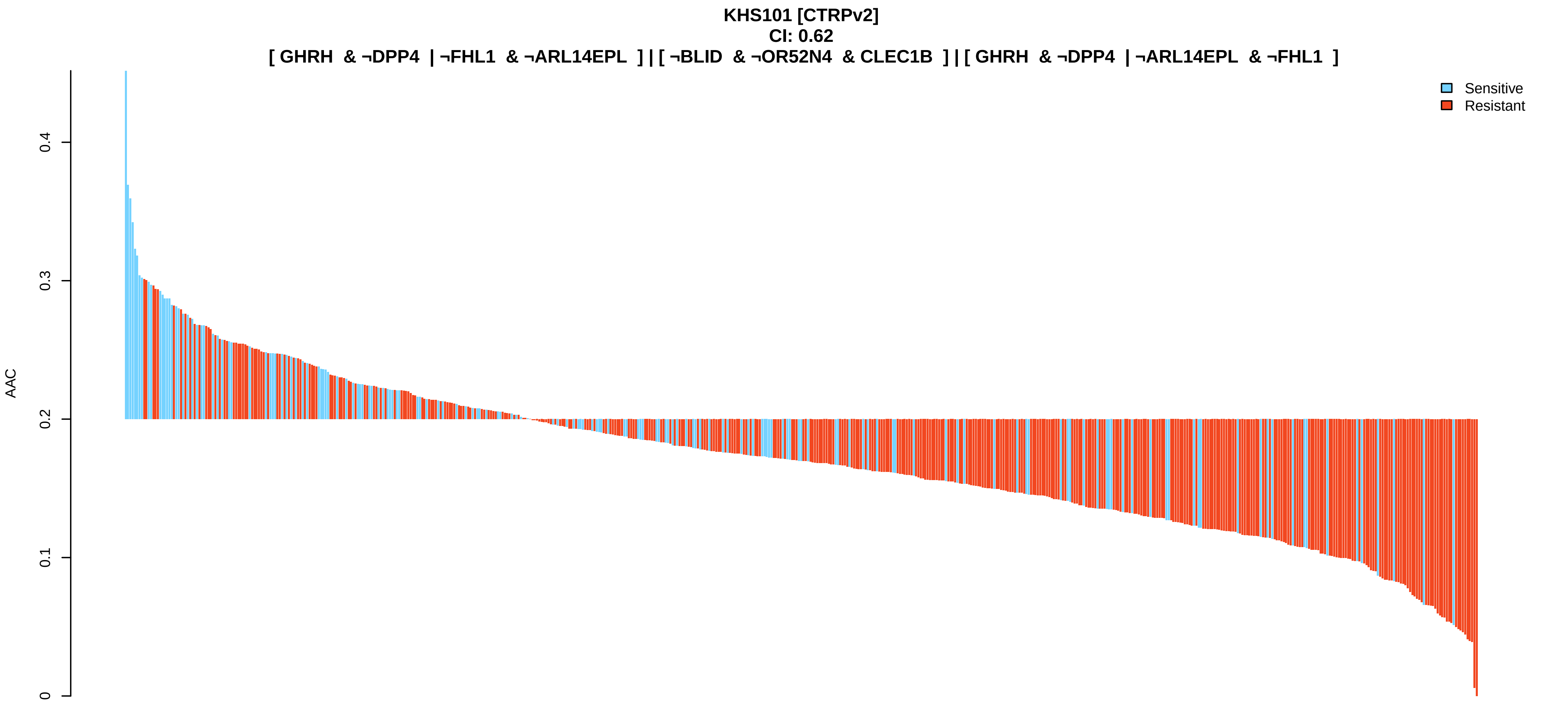
KHS101 [CTRPv2]
CI: 0.62
[ GHRH & ¬DPP4 | ¬FHL1 & ¬ARL14EPL ] | [ ¬BLID & ¬OR52N4 & CLEC1B ] | [ GHRH & ¬DPP4 | ¬ARL14EPL & ¬FHL1 ]
Sensitive
Resistant
AAC
0
0.1
0.2
0.3
0.4

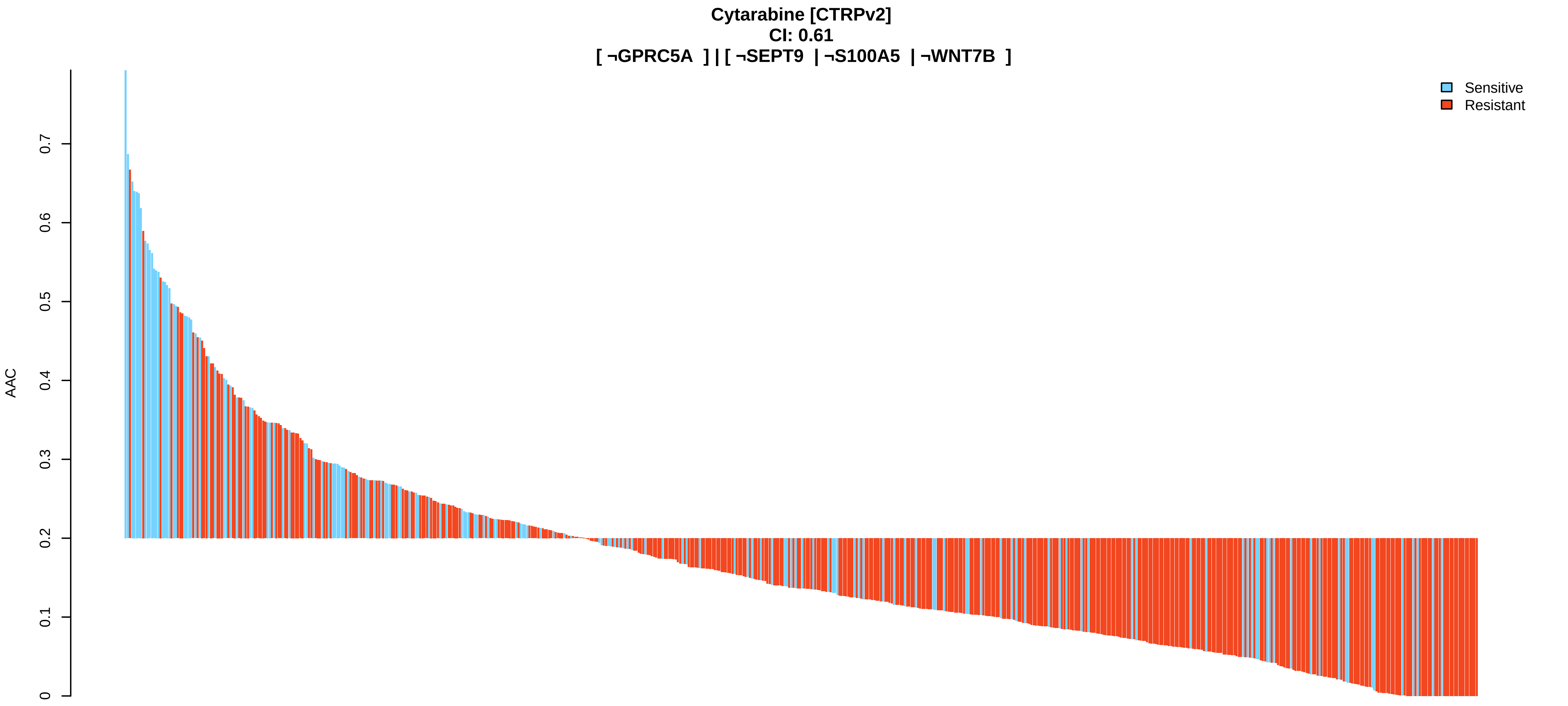
Cytarabine [CTRPv2]
CI: 0.61
[ ¬GPRC5A ] | [ ¬SEPT9 | ¬S100A5 | ¬WNT7B ]
Sensitive
Resistant
AAC
0
0.1
0.2
0.3
0.4
0.5
0.6
0.7

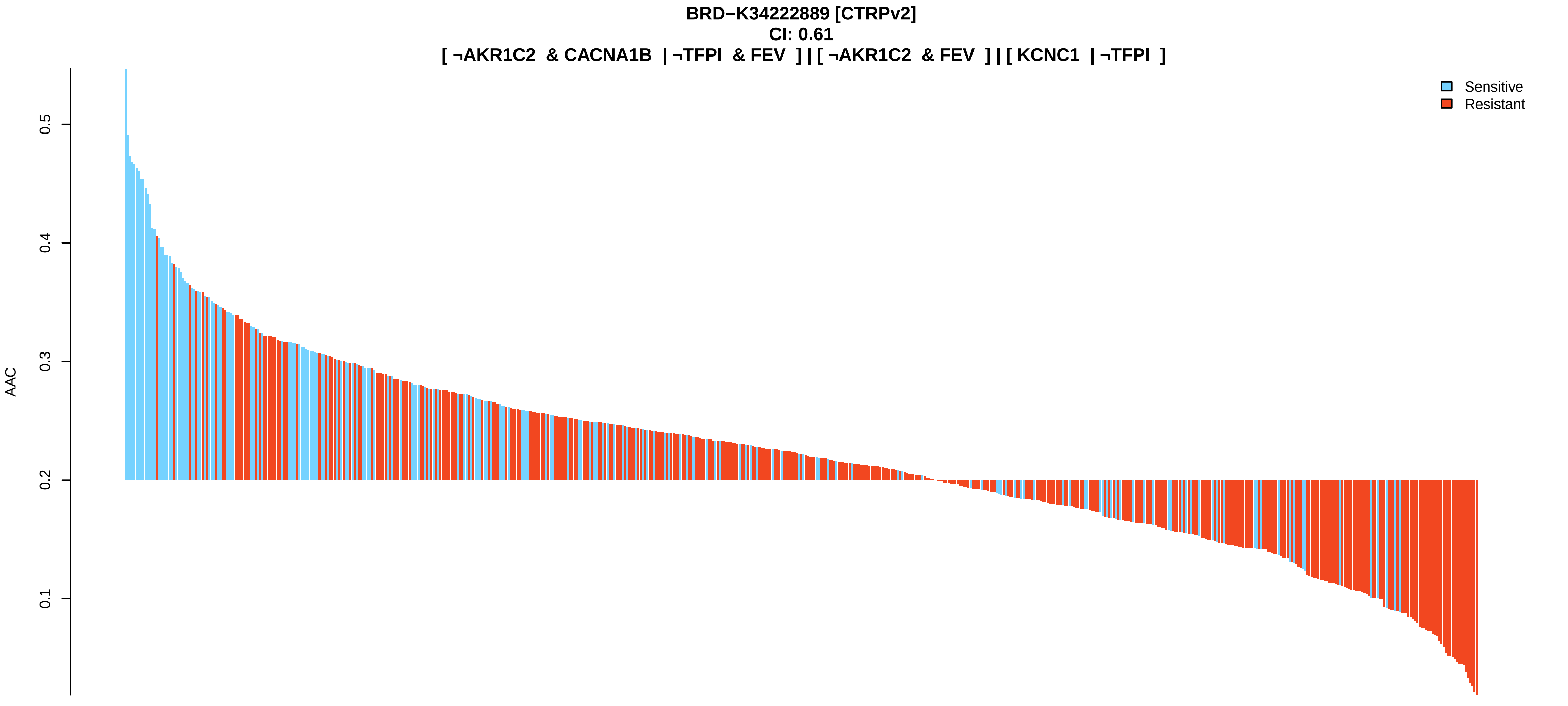
BRD−K34222889 [CTRPv2]
CI: 0.61
[ ¬AKR1C2 & CACNA1B | ¬TFPI & FEV ] | [ ¬AKR1C2 & FEV ] | [ KCNC1 | ¬TFPI ]
Sensitive
Resistant
AAC
0.1
0.2
0.3
0.4
0.5

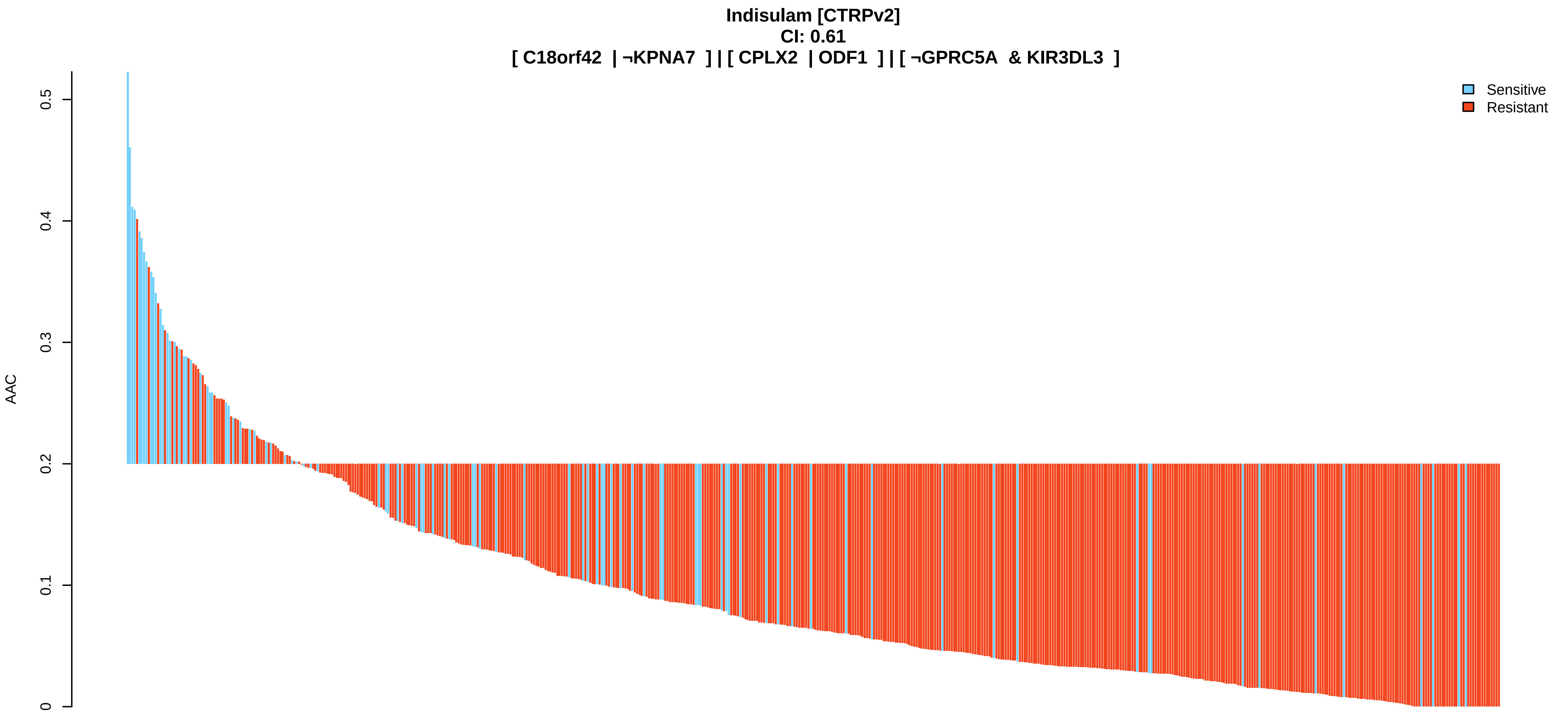
Indisulam [CTRPv2]
CI: 0.61
[ C18orf42 | ¬KPNA7 ] | [ CPLX2 | ODF1 ] | [ ¬GPRC5A & KIR3DL3 ]
AAC
0
0.1
0.2
0.3
0.4
0.5
Sensitive
Resistant

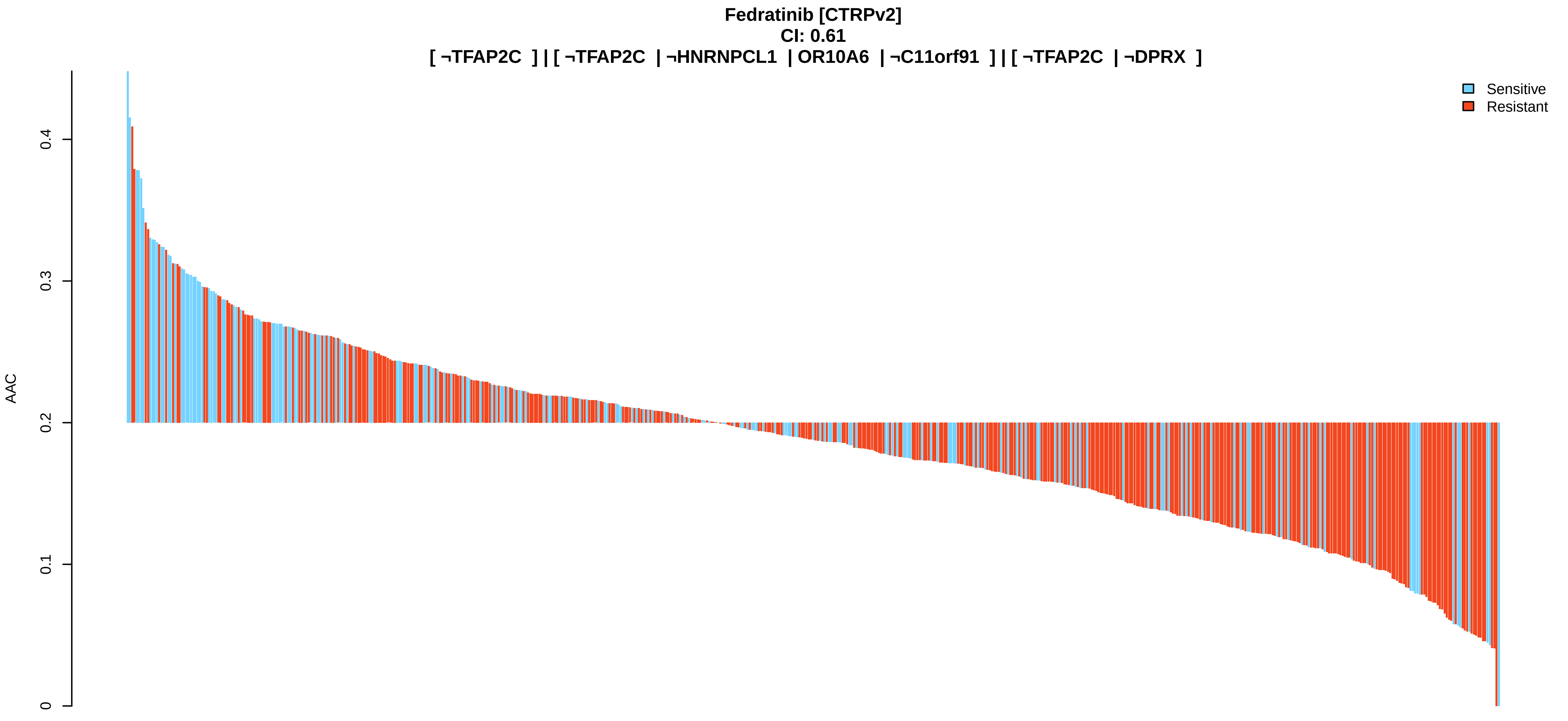
Fedratinib [CTRPv2]
CI: 0.61
[ ¬TFAP2C ] | [ ¬TFAP2C | ¬HNRNPCL1 | OR10A6 | ¬C11orf91 ] | [ ¬TFAP2C | ¬DPRX ]
AAC
0
0.1
0.2
0.3
0.4
Sensitive
Resistant

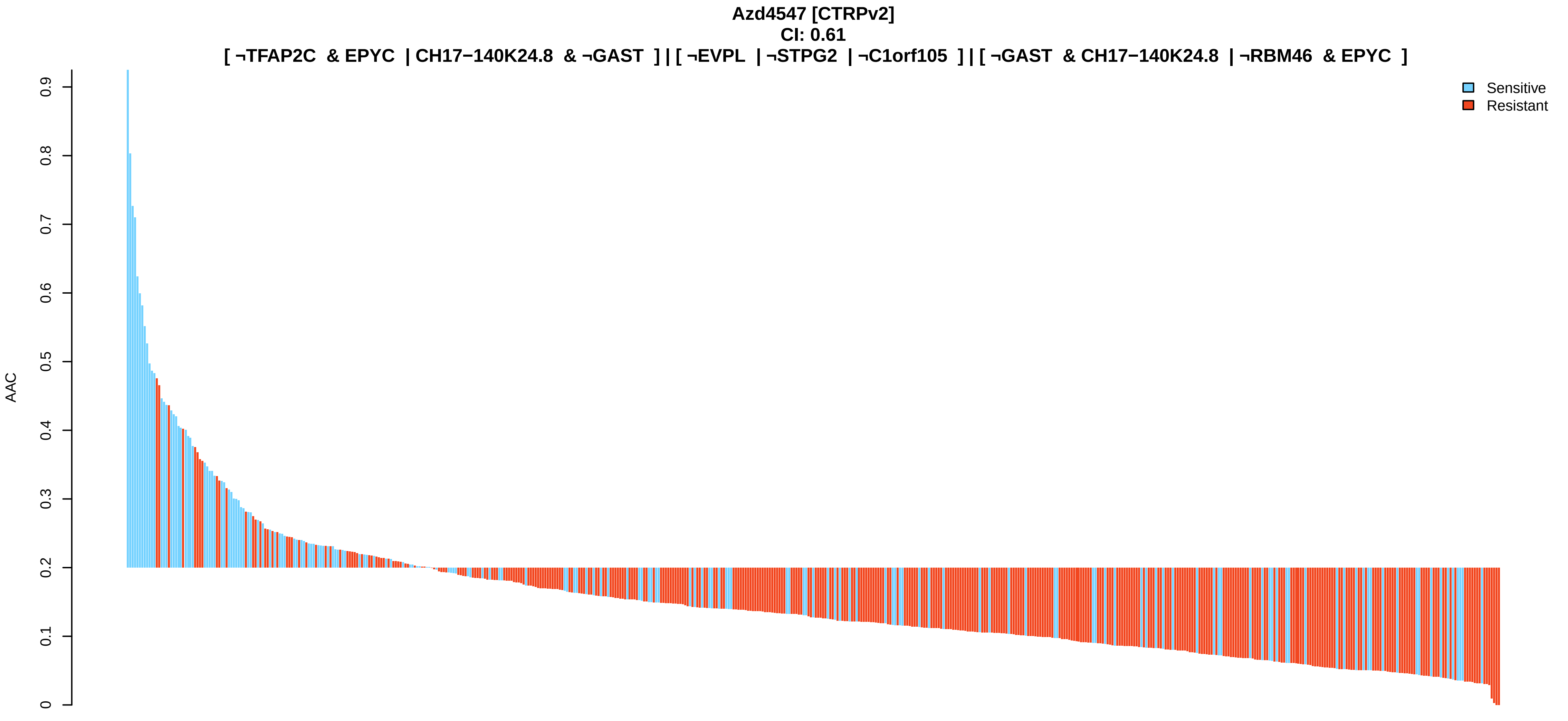
Azd4547 [CTRPv2]
CI: 0.61
[ ¬TFAP2C & EPYC | CH17−140K24.8 & ¬GAST ] | [ ¬EVPL | ¬STPG2 | ¬C1orf105 ] | [ ¬GAST & CH17−140K24.8 | ¬RBM46 & EPYC ]
AAC
0
0.1
0.2
0.3
0.4
0.5
0.6
0.7
0.8
0.9
Sensitive
Resistant

**BRD−K35604418 [CTRPv2]**
**CI: 0.61**
**[ SLC7A10 & ¬HLA−E ] | [ ¬TINAGL1 & MKRN3 | TRIM27 & DPCR1 ] | [ ¬C9orf57 & IBSP & RP11−834C11.12 ]**

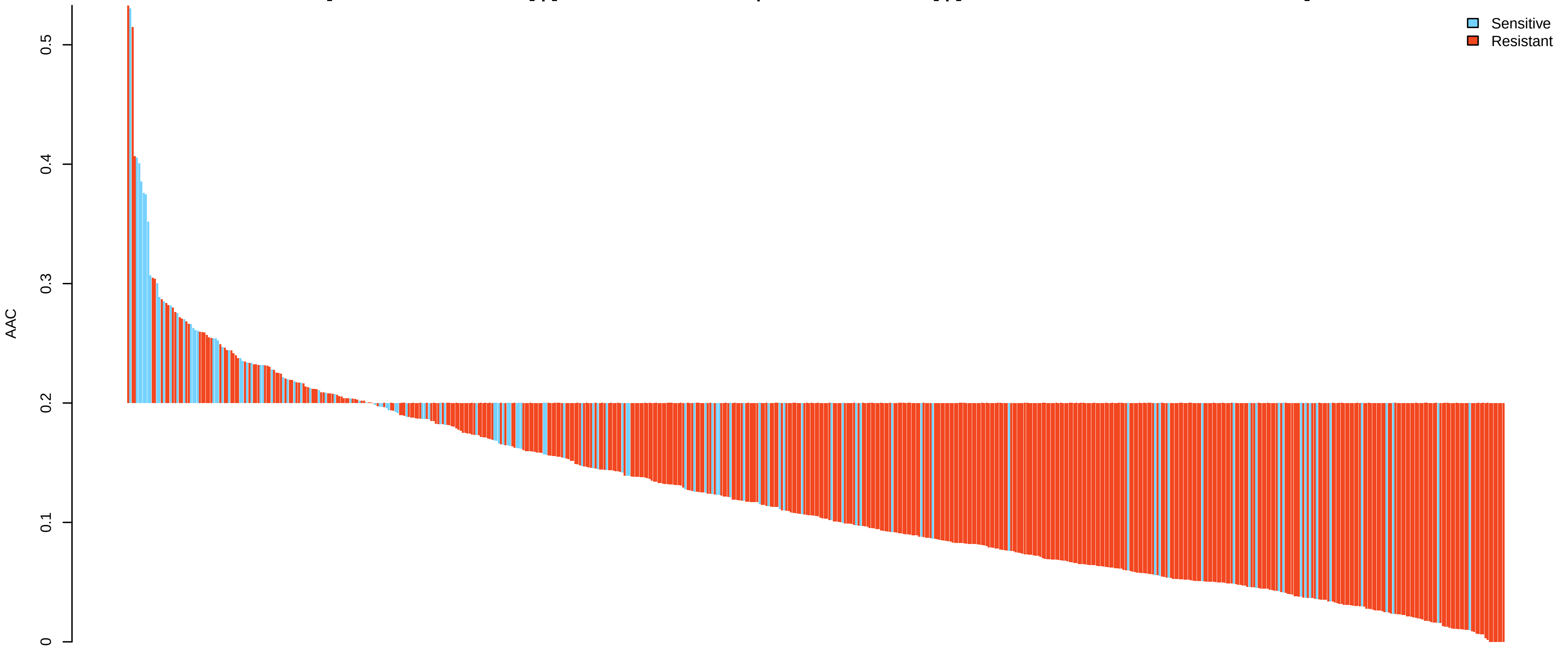


**STF−31 [CTRPv2]**
**CI: 0.61**
**[ ¬LIPH & ¬ZNF311 | ¬LIPH & ¬CST2 ] | [ ¬LIPH & ¬CST2 | ¬LIPH & ¬ZNF311 ] | [ ¬OR2A25 & ¬IL36A | CACNA1B & ¬ABCC3 ]**

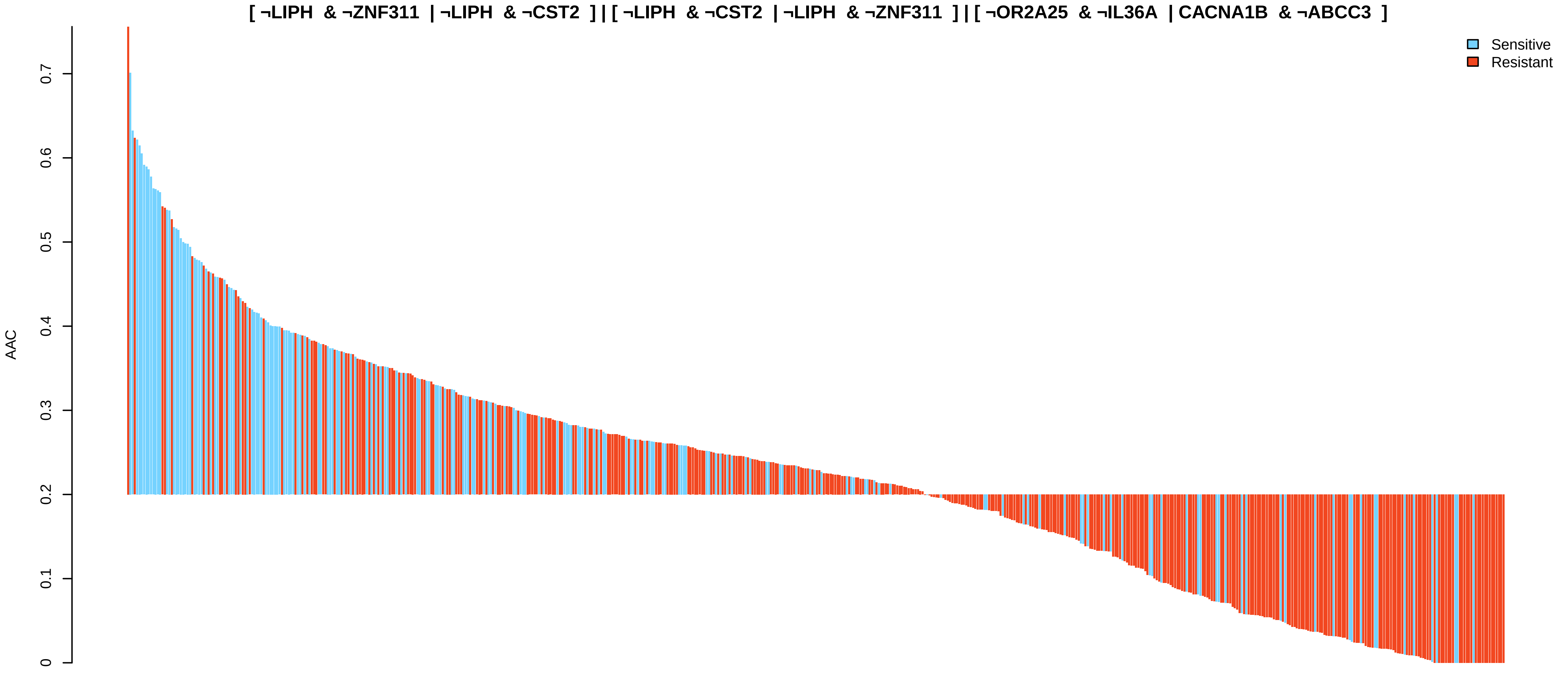


**Cediranib [CTRPv2]**
**CI: 0.61**
**[ ¬GPRC5A | ¬RP11−637O19.3 ] | [ UTS2 & TUBB ] | [ ¬GPRC5A & ¬PPP1R11 | ¬PSORS1C1 & GPR25 ]**

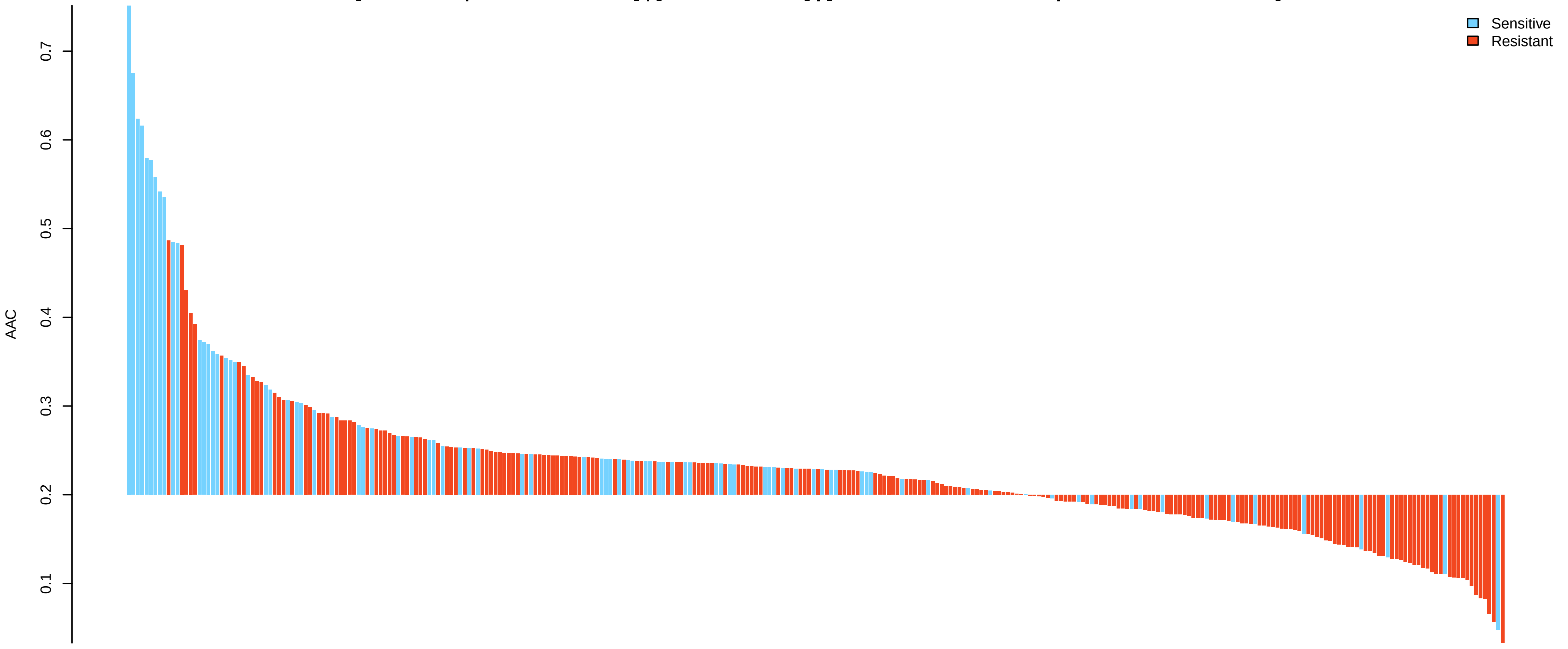

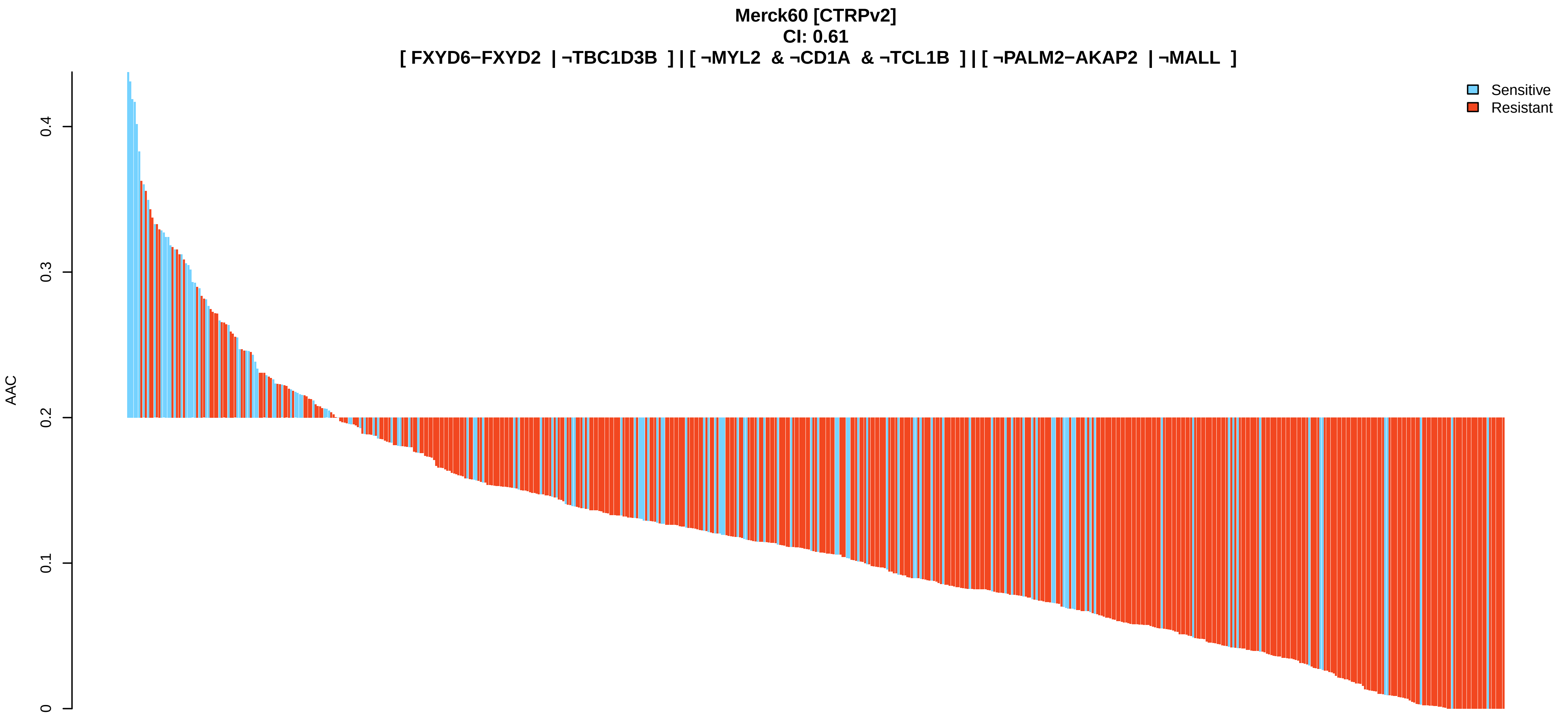
Merck60 [CTRPv2]
CI: 0.61
[ FXYD6−FXYD2 | ¬TBC1D3B ] | [ ¬MYL2 & ¬CD1A & ¬TCL1B ] | [ ¬PALM2−AKAP2 | ¬MALL ]
Sensitive
Resistant
AAC
0
0.1
0.2
0.3
0.4

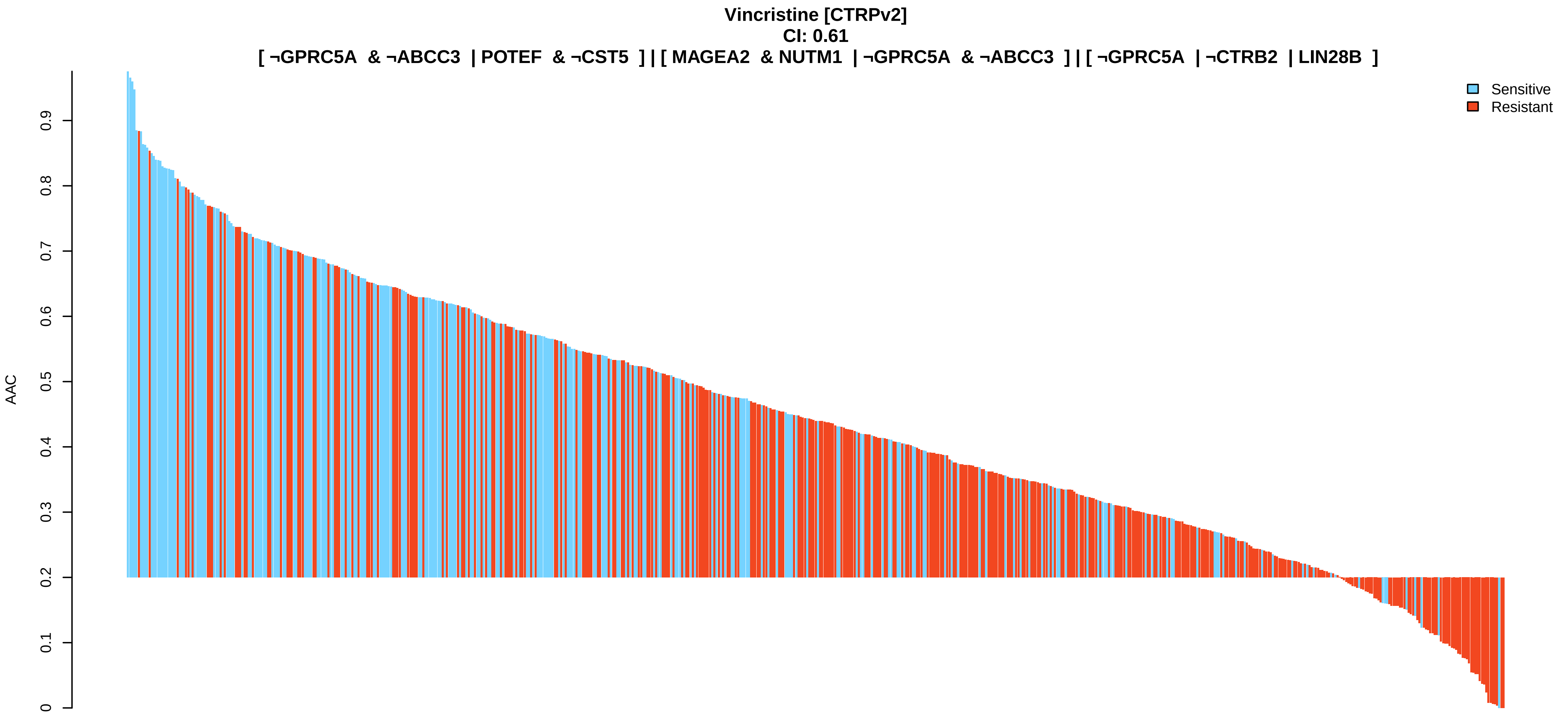
Vincristine [CTRPv2]
CI: 0.61
[ ¬GPRC5A & ¬ABCC3 | POTEF & ¬CST5 ] | [ MAGEA2 & NUTM1 | ¬GPRC5A & ¬ABCC3 ] | [ ¬GPRC5A | ¬CTRB2 | LIN28B ]
Sensitive
Resistant
AAC
0
0.1
0.2
0.3
0.4
0.5
0.6
0.7
0.8
0.9

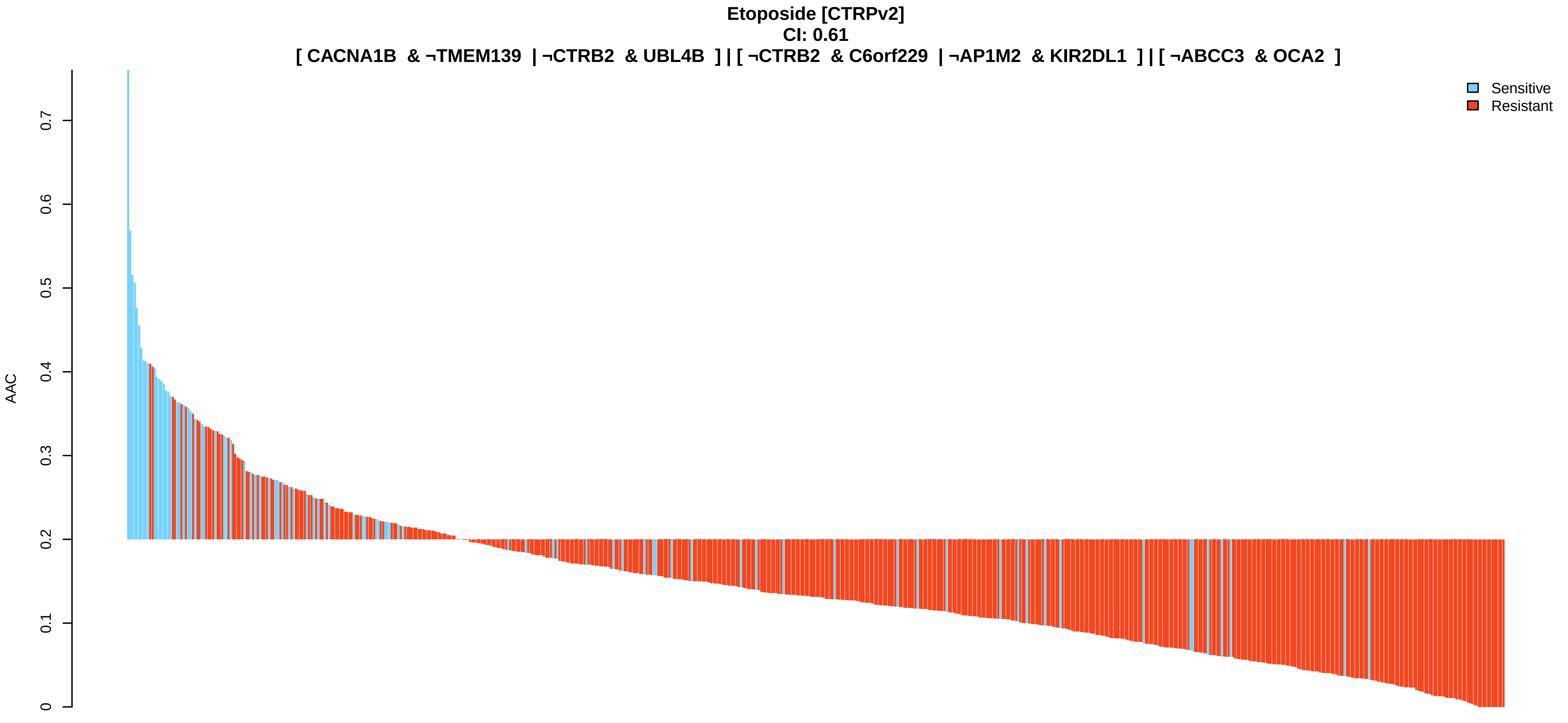
Etoposide [CTRPv2]
CI: 0.61
[ CACNA1B & ¬TMEM139 | ¬CTRB2 & UBL4B ] | [ ¬CTRB2 & C6orf229 | ¬AP1M2 & KIR2DL1 ] | [ ¬ABCC3 & OCA2 ]
Sensitive
Resistant
AAC
0
0.1
0.2
0.3
0.4
0.5
0.6
0.7

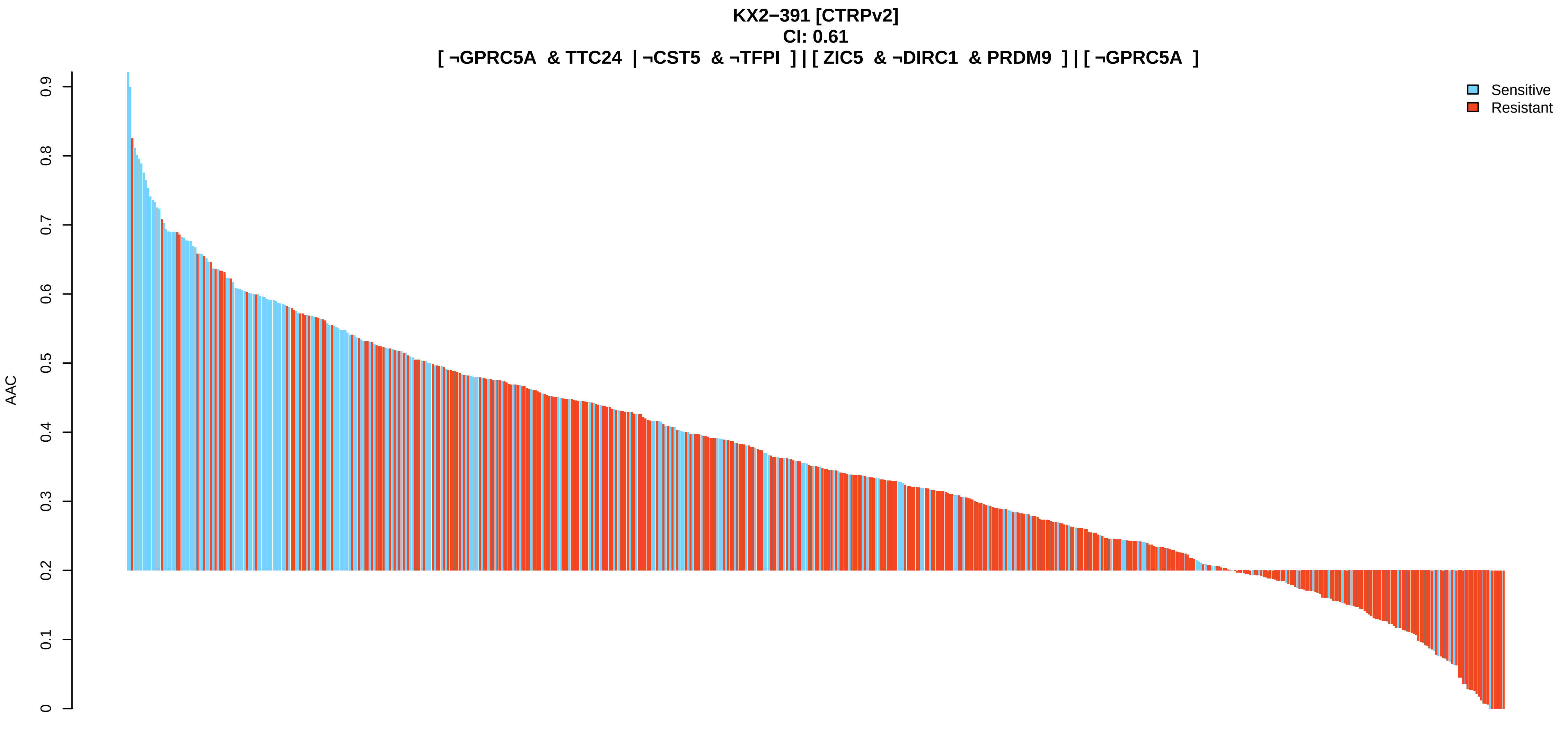
KX2−391 [CTRPv2]
CI: 0.61
[ ¬GPRC5A & TTC24 | ¬CST5 & ¬TFPI ] | [ ZIC5 & ¬DIRC1 & PRDM9 ] | [ ¬GPRC5A ]
AAC
0
0.1
0.2
0.3
0.4
0.5
0.6
0.7
0.8
0.9
Sensitive
Resistant

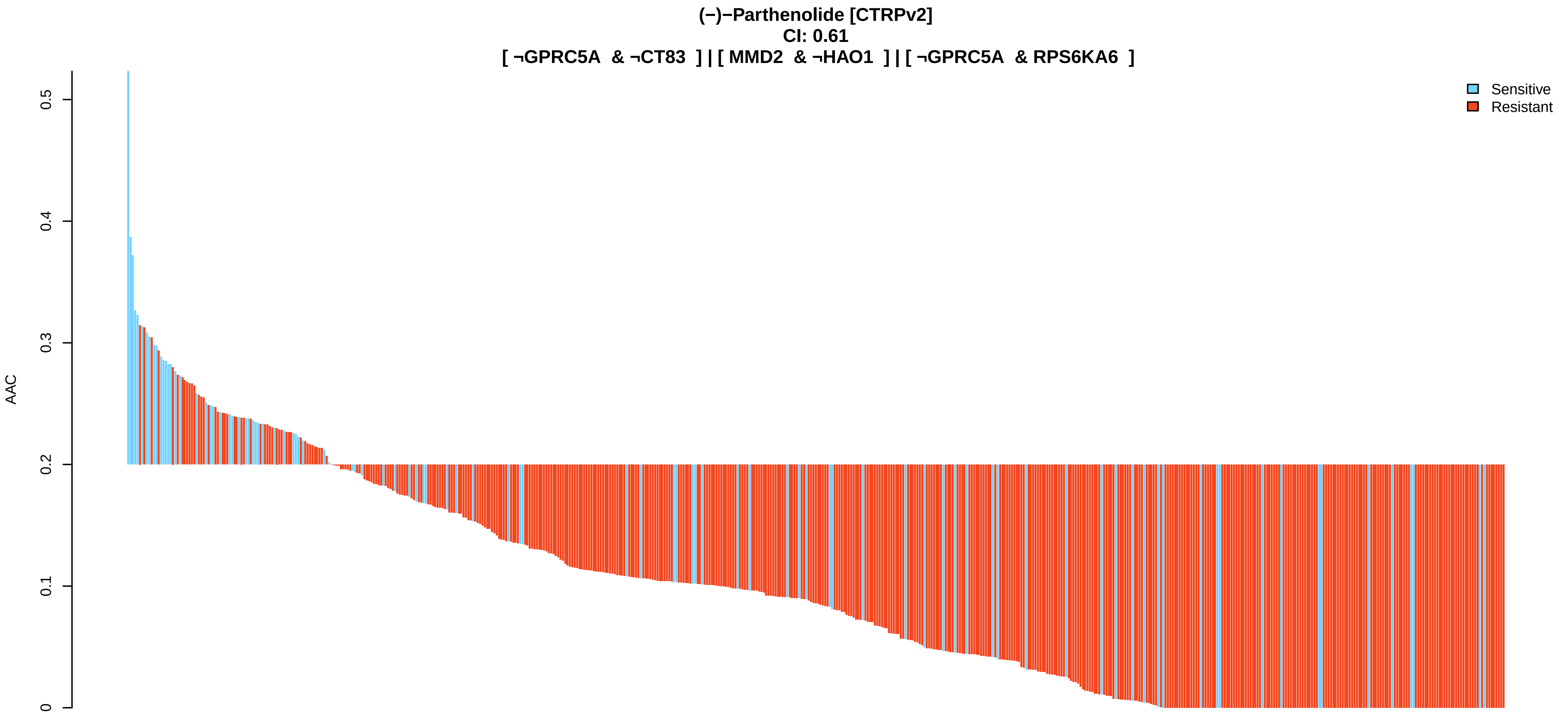
(−)−Parthenolide [CTRPv2]
CI: 0.61
[ ¬GPRC5A & ¬CT83 ] | [ MMD2 & ¬HAO1 ] | [ ¬GPRC5A & RPS6KA6 ]
AAC
0
0.1
0.2
0.3
0.4
0.5
Sensitive
Resistant

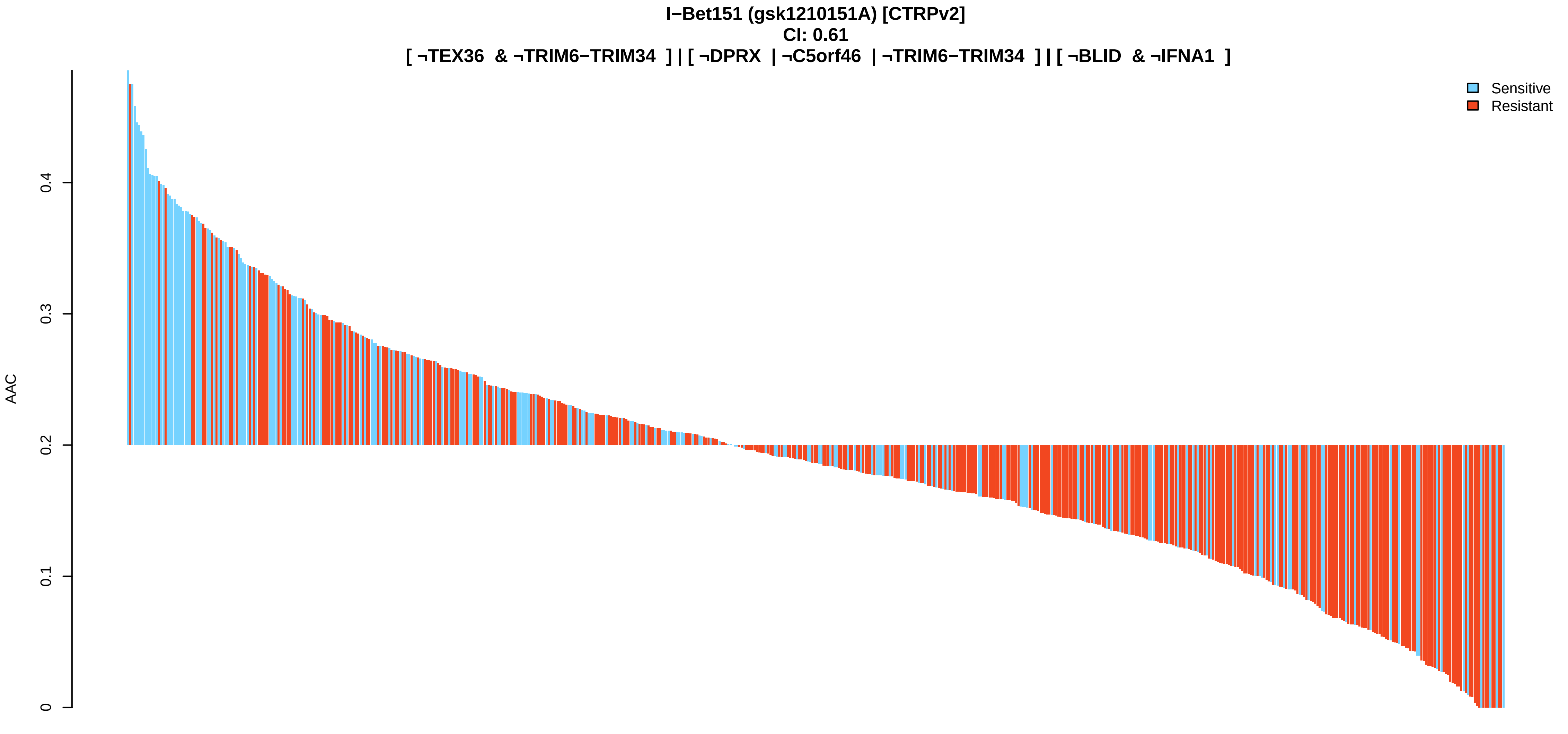
I−Bet151 (gsk1210151A) [CTRPv2]
CI: 0.61
[ ¬TEX36 & ¬TRIM6−TRIM34 ] | [ ¬DPRX | ¬C5orf46 | ¬TRIM6−TRIM34 ] | [ ¬BLID & ¬IFNA1 ]
AAC
0
0.1
0.2
0.3
0.4
Sensitive
Resistant

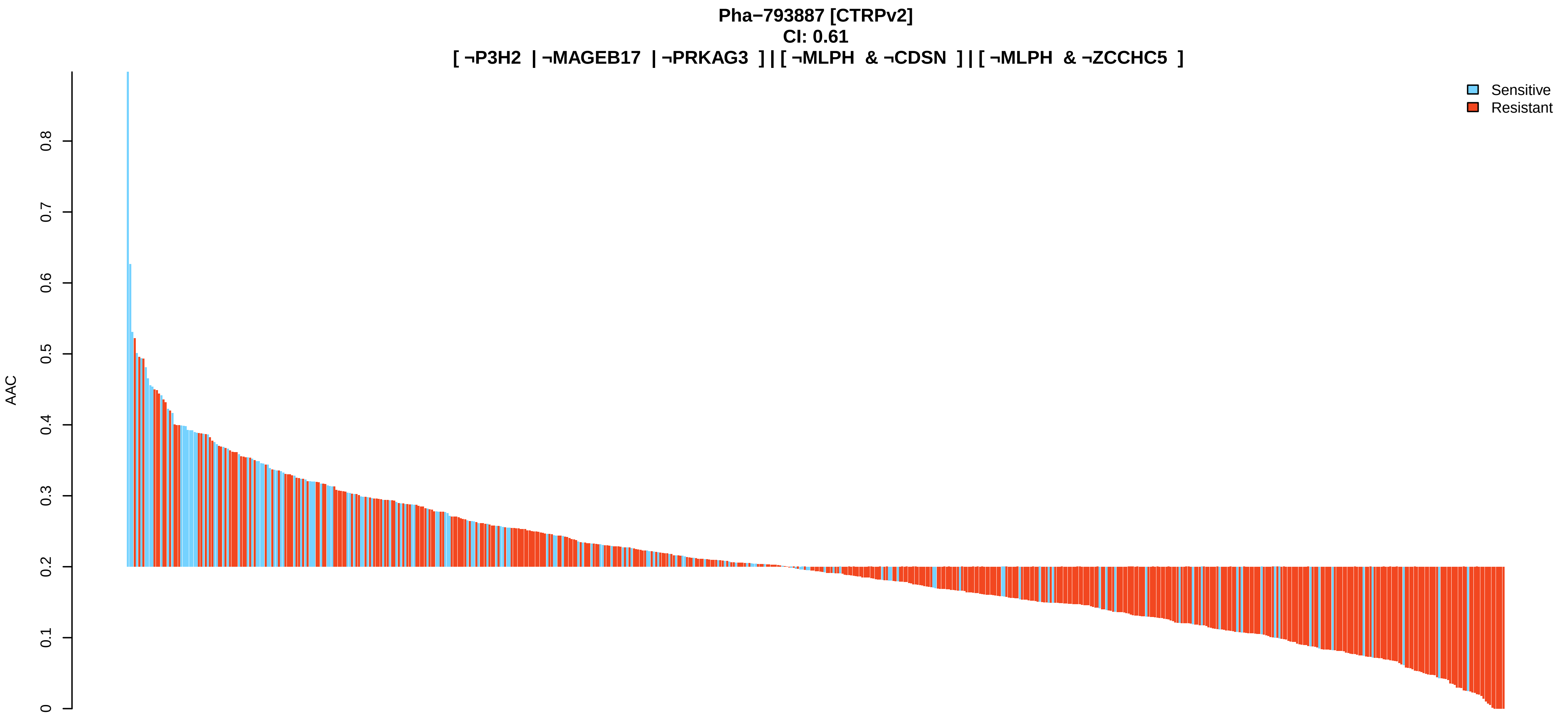
Pha−793887 [CTRPv2]
CI: 0.61
[ ¬P3H2 | ¬MAGEB17 | ¬PRKAG3 ] | [ ¬MLPH & ¬CDSN ] | [ ¬MLPH & ¬ZCCHC5 ]
AAC
0
0.1
0.2
0.3
0.4
0.5
0.6
0.7
0.8
Sensitive
Resistant

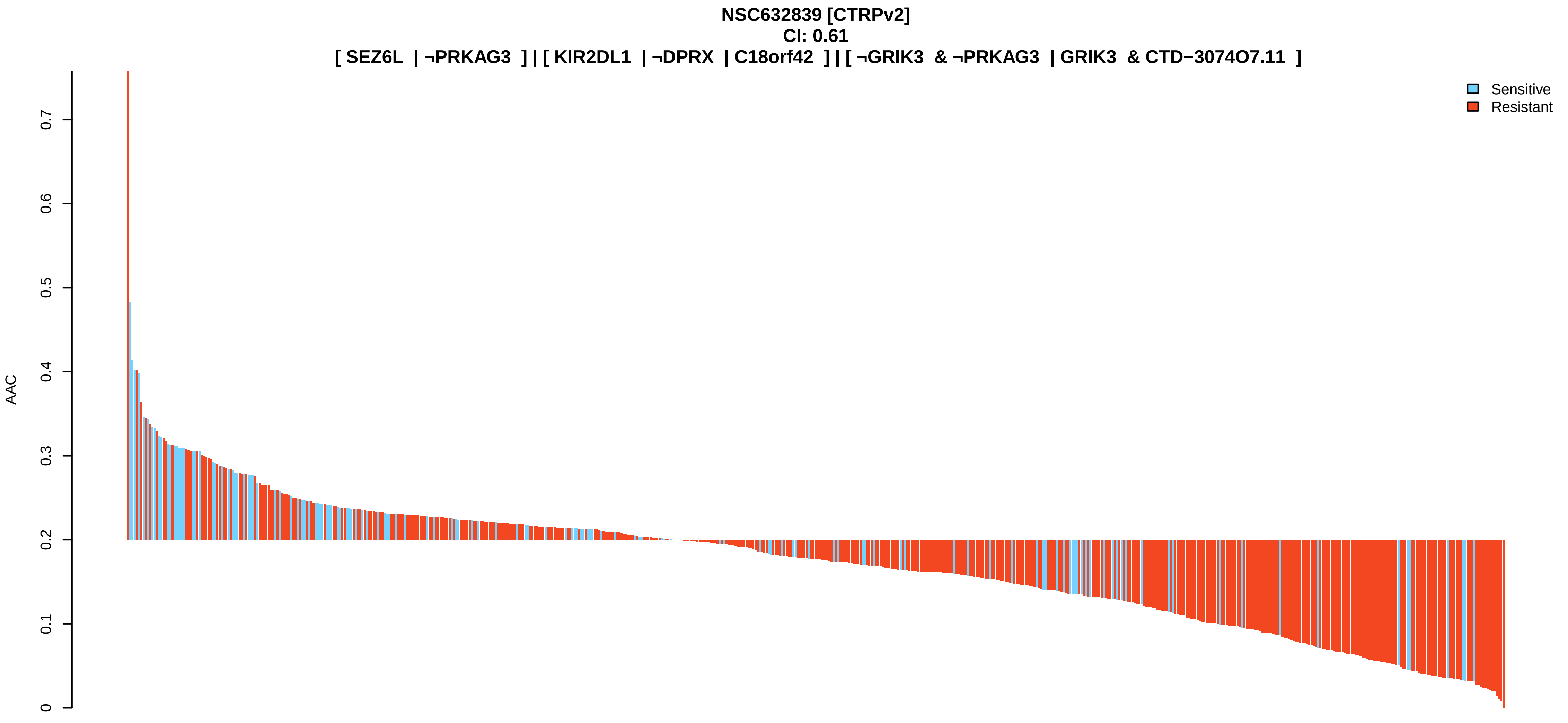
NSC632839 [CTRPv2]
CI: 0.61
[ SEZ6L | ¬PRKAG3 ] | [ KIR2DL1 | ¬DPRX | C18orf42 ] | [ ¬GRIK3 & ¬PRKAG3 | GRIK3 & CTD−3074O7.11 ]
AAC
0
0.1
0.2
0.3
0.4
0.5
0.6
0.7
Sensitive
Resistant

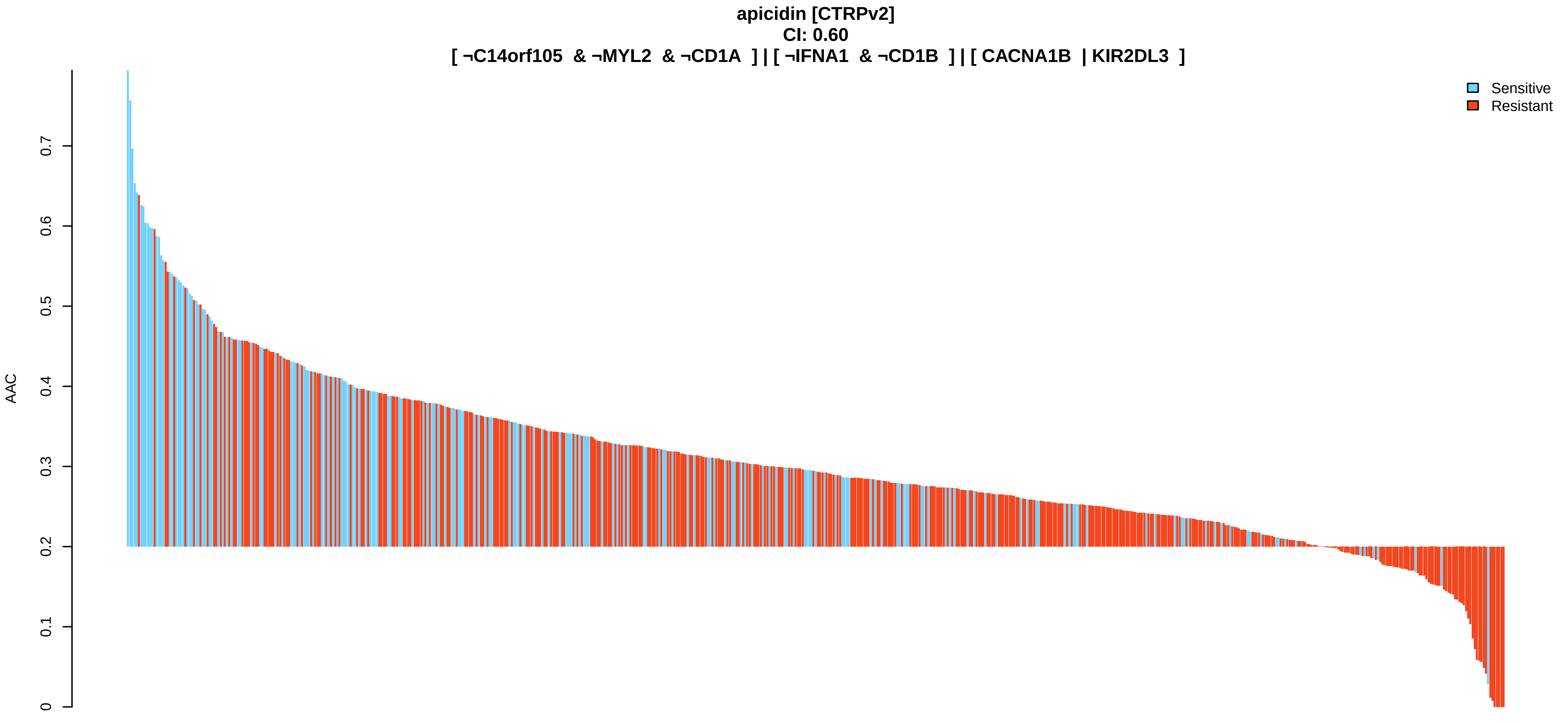
apicidin [CTRPv2]
CI: 0.60
[ ¬C14orf105 & ¬MYL2 & ¬CD1A ] | [ ¬IFNA1 & ¬CD1B ] | [ CACNA1B | KIR2DL3 ]
AAC
0
0.1
0.2
0.3
0.4
0.5
0.6
0.7
Sensitive
Resistant

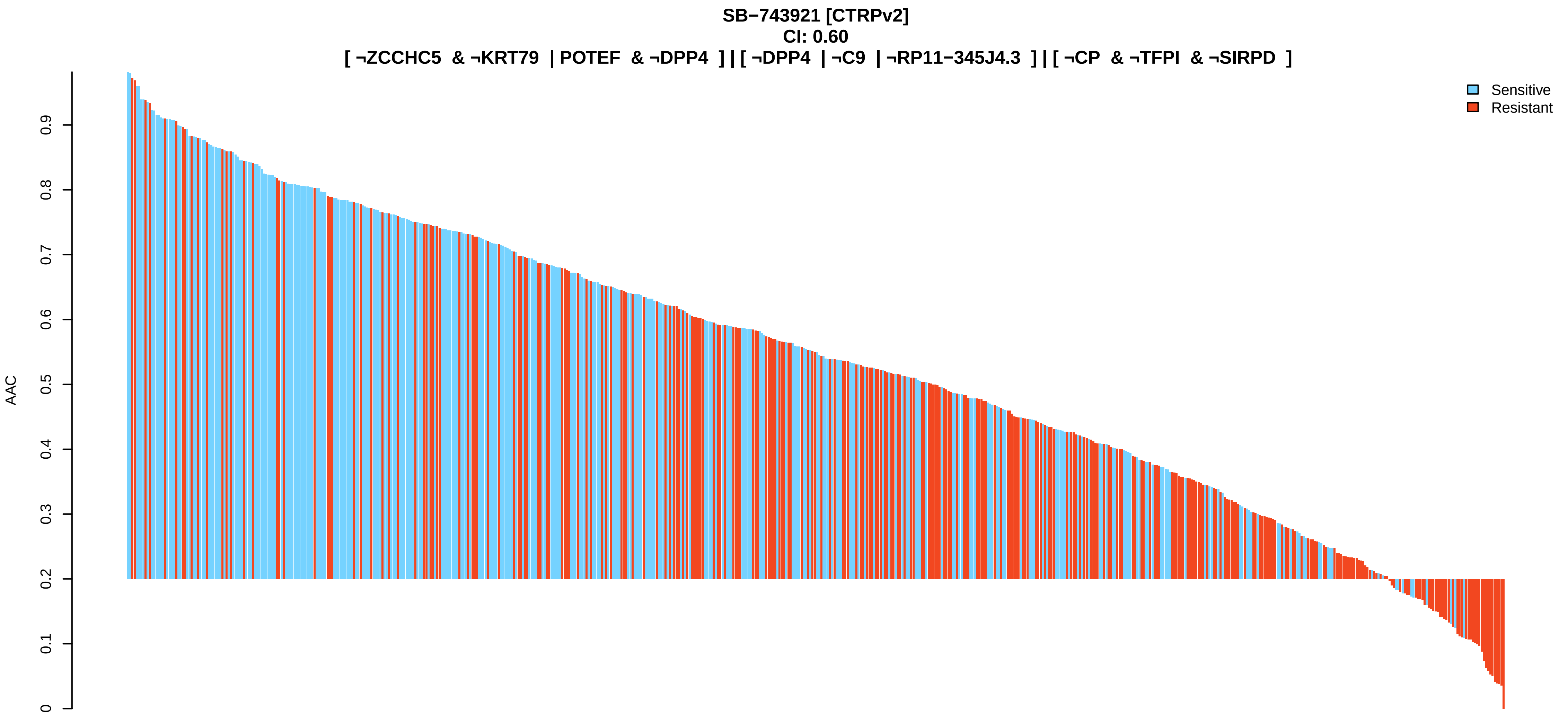
SB−743921 [CTRPv2]
CI: 0.60
[ ¬ZCCHC5 & ¬KRT79 | POTEF & ¬DPP4 ] | [ ¬DPP4 | ¬C9 | ¬RP11−345J4.3 ] | [ ¬CP & ¬TFPI & ¬SIRPD ]
Sensitive
Resistant
AAC
0
0.1
0.2
0.3
0.4
0.5
0.6
0.7
0.8
0.9

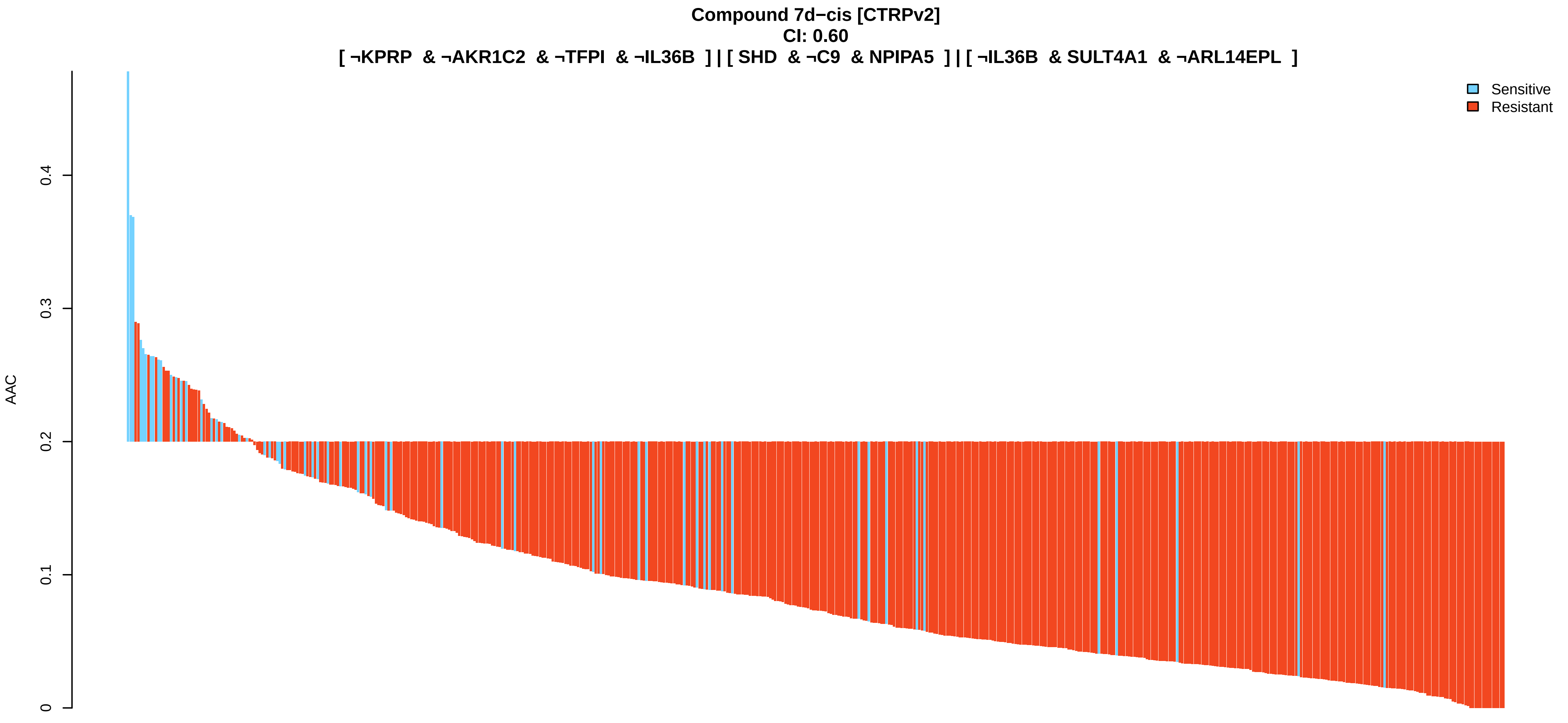
Compound 7d−cis [CTRPv2]
CI: 0.60
[ ¬KPRP & ¬AKR1C2 & ¬TFPI & ¬IL36B ] | [ SHD & ¬C9 & NPIPA5 ] | [ ¬IL36B & SULT4A1 & ¬ARL14EPL ]
Sensitive
Resistant
AAC
0
0.1
0.2
0.3
0.4

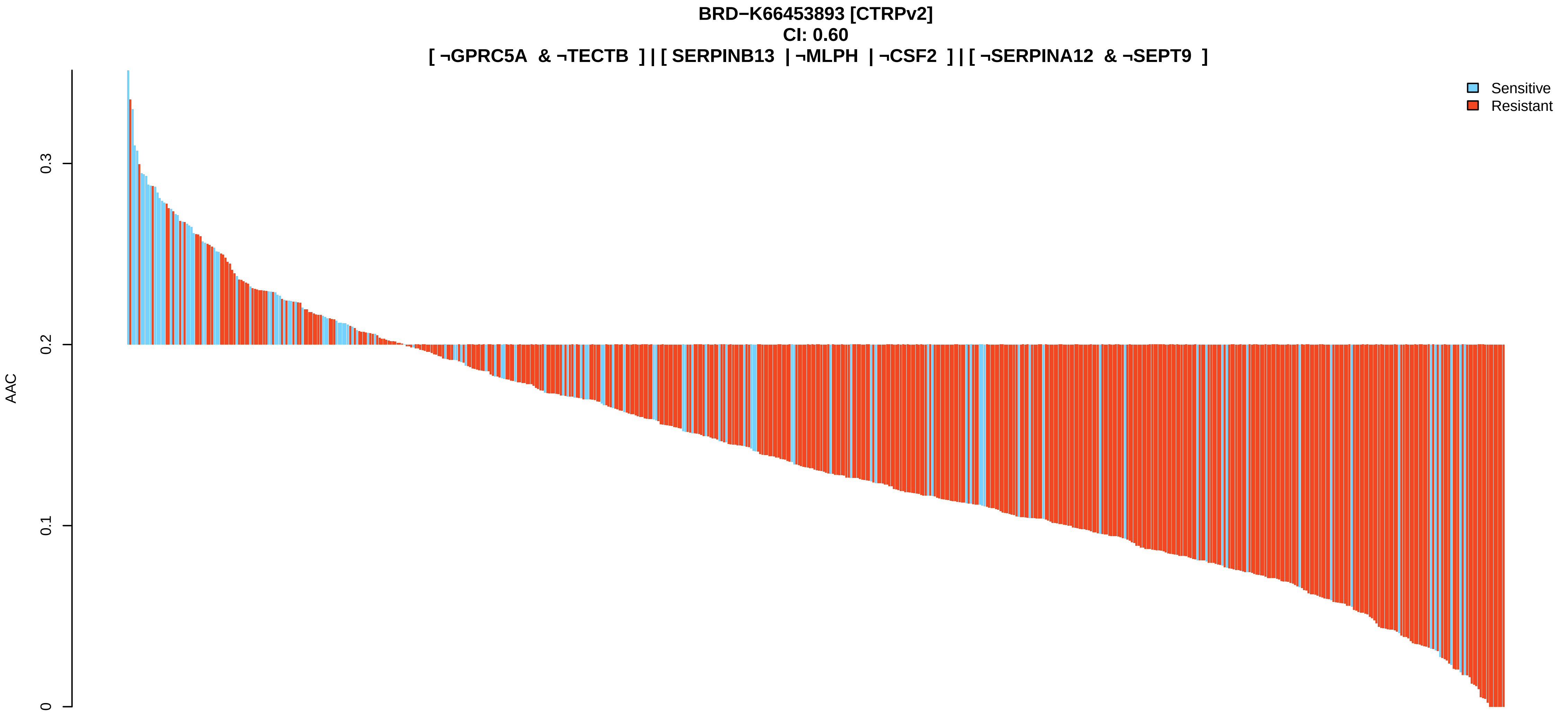
BRD−K66453893 [CTRPv2]
CI: 0.60
[ ¬GPRC5A & ¬TECTB ] | [ SERPINB13 | ¬MLPH | ¬CSF2 ] | [ ¬SERPINA12 & ¬SEPT9 ]
Sensitive
Resistant
AAC
0
0.1
0.2
0.3

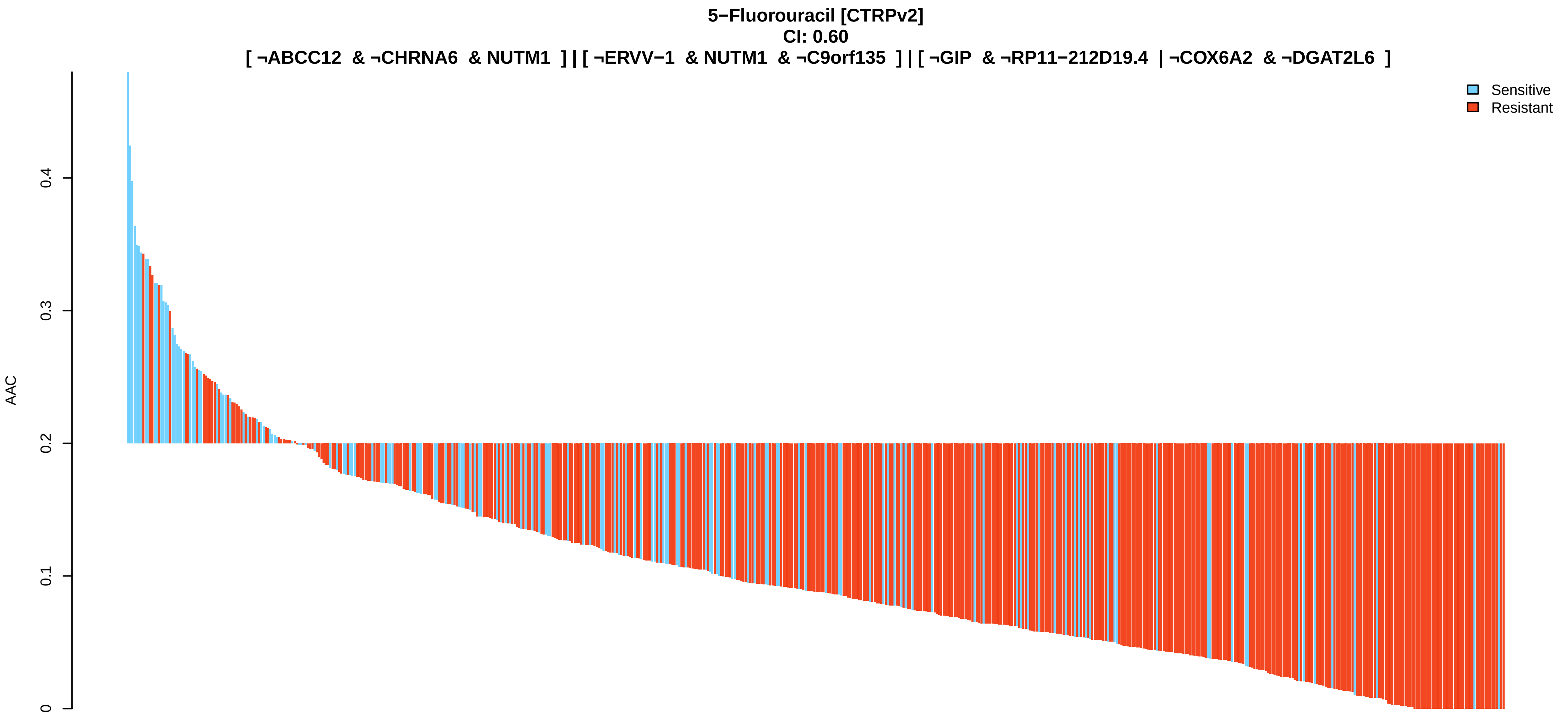
5−Fluorouracil [CTRPv2]
CI: 0.60
[ ¬ABCC12 & ¬CHRNA6 & NUTM1 ] | [ ¬ERVV−1 & NUTM1 & ¬C9orf135 ] | [ ¬GIP & ¬RP11−212D19.4 | ¬COX6A2 & ¬DGAT2L6 ]
Sensitive
Resistant
AAC
0
0.1
0.2
0.3
0.4

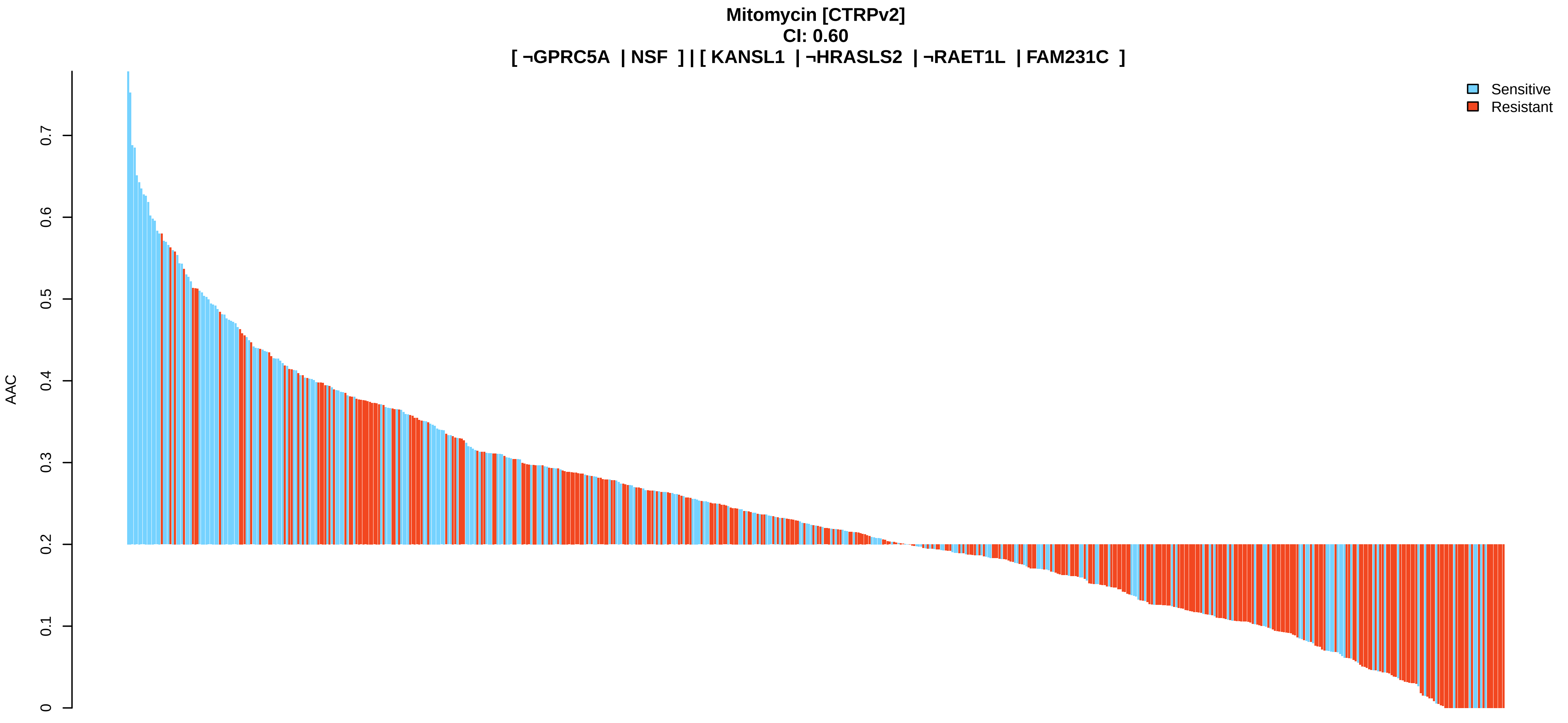
Mitomycin [CTRPv2]
CI: 0.60
[ ¬GPRC5A | NSF ] | [ KANSL1 | ¬HRASLS2 | ¬RAET1L | FAM231C ]
Sensitive
Resistant
AAC
0
0.1
0.2
0.3
0.4
0.5
0.6
0.7

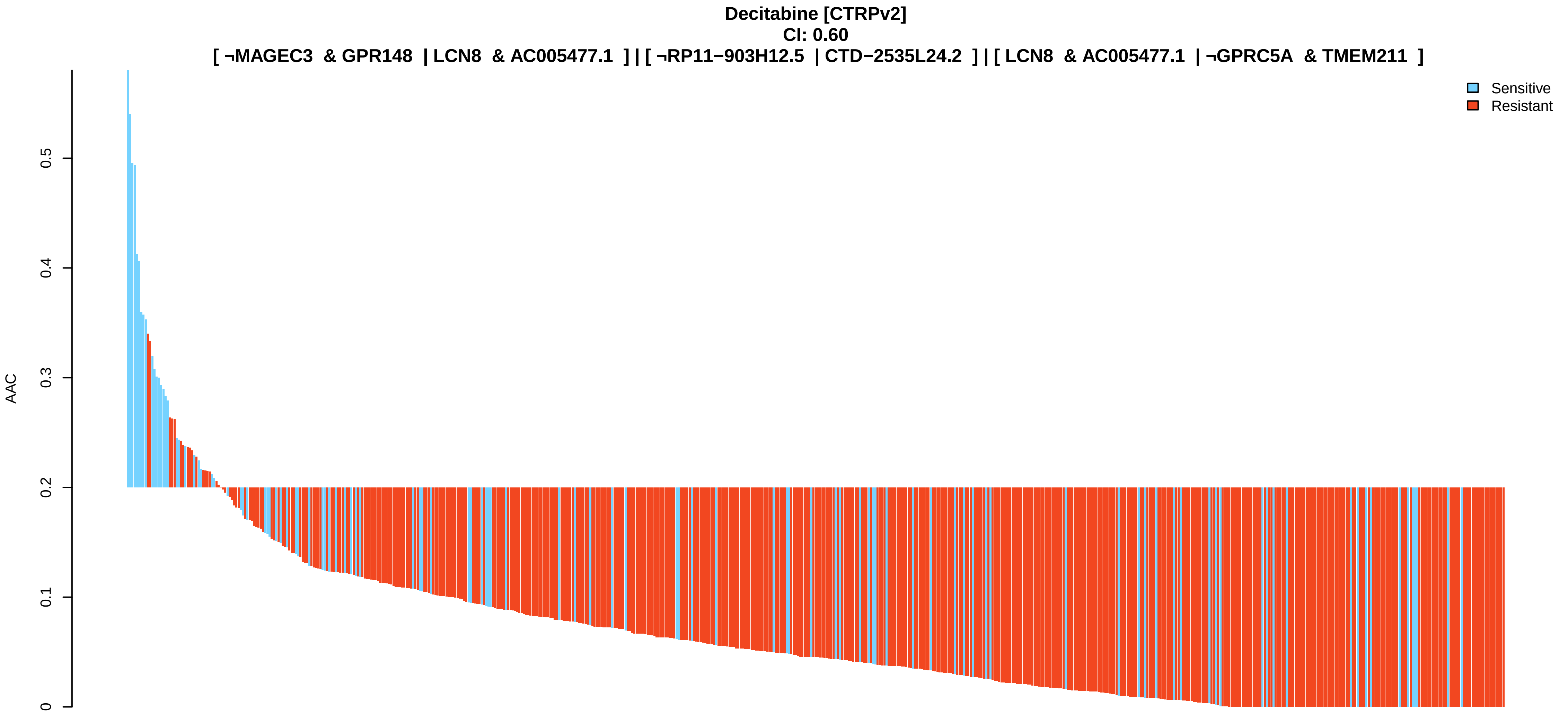
Decitabine [CTRPv2]
CI: 0.60
[ ¬MAGEC3 & GPR148 | LCN8 & AC005477.1 ] | [ ¬RP11−903H12.5 | CTD−2535L24.2 ] | [ LCN8 & AC005477.1 | ¬GPRC5A & TMEM211 ]
Sensitive
Resistant
AAC
0
0.1
0.2
0.3
0.4
0.5

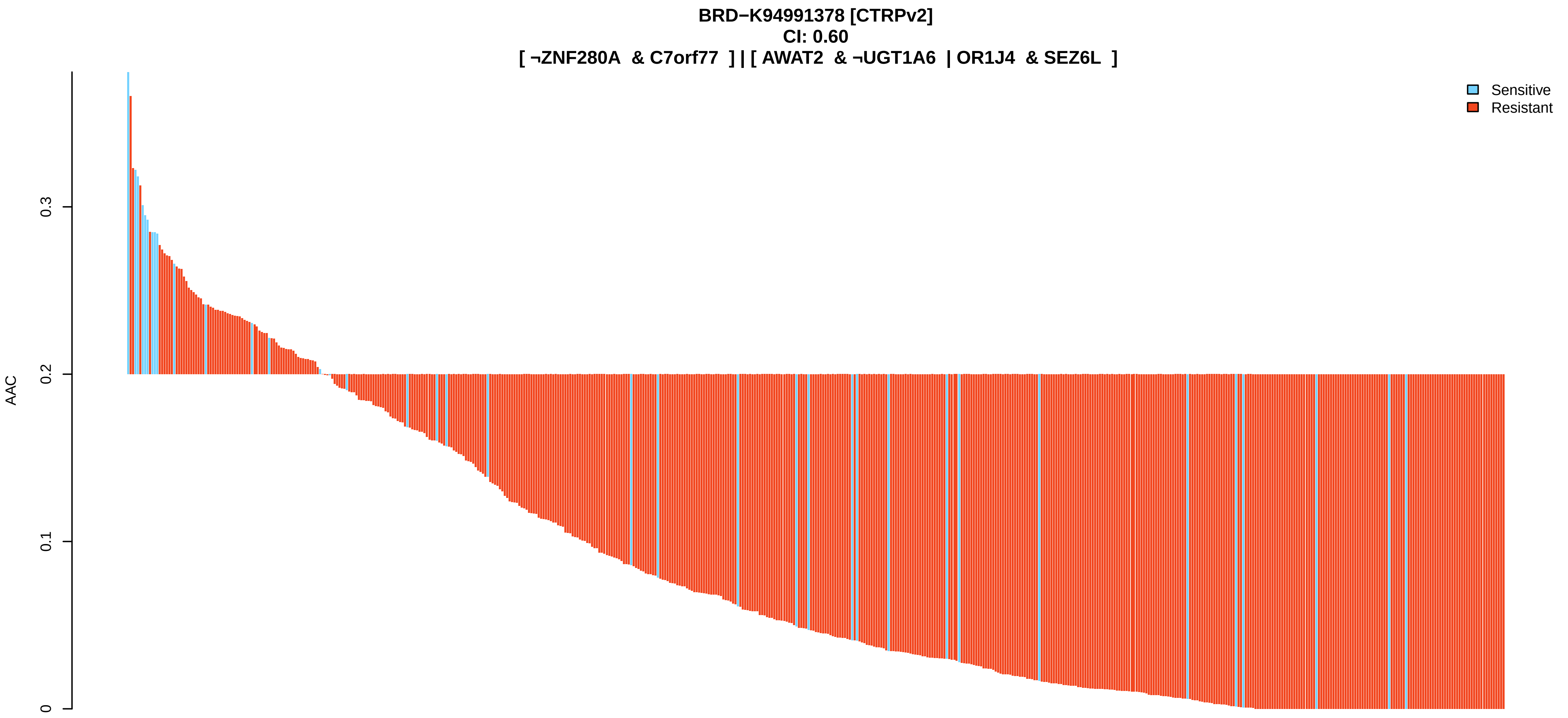
BRD−K94991378 [CTRPv2]
CI: 0.60
[ ¬ZNF280A & C7orf77 ] | [ AWAT2 & ¬UGT1A6 | OR1J4 & SEZ6L ]
Sensitive
Resistant
AAC
0.3
0.2
0.1
0

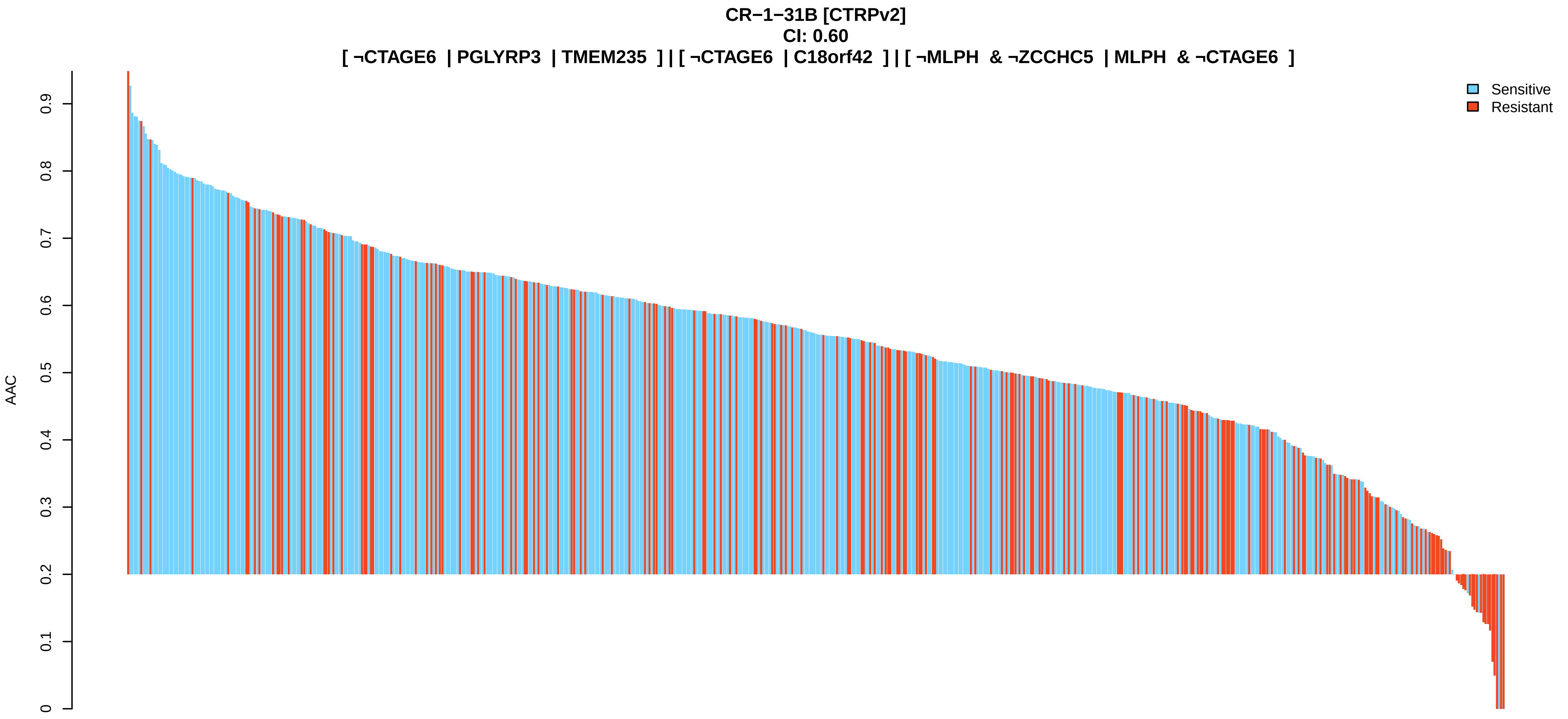
CR−1−31B [CTRPv2]
CI: 0.60
[ ¬CTAGE6 | PGLYRP3 | TMEM235 ] | [ ¬CTAGE6 | C18orf42 ] | [ ¬MLPH & ¬ZCCHC5 | MLPH & ¬CTAGE6 ]
Sensitive
Resistant
AAC
0.9
0.8
0.7
0.6
0.5
0.4
0.3
0.2
0.1
0

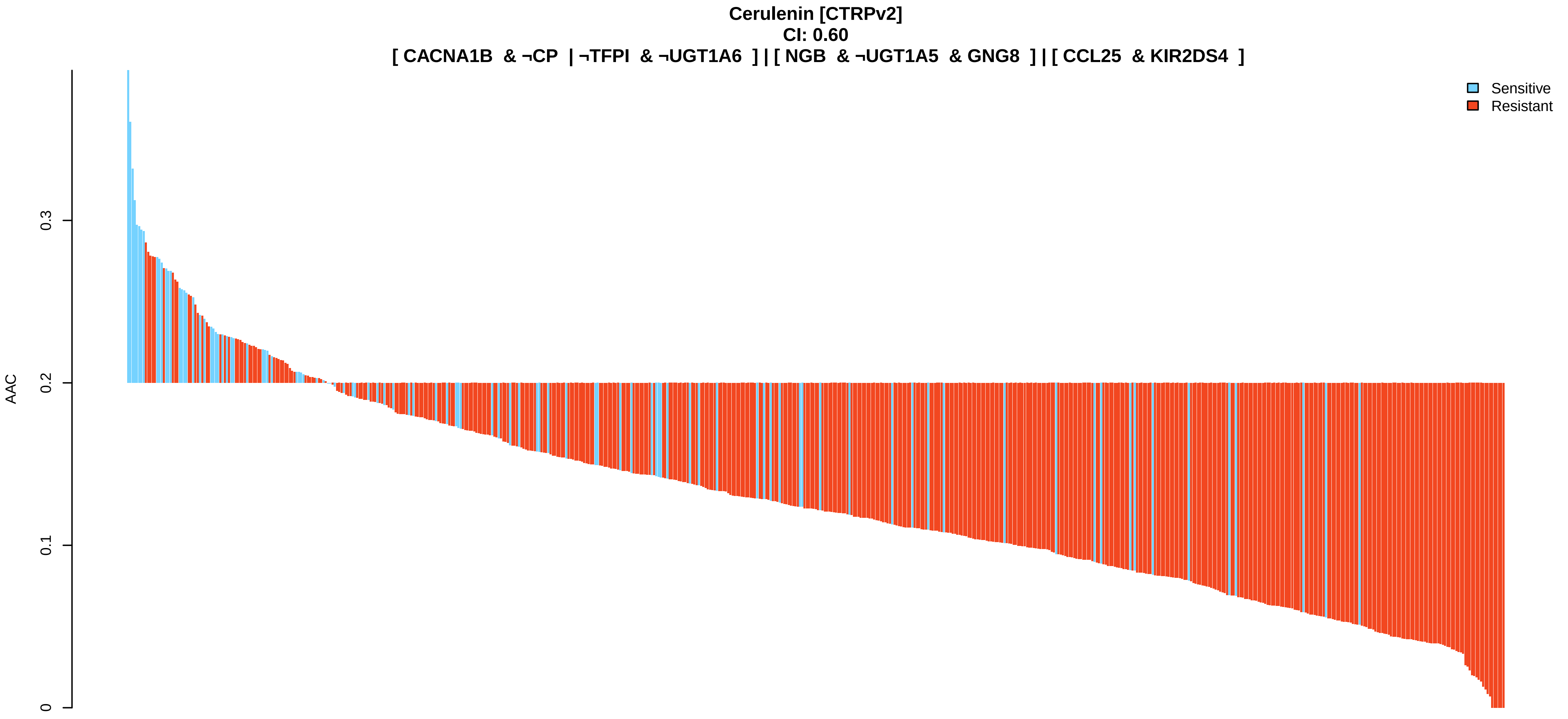
Cerulenin [CTRPv2]
CI: 0.60
[ CACNA1B & ¬CP | ¬TFPI & ¬UGT1A6 ] | [ NGB & ¬UGT1A5 & GNG8 ] | [ CCL25 & KIR2DS4 ]
Sensitive
Resistant
AAC
0.3
0.2
0.1
0

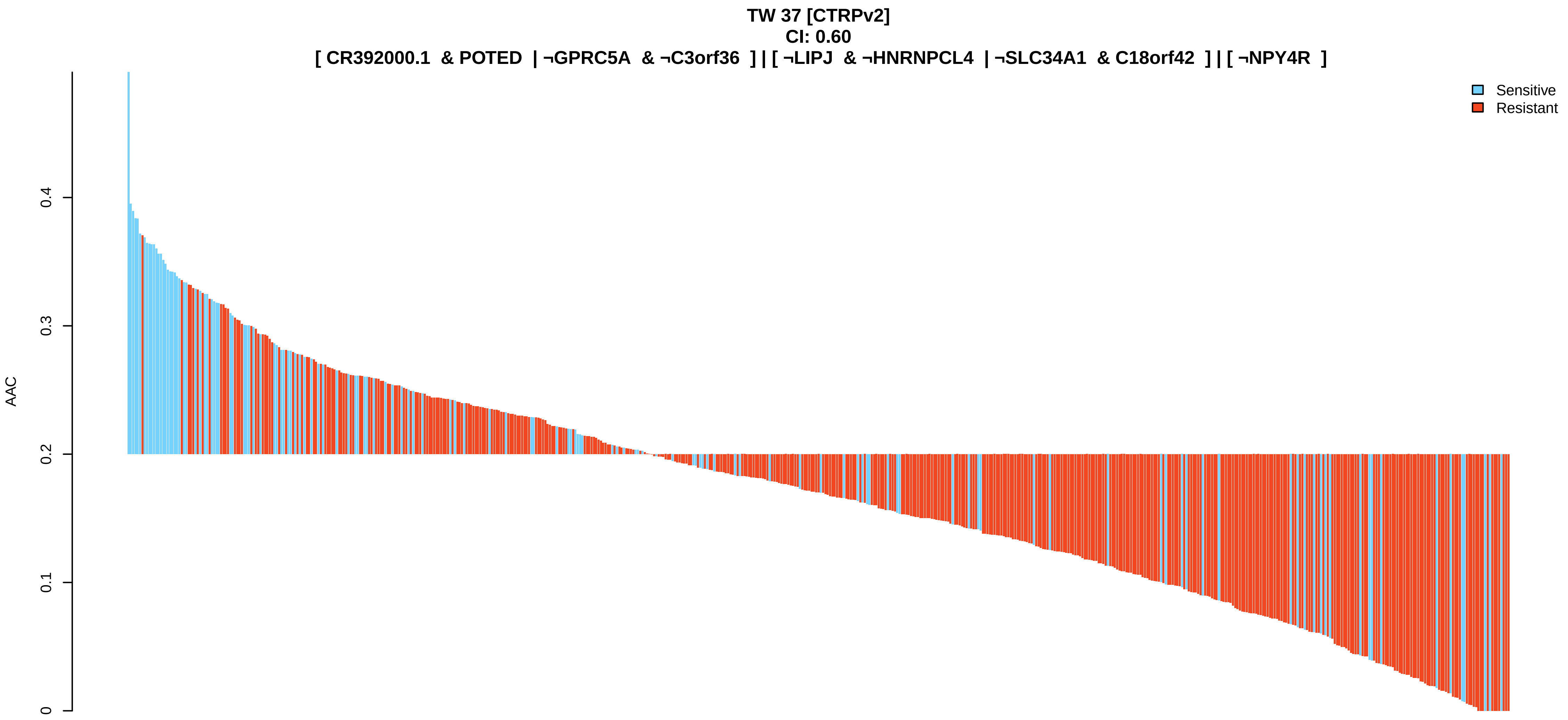
TW 37 [CTRPv2]
CI: 0.60
[ CR392000.1 & POTED | ¬GPRC5A & ¬C3orf36 ] | [ ¬LIPJ & ¬HNRNPCL4 | ¬SLC34A1 & C18orf42 ] | [ ¬NPY4R ]
Sensitive
Resistant
AAC
0
0.1
0.2
0.3
0.4

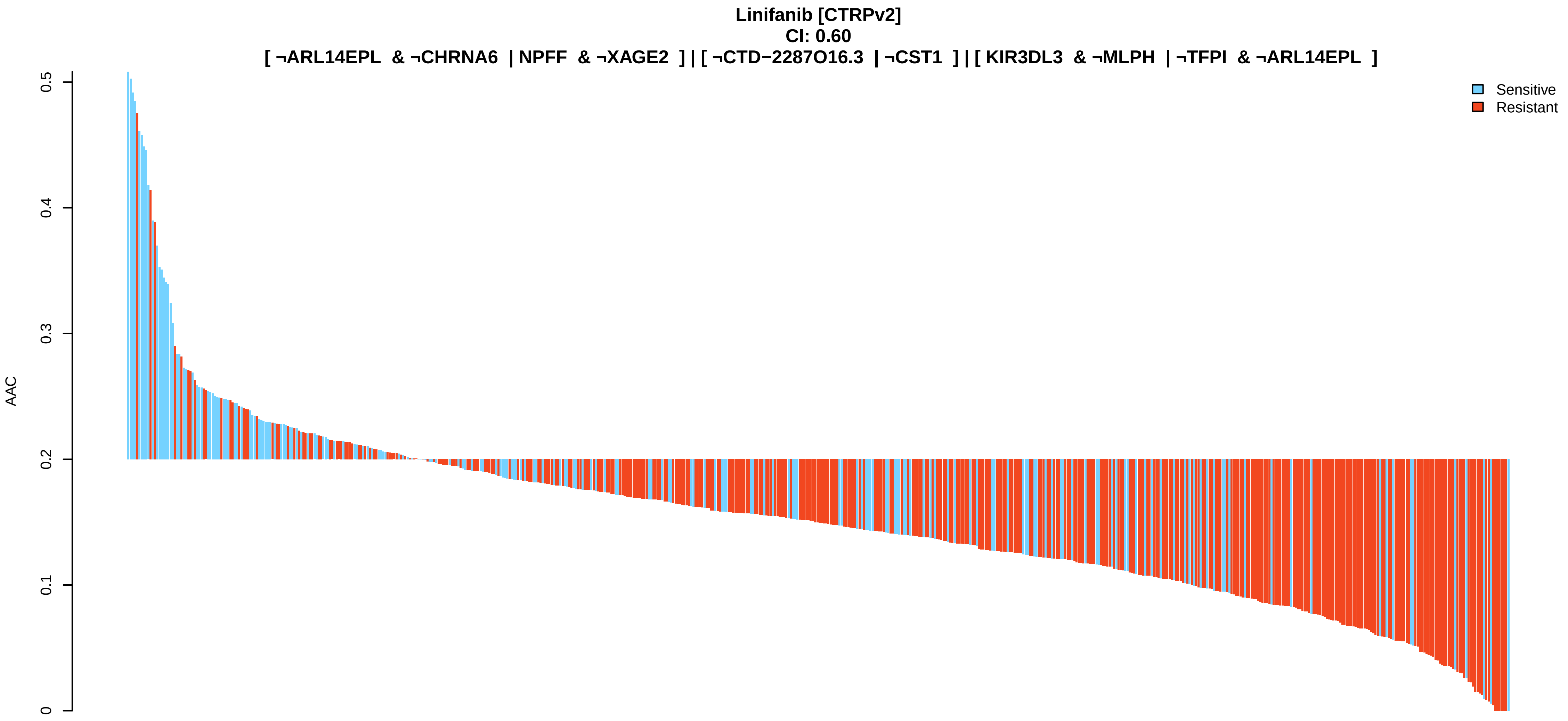
Linifanib [CTRPv2]
CI: 0.60
[ ¬ARL14EPL & ¬CHRNA6 | NPFF & ¬XAGE2 ] | [ ¬CTD−2287O16.3 | ¬CST1 ] | [ KIR3DL3 & ¬MLPH | ¬TFPI & ¬ARL14EPL ]
Sensitive
Resistant
AAC
0
0.1
0.2
0.3
0.4
0.5
